# Tumor- and immune-derived *N*-acetyl-β-D-hexosaminidase drive colorectal cancer and stratify patient risk

**DOI:** 10.1101/2024.07.02.601644

**Authors:** Rebeca Kawahara, Liisa Kautto, Naaz Bansal, Priya Dipta, The Huong Chau, Benoit Liquet-Weiland, Seong Beom Ahn, Morten Thaysen-Andersen

## Abstract

Non-invasive prognostic markers are needed to improve survival of colorectal cancer (CRC) patients. Towards this goal, we applied integrative systems glycobiology approaches to tumor tissues and PBMCs from CRC patients and matching controls as well as to a CRC patient-derived cell line. Firstly, quantitative glycomics and glycoproteomics revealed that non-canonical paucimannosidic proteins from monocytic and cancer cell origins are prominent signatures in CRC tumor tissues, and that their expression associates with CRC progression. Guided by these associations, we then showed that *N*-acetyl-β-D-hexosaminidase (Hex) facilitates paucimannosidic protein biosynthesis in CRC cells and is intimately involved in processes underpinning CRC metastasis (adhesion, migration, invasion, proliferation). Finally, Hex activity was found to be elevated in PBMCs and plasma from patients with advanced CRC relative to matching controls while plasma Hex activity correlated strongly with CRC patient survival. Our study opens avenues for better prognostication, disease risk stratification and therapeutic interventions in CRC.

**Highlights:**

• Non-canonical truncated glycans of immune and cancer origins dominate in CRC tumors
• *N*-acetyl-β-D-hexosaminidase, a truncating glycoenzyme, drives tumorigenesis in CRC
• Activity of plasma *N*-acetyl-β-D-hexosaminidase stratifies risk in CRC patients

## Introduction

Colorectal cancer (CRC) is the second-leading cause of cancer-related deaths worldwide, accounting for approximately 1 million deaths annually^1^. Currently, prognostication and treatment decisions are guided by the tumor-node-metastasis (TNM) staging system, determined subjectively through clinical and histopathological features of tissue biopsies at the time of diagnosis^2^. Surgical removal is preferred for early and locally advanced CRC, while adjuvant chemotherapy is recommended if the CRC has already spread to adjacent lymph nodes.

CRC is a heterogeneous disease with variable outcomes even across cases with the same TNM stage^3^. Most patients diagnosed with locoregional CRC will benefit from surgical resection of primary tumors. However, 30-40% of those patients will develop metastasis in the subsequent years, and the cure rates for this large patient group are dishearteningly low^4^. Although the detection of genetic mutations in KRAS, APC and TP53 in CRC tissues are currently used to predict response to therapy and patient risk (survival chances), the invasive nature and costs associated with such genetic analyses limit their clinical utility^5^. Thus, there is a need for novel non-invasive prognostic markers that can identify CRC patients with high risk of poor outcome, thereby providing opportunities for early intervention and consequently improved patient survival while avoiding unnecessary over-treatment of lower-risk patient groups^6^.

Aberrant asparagine (*N*)-linked glycosylation has repeatedly been linked to malignant transformation and tumor progression in CRC^7–9^ and across other cancers^10–12^, thus offering a considerable and still largely untapped potential for biomarker discovery and therapeutic applications. However, glycoproteins are analytically challenging to study within complex cellular and tissue environments even in specialized biochemical laboratories due to the inherent glycoproteome heterogeneity and their unpredictable structural characteristics arising from non- templated biosynthetic processes^13^. While altered *N*-glycosylation has previously been reported in tumor tissues and sera of CRC patients^14–17^ and was found to hold a potential to inform on drug resistance^18^, prognosis^19^, staging^20^ and recurrence^16^ in CRC, the lack of biomarker specificity and the need for sophisticated glyco-analytical workflows have hitherto precluded the translation of these findings into the clinic.

Aiming to discover robust and clinically-compatible (easy-to-assay) glyco-markers to aid the management and outcomes of CRC patients, in this work, we first applied systems glycobiology approaches to tumor tissues and PBMCs from CRC patients and matching controls as well as a patient-derived CRC cell line to comprehensively characterize aberrations occurring in the glycoproteome during disease progression. Enabled by quantitative glycomics and glycoproteomics, we identified that non-canonical paucimannosidic proteins of both immune and cancer cell origins form prominent glyco-signatures in CRC tumor tissues and that their expression correlates with disease progression. Informed by these relationships, we then showed *in vitro* that *N*-acetyl-β-D-hexosaminidase (Hex) drives paucimannosidic protein formation and, using a pharmacological Hex inhibitor, demonstrated the involvement of Hex in metastatic processes including adhesion, migration, invasion and proliferation in CRC cells. Finally, we used a simple and robust enzyme assay to document that the Hex activity in neat plasma exhibits a potential to stratify CRC patient risk in terms of their five-year survival. Collectively, our findings open new avenues for effective prognostication and therapeutic intervention in patients suffering from CRC.

## Results

### Paucimannosylation is raised in CRC tumors and PBMCs and associates with disease progression

Aiming to discover new glyco-markers for CRC, this study firstly employed a multi-faceted -omics approach to study various biospecimens (fresh and FFPE tissues, PBMCs, plasma) from CRC patients and matching controls as well as a patient-derived CRC cell line (see **Supplementary Table S1** for an overview of samples and experimental approach). Systems glycobiology, namely integrated glycomics, glycoproteomics and proteomics paired with bespoke informatics workflows, was used to globally quantify *N*-glycan structures, their sites and protein carriers, as well as provide clues to their cellular origins from tissues and PBMCs of 28 CRC patients (n = 7/stage) (**Figure 1A-B**). Adjacent normal tissues from CRC patients in disease stage I (n = 8) and PBMCs from healthy donors (n = 8) were used as controls.

**Figure 1.**
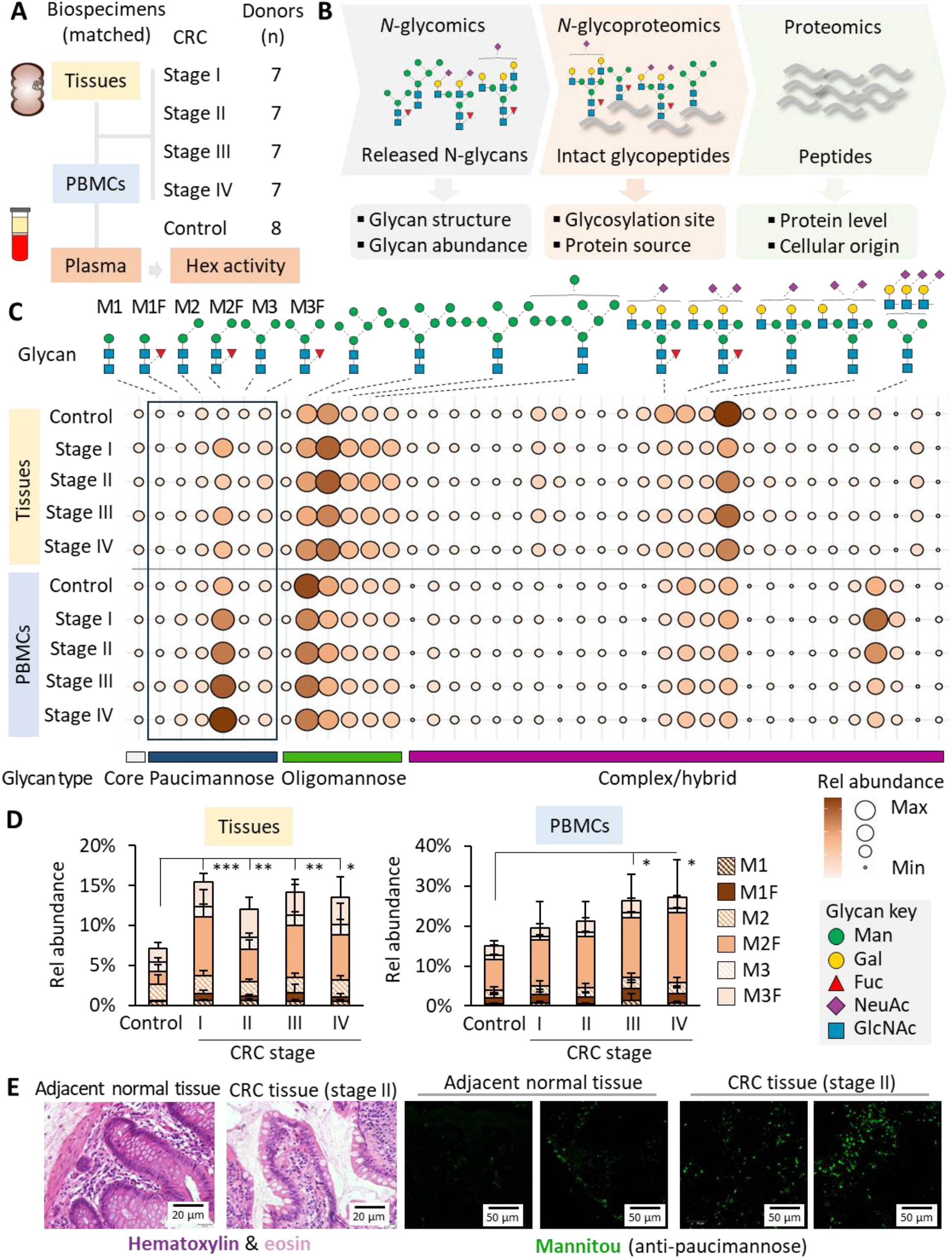
Strong paucimannosidic glyco-signatures in CRC tumor tissues and PBMCs. **A)** Sample cohort used for the discovery phase of the study including tumor tissues, PBMCs and plasma from 28 patients with CRC (stage I-IV). Adjacent normal tissues from 8 CRC patients (stage I) and paired PBMCs and plasma from 8 non-CRC (normal) donors were used as controls (see **Supplementary Table S1** for details). **B**) Systems glycobiology workflows used to map the expression and cellular origins of glycoproteins across the CRC stages (relative to controls) in tumor tissues and PBMCs. **C**) Comparative glycomics of matched tumor tissues and PBMCs across the CRC stages compared to controls. Boxed: Expression of paucimannosidic *N*-glycans (M1-M3F) (**Supplementary Table S2-S3**). **D**) Stage-specific expression of protein paucimannosylation in CRC tumor tissues and PBMCs relative to controls as measured by glycomics (n = 7/CRC stage, n = 8 control, student’s T-test, \**p* < 0.05, \*\**p* < 0.01, \*\*\**p* < 0.001). **E**) Representative examples of tissue architecture and paucimannose expression as determined with H&E (left) and staining with a paucimannose-reactive antibody (Mannitou, right), respectively, in matched CRC tumor tissues and normal adjacent tissues from a CRC patient (stage II).

Unlike the conventional *N*-glycan types (oligomannosidic-, hybrid-, complex-type), the less investigated non-canonical paucimannosidic-type glycans i.e. Man1-3GlcNAc2Fuc0-1 (short-hand M1-M3F)^21–23^ were strongly elevated in the *N*-glycome of CRC tumor tissues (CRC stage I-IV: 13.5-15.4%, adjacent normal tissues: 6.7%) and in PBMCs (CRC stage I-IV: 19.5-27.1%, healthy donors: 14.9%) as determined using quantitative glycomics (**Figure 1C** and **Supplementary Table S2-S3** and **Supplementary Data S1**). Significant elevation in protein paucimannosylation in the CRC tumor tissues was indeed observed across all disease stages compared to adjacent normal tissues and in PBMCs from CRC patients with advanced disease (stage III-IV) (**Figure 1D**). The core fucosylated M2F and M3F structures (Man2-3GlcNAc2Fuc1) dominated in both CRC tumor tissues and PBMCs covering 60-70% of the paucimannosidic glycans. Recapitulating findings from the glycomics experiments, an increased staining response to Mannitou, a paucimannose-reactive antibody^24,25^, was apparent for tumor tissues from CRC patients (stage II) compared to adjacent normal tissue sections from the same patients (**Figure 1E**).

### Paucimannosidic proteins from immune and cancer cells dominate in CRC tumor tissues

We then used quantitative glycoproteomics to map the relative expression of paucimannosidic proteins across CRC tumor tissues and PBMCs from patients and controls as well as a CRC cell line (LIM2405) originating from a patient with poorly differentiated adenocarcinoma of cecum (**Figure 2A**, see **Supplementary Tables S4-S6** for data). The PBMCs and CRC cell line were included to test for potential immune and cancer cell sources of the paucimannosidic proteins observed in the CRC tumor microenvironment (TME). Indeed, of the 51 paucimannosidic proteins identified in the CRC tumor tissues, 7 (13.7%) and 13 (25.5%) proteins were also identified to carry paucimannosidic *N*-glycans in the PBMCs and CRC cell line, respectively. Cellular component analysis showed that the paucimannosidic proteins identified in the CRC tumor tissues were mainly from lysosomal and extracellular regions (**Figure 2B**) and spanned both soluble and membrane-tethered proteins (**Figure 2C**).

**Figure 2.**
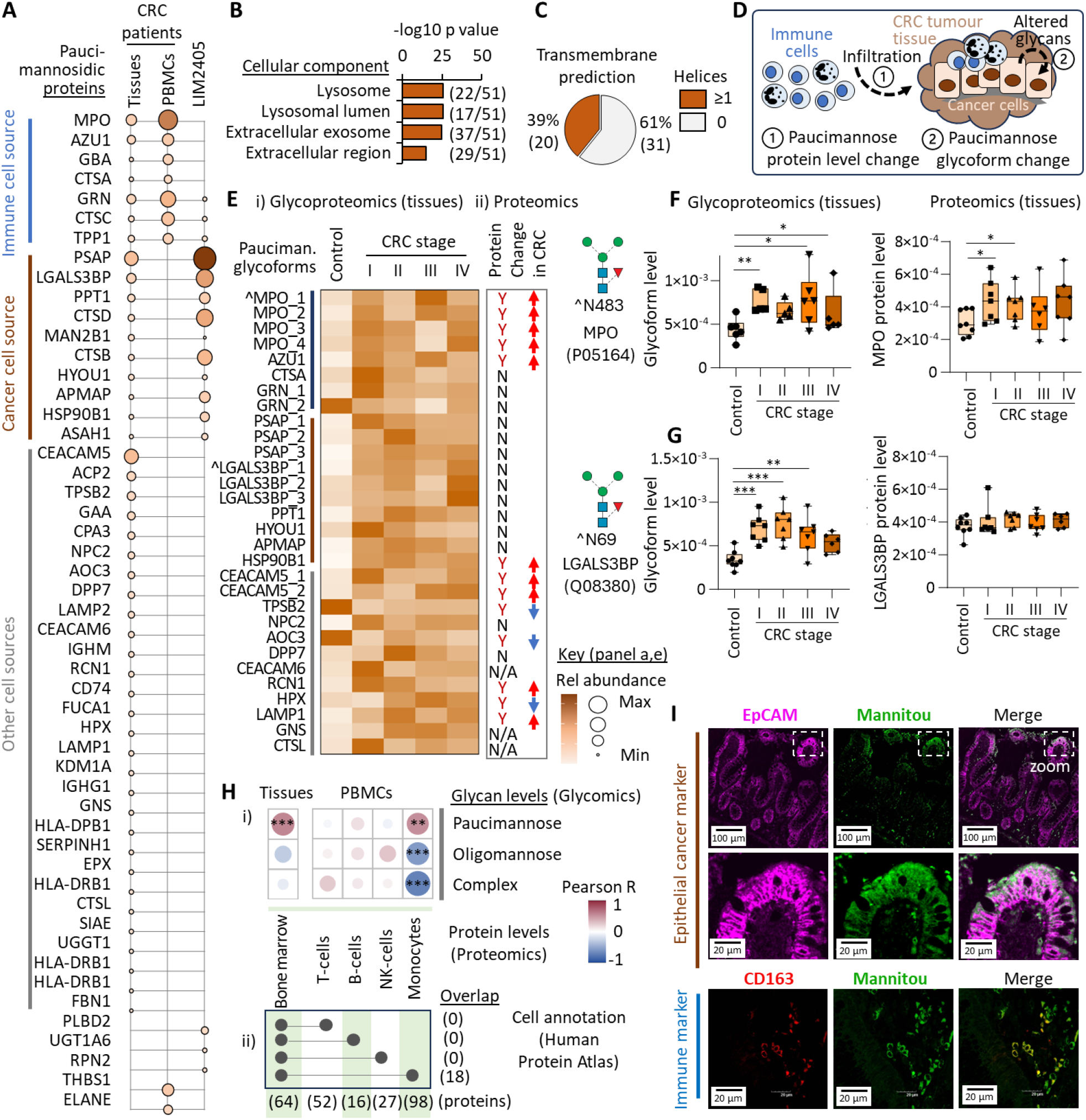
Paucimannosidic proteins of monocytic and cancer cell origins are prominent features in CRC tumor tissues. **A)** Distribution of paucimannosidic glycopeptides from 51 proteins identified in CRC tumor tissues, PBMCs and in a CRC patient-derived cell line (LIM2405) using glycoproteomics (see **Supplementary Table S4-S6** for data and panel E for key). Prediction of **B)** cellular localization (Gene Ontology) and **C)** transmembrane regions (TMHMM 2.0) of paucimannosidic proteins identified in CRC tumor tissues, PBMCs and LIM2405 cells. **D)** Schematics of two mechanisms (labelled 1 and 2) hypothesized to contribute to the paucimannosidic signatures in CRC tumor tissues. **E)** Changes in (i) site-specific paucimannosidic glycoform distribution as measured by glycoproteomics and (ii) protein levels (proteomics) in CRC tumor tissues relative to adjacent normal tissues (see **Supplementary Table S4** and **Supplementary Table S7** for data). Each row represents a unique protein glycoform (unique glycosylation site and paucimannosidic structure). Only paucimannosidic glycoforms showing significant changes are listed (n = 7/stage, n = 8 controls, T-tests, *p* < 0.05, N/A = protein not detected in proteomics dataset). Arrows: Direction of change in CRC tumors vs adjacent normal tissues. Representative examples of changes in the site-specific paucimannosidic glycoform distribution and/or protein levels in CRC tumors vs adjacent normal tissues for **F)** MPO (immune origin) and **G)** LGALS3BP (cancer cell origin). **H)** i) Correlation analysis of cell-specific protein markers (proteomics) and glycosylation type (glycomics) in CRC tumor tissues and PBMCs. Pearson correlation (n = 36), \**p* < 0.05, \*\**p* < 0.01, \*\*\**p* < 0.001. ii) Proteins identified in the CRC tumor tissues and PBMCs were annotated for cellular origin using the Human Protein Atlas and their overlap plotted. **I)** IHC analysis targeting paucimannosidic glycoepitopes (Mannitou, Alexa Fluor 488, green), epithelial cancer cells (anti-EpCAM antibodies, Alexa Fluor 647, magenta) and anti-inflammatory macrophages arising from tumor-infiltrating monocytes (anti-CD163 antibodies, Alexa Fluor 405, red) in CRC tumor tissues (stage II). Co-localization (yellow) was visually assessed (merge).

Building on our recent reports that circulating innate immune cells (e.g. neutrophils, monocytes) constitutively express and store paucimannosidic proteins that may be released upon activation^26–28^ and the established knowledge that innate immune cells abundantly infiltrate the CRC TME^29,30^, these observations led us to hypothesize that the strong paucimannosidic signatures in the tumor tissues may, in part, originate from infiltrating immune cells and, in part, arise from aberrations in the glycosylation machinery of the growing epithelial cancer cells (**Figure 2D**).

To support our hypothesis, we performed an integrative analysis combining our quantitative glycoproteomics data (measuring protein glycoform levels) with quantitative proteomics data (protein level measurements), which revealed that for key innate immune-derived proteins (e.g. MPO, AZU1), the elevation in paucimannosidic glycoforms were accompanied by an elevation in their respective protein levels in CRC tumors relative to adjacent normal tissues (**Figure 2E**, **Supplementary Table S7-S8**). Site-specific glycoprofiling of MPO (Asn483) substantiated this relationship (**Figure 2F**). In contrast, for proteins annotated to putatively arise from a CRC cell origin (e.g. PSAP, LGALS3BP), the elevation in paucimannosidic glycoforms were generally not accompanied by changes in their protein levels, as exemplified by the site-specific glycoprofiling of LGALS3BP (Asn69) (**Figure 2G**).

To further explore the immune cell origin(s) of the paucimannosidic signatures in the CRC TME, proteins identified in the CRC tumor tissues were annotated using the Human Protein Atlas (tissue section) and those of immune (bone marrow) origin were correlated with glycan features observed across the paired proteomics-glycomics datasets (**Figure 2Hi**, see also **Supplementary Table S2** and **Supplementary Table S7** for data). The expression of bone marrow-derived immune proteins displayed a strong association with the relative levels of protein paucimannosylation (Pearson correlation, R = 0.57, *p* = 0.0003 n = 36) and not with oligomannosidic- and complex-type glycosylation. A similar approach was used for proteins identified in the PBMC sample in this case employing the blood atlas within the Human Protein Atlas to annotate proteins of their immune cell origin (T-, B- and NK-cells and monocytes). Associations (Pearson R = 0.52, p = 0.001, n = 36) were observed between the paucimannose levels (glycomics) and the expression of monocyte- derived proteins (proteomics), not towards proteins annotated to be of lymphocytic origin (T-, B- and NK-cells) (see **Supplementary Table S3** and **Supplementary Table S8** for data). Notably, only proteins of monocytic origin (not from T-, B- and NK-cells) were found amongst the bone marrow-annotated proteins identified in the CRC tumor tissues, indicating that the paucimannosidic signatures in the CRC TME, at least in part, originate from tumor-infiltrating monocytes (**Figure 2Hii**).

To visually document the cellular origin(s) of the paucimannosidic features in CRC tumor tissues, immunohistochemistry (IHC) analysis was performed employing Mannitou to map the spatial expression of paucimannosidic glycoepitopes relative to either anti-inflammatory (CD163+) macrophages arising from infiltrating monocytes or epithelial (EpCAM+) cancer cells within CRC tumor tissue sections from a patient with stage II disease (**Figure 2I**). While little direct co- localization between paucimannosylation and EpCAM was observed, considerable Mannitou staining was observed immediately outside the epithelial cancer cells possibly arising from lysosomal exocytosis of soluble paucimannosidic proteins (Figure 2B-C). Importantly, paucimannosidic glycoepitopes were also found to co-localize directly with the anti-inflammatory macrophages in the CRC tumor tissues. These observations therefore support that both infiltrating innate immune cells and epithelial cancer cells contribute to the strong paucimannosidic signatures present in CRC tumor tissues.

### Hex facilitates paucimannose biosynthesis in CRC cells and drives key tumorigenic processes

Using genetic knockouts of *HEXA* and *HEXB* encoding the α- and β-subunits of *N*-acetyl-β-D- hexosaminidase (Hex), respectively, we have previously shown in human myeloid cells that the hetero- and homo-dimeric Hex isoenzymes i.e. Hex A (αβ) and Hex B (ββ) catalyze the formation of paucimannosidic proteins via an unconventional truncation pathway^31^ (**Figure 3A**). As it remains unexplored if Hex also drives the biosynthesis and biological functions of paucimannosidic proteins in CRC cells, we used systems glycobiology approaches and a known pharmacological Hex inhibitor (M31850)^32^ to comprehensively test these putative relationships in a patient-derived CRC cell line (LIM2405) (**Figure 3B**).

**Figure 3.**
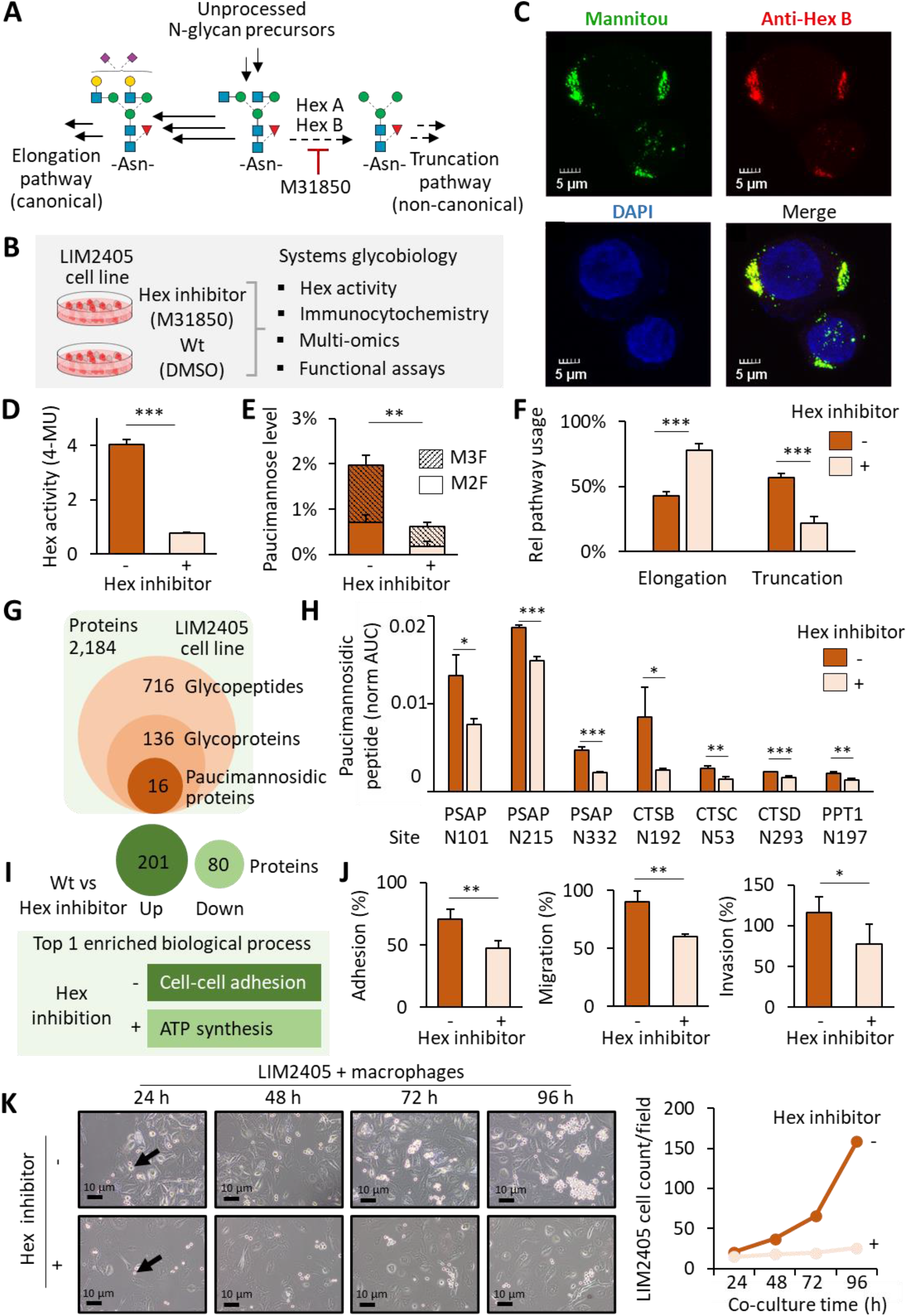
Hex-mediated biosynthesis of paucimannose and pro-tumor roles in CRC. **A)** In human myeloid cells, a non-canonical *N*-acetyl-β-D-hexosaminidase (Hex)-driven truncation pathway co-exists alongside the canonical elongation pathway responsible for *N*-glycoprotein biosynthesis^31^. M31850 is a known pharmacological Hex inhibitor^32^. **B)** The effect of Hex inhibition by M31850 was studied in a patient-derived CRC cell line (LIM2405) using systems glycobiology approaches. **C)** Hex B co-localizes with paucimannosidic glycoepitopes in LIM2405 cells as shown by immunocytochemistry using paucimannose-reactive Mannitou (green) and anti- Hex B (red) antibodies. DAPI stain in blue. **D)** Total Hex activity as measured with a Hex activity assay using a MUG substrate (effectively a proxy for Hex B (ββ) activity given insignificant levels of Hex subunit α). **E)** Levels of fucosylated paucimannose (M2F, M3F) and **F)** relative usage of the truncation pathway as determined by glycomics. For panel D-F: Hex inhibited LIM2405 cells were compared to untreated (DMSO) controls using t-tests (n = 3/experiment, \*\**p* < 0.01, \*\*\**p* < 0.001, see **Supplementary Table S9** for data). **G)** Overview of (glyco)proteins identified from proteomics (green) and glycoproteomics (orange) data of LIM2405 cells (see **Supplementary Table S6** and **Supplementary Table S10** for data). **H)** Site-specific reduction of paucimannosylation across a subset of the identified paucimannosidic proteins in Hex inhibited compared to untreated LIM2405 cells as measured by glycoproteomics (n = 3/experiment, \**p* < 0.05, \*\**p* < 0.01, \*\*\**p* < 0.001, see **Supplementary Table S10** for data). **I)** Proteome expression changes (*p* < 0.05) in LIM2405 cells upon Hex inhibition and the involvement of altered proteins in biological pathways. **J)** Hex inhibition leads to a dose-dependent reduction in cell adhesion (n = 3, t-test, \*\**p* < 0.01), migration (n = 4/experiment, t-test, \*\**p* < 0.01) and invasion (n = 5/experiment, t-test, \**p* < 0.05) in LIM2405 cells, see **Supplementary Figure S2D** for details. **K)** LIM2405 cells were pre-treated for 24 h with or without Hex inhibitor and co-cultured with anti-inflammatory macrophages in inhibitor-free media. Arrows: Examples of LIM2405 cells. Longitudinal plot of LIM2405 cell count. Images: 10x magnification.

The Hex β-subunit (rather than the α-subunit) was found to consistently dominate in CRC tumor tissues and PBMCs as determined by proteomics (**Supplementary** Figure 1A-B). Supporting the involvement of the Hex B isoenzyme (ββ) in the formation of paucimannosidic proteins in CRC cells, Hex B was found to accurately co-localize with paucimannosidic glycoepitopes in LIM2405 cells as assessed by immunocytochemistry (**Figure 3C**).

To enable a series of functional experiments with M31850, we first tested across an extended concentration range the impact of the Hex inhibitor and its vehicle i.e. DMSO on LIM2405 cells (**Supplementary** Figure 2A). At 31.25 µM M31850, the Hex activity was effectively inhibited without compromising the cell viability caused by higher vehicle concentrations (**Figure 3D** and **Supplementary** Figure 2B). Glycomics profiling showed a M31850 dose-dependent decrease in the levels of paucimannosylation relative to the vehicle control documenting that Hex, as expected, catalyze paucimannosidic protein formation in CRC cells (**Supplementary** Figure 2C). At 31.25 µM M31850, the LIM2405 cells displayed a significant reduction in protein paucimannosylation and a lower utilization of the truncation pathway relative to the DMSO controls (**Figure 3E-F** and **Supplementary Table S9**). These Hex inhibition conditions (31.25 µM M31850, 24 h) were therefore chosen for the downstream functional experiments.

Next, we performed glycoproteomics and proteomics of LIM2405 cells with and without M31850 treatment to map quantitative changes to the glycoproteome in this CRC model system upon Hex inhibition. In total, 16 paucimannosidic proteins were identified, covering 31 glycosylation sites and several different paucimannosidic glycan structures (**Figure 3G**, **Supplementary Table S6** and **Supplementary Table S10**). In line with observations made for the CRC tumor tissues (Figure 2B), the paucimannosidic proteins were predominantly of lysosomal origin (KEGG pathway, Benjamini FDR = 2.0^-9^, data not shown), including ASAH1, CTSB, CTSC, CTSD, MAN2B1, PPT1, PSAP and TPP1, five of which showed a significant site-specific reduction of paucimannose upon Hex inhibition (**Figure 3H**).

Quantitative proteomics showed that Hex inhibition also altered the expression of the proteome (**Figure 3I**). Specifically, 201 proteins (out of 2,184, 9.2%) were under-expressed, and 80 proteins (3.7%) were over-expressed upon Hex inhibition. Notably, proteins that were reduced upon Hex inhibition were significantly associated with cell-cell adhesion processes (23/183 proteins, 11.5% e.g. testin and cingulin, FDR = 3.1 x 10^-10^) (Supplementary Table S10).

We then explored the effect of Hex inhibition in the LIM2405 model system with respect to cellular processes associated with cancer progression and metastasis using established functional assays and co-culture experiments. Interestingly, the Hex inhibited LIM2405 cells showed a reduced ability to adhere to extracellular matrix (*p* = 0.004, n = 3/experiment), to migrate (*p* = 0.003, n = 4/experiment) and to invade matrix (*p* = 0.027, n = 5/experiment), supporting the functional involvement of Hex and/or the paucimannosidic protein products in key CRC processes (**Figure 3J**). Supporting further these important functional relationships, the LIM2405 cells exhibited a dose-dependent reduction of the tumorigenic traits upon Hex inhibition (**Supplementary Figure S2D**). Notably, impaired proliferation was observed in Hex inhibited LIM2405 cells compared to untreated controls when co-cultured for four days with primary monocyte-derived anti-inflammatory macrophages (**Figure 3K**) while similar expansion rates were observed for LIM2405 cells cultured with and without Hex inhibition in media only (**Supplementary Figure S3**). These results suggest that Hex contributes to CRC cell proliferation in micro-environments containing anti-inflammatory macrophages.

### Plasma Hex activity stratifies patient risk

Given the pro-tumorigenic functional roles of Hex, we sought to investigate the prognostic value of using Hex activity in plasma to inform on CRC patient risk in terms of their five-year survival chances. As routinely performed to diagnose Hex-deficiency in Tay-Sachs (*HEXA*^-/-^) and Sandhoff (*HEXB*^-/-^) disease^33^, Hex activity can be readily measured in blood plasma using fluorescent substrates i.e. 4-methylumbelliferyl-2-acetamido-2-deoxy-β-D-glucopyranoside (MUG, measuring total Hex activity) and 6-sulfated MUG (MUGS, measuring Hex A-specific activity). Consistent with the observation that the Hex β subunit dominates in CRC tumor tissues and PBMCs (**Supplementary** Figure 1A-B), the activity of Hex B (ββ) (measured as MUG-MUGS) was found to be the principal Hex isoenzyme form in CRC plasma (**Supplementary** Figure 1C). Given the insignificant activity of the Hex A (αβ) isoform, the downstream experiments were limited to measuring total Hex activity (MUG) in effect serving as a proxy for the Hex B-specific activity.

High total Hex activity was observed in PBMCs and plasma from CRC patients with late-stage disease (stage III-IV) compared to controls, supporting the pro-tumorigenic roles of Hex in CRC progression (**Figure 4A-B**). Notably, correlations (Pearson R = 0.49, *p* = 0.0028) were observed between the total Hex activity in plasma and PBMCs across the 35 matched sample sets, indicating that circulating plasma Hex may, at least in part, arise from PBMCs (**Figure 4C**).

**Figure 4.**
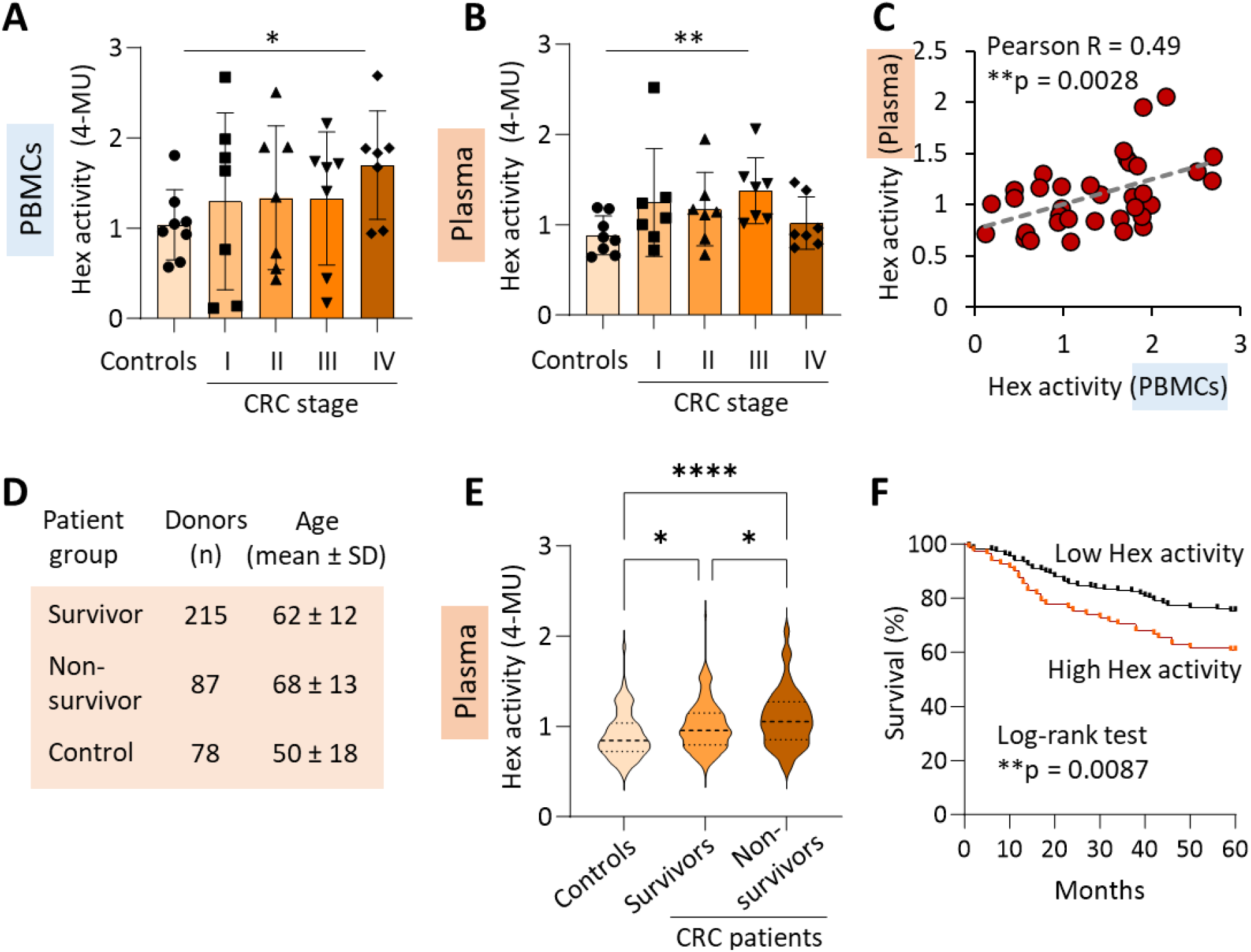
Plasma Hex activity is associated with CRC mortality enabling patient risk stratification. Stage-specific elevation of total Hex activity in matched **A**) PBMCs and **B**) plasma samples from CRC patients relative to controls as measured using a fluorescent-based substrate (MUG) assay. **C)** Correlation between total Hex activity in paired PBMCs and plasma matched from the same individuals (Pearson, 95% confidence interval, n = 35. One outlier was identified in plasma using the ROUT method and was removed). **D)** Plasma cohort used for total Hex activity screening of CRC survivors and non-survivors and control donors. See **Supplementary Table S11** for details. **E**) High total Hex activity in CRC non-survivors (n = 87, mortality within five years post-surgery) compared to CRC survivors (n = 215) and controls (n = 78) as assessed using ANOVA followed by Tukey multiple comparison (**** adjusted *p* < 0.0001, *adj *p* < 0.05). **F**) CRC patients with high Hex activity in plasma (red trace) exhibited lower five-year survival relative to CRC patients with lower plasma Hex activity (black trace) (log-rank test, *p* = 0.0087). Boundary for high/low Hex activity was determined by 75^th^ percentile (high, n = 81, low, n = 221).

To demonstrate clinical compatibility, we then performed a series of benchmarking tests to document that the employed Hex activity assay provides reproducible, sensitive, and robust measurements of the Hex activity using only 2 µl neat plasma per donor. Importantly, the plasma Hex activity appeared both biologically stable as measured across repeat drawings from the same donor (**Supplementary Figure S4A**) and technically stable as shown across multiple freeze/thaw cycles and storage conditions (**Supplementary Figure S4B-C**). Importantly, the Hex activity assay showed robustness and relatively low variability across different blood sample types, including serum, EDTA- or heparin-plasma (**Supplementary Figure S4D**). Collectively, these favorable characteristics indicate that the Hex activity assay is compatible with use in a clinical setting.

To investigate the potential of the plasma Hex activity to stratify patient risk in terms of mortality, total Hex activity was measured in plasma from a cohort of 302 CRC patients for which we had five-year survival data and of another 78 healthy donors (**Figure 4D**). Patients were grouped by patients who survived (n = 215) and those who did not survive (n = 87) five years post- surgery/treatment. Notably, the Hex activity was significantly higher in plasma from non-surviving CRC patients compared to plasma from both the CRC survivors (adjusted *p* = 0.0372) and heathy donor controls (adjusted *p* < 0.0001) (**Figure 4E**). Accordingly, we found that CRC patients featuring high total Hex activity in plasma (1.39 ± 0.21 relative Hex activity units, n = 81) had significant higher five-year mortality risk compared to patients displaying low total plasma Hex activity (0.87 ± 0.16 relative Hex activity units, n = 221, log-rank test, *p* = 0.0087) (**Figure 4F**). Intuitively, mortality is associated with age. Thus, we performed a hazard ratio analysis using a Cox-model to determine the prognostic value of total Hex activity when considering age as a confounding factor (**Table 1**). When adjusted for age, the total Hex activity was still significantly associated with mortality; CRC patients had 2.04 times higher risk of mortality over five years per unit of plasma Hex activity.

**Table 1.** Hazard ratio (HR) to assess impact of age and Hex activity on five-year CRC patient survival.

|  | Mean (SD) | HR (univariable) | HR (multivariable) |
| --- | --- | --- | --- |
| Age (years) | 63.9 (12.7) | 1.03 (1.01-1.05) $p = 0.001$ | 1.03 (1.01-1.05)<br>$p = 0.004$ |
| Hex activity (rel. activity units) | 1.0 (0.3) | 2.49 (1.27-4.86) $p = 0.008$ | 2.04 (1.01-4.11)<br>$p = 0.046$ |

## Discussion

Colorectal cancer (CRC) remains a life-threatening disease featuring high morbidity and mortality across the world^1^. With the goal of identifying new easy-to-assay molecular markers for CRC to inform the clinical decision-making and improve patient outcomes, we have here applied innovative systems glycobiology approaches to valuable collections of CRC and donor biospecimens revealing that non-canonical paucimannosidic proteins and their underpinning biosynthetic glyco-enzyme, *N*-acetyl-β-D-hexosaminidase (Hex), functionally link to CRC progression and patient survival.

While protein paucimannosylation is a constitutively expressed and tissue-wide *N*-glycan modification in plants and lower vertebrates and until recently considered absent in mammals^34,35^, our developing knowledge of paucimannose in the context of human glycobiology points to a tissue-restricted and context-dependent expression of this non-canonical class of *N*-glycoproteins in humans^23^ with reports mainly related to immunity^26,28,36^ and cancer^37–41^ including several in CRC^14,42–44^. These recent observations of non-canonical paucimannosylation in cancer biospecimens including those reported in this study were critically enabled by technology improvements in the emerging field of systems glycobiology including the advent of powerful quantitative glycomics and glycoproteomics^45^. These -omics methods are namely uniquely able to comprehensively survey the glycoproteome at scale even in complex tissues and cellular environments and accurately report on quantitative changes of glycosylation features including unusual (unexpected) structures previously overlooked with traditional methods. The fact that paucimannosylation in this study was found to be a prominent signature in both CRC tumor tissues and PBMCs and was amongst the glycan types displaying the greatest elevation across the CRC stages relative to matched controls made this initial observation an interesting lead to follow.

Identifying the cell source(s) of proteins let alone their glycan modifications in complex biological samples such as the investigated CRC tumor tissues and PBMC samples comprising diverse cell populations remains a considerable challenge in glycobiology. Enabled by IHC and a new innovative correlation approach that made use of our quantitative -omics data from multiple CRC specimens and robust annotation of proteins to their respective cell- and tissue origins by the Human Protein Atlas^46^, in this study, we were able to suggest that the paucimannosidic signatures in CRC tumor tissues arise from multiple cellular sources including from both infiltrating monocytic and epithelial cancer cells present in the CRC tumor microenvironment (TME). Supporting these findings, we and others have indeed found paucimannosylation to be an abundant glycan feature in isolated blood monocytes and in macrophage effector cells derived from monocytes^27,47,48^ as well as in various CRC cell lines^15,43^. We have recently shown that cancerous epithelial cells frequently modify their lysosomal proteins with paucimannosidic glycans and have mechanistically detailed how the lysosomal content in such cells may be released to the extracellular environment via regulated exocytosis through focal adhesion points (unpublished observation, *under consideration*). However, the sheer abundance of paucimannose in blood monocytes (∼30% of the monocytic *N*- glycome)^27^ and their significant infiltration rate into the CRC TME^49^, point to the tumor-associated myeloid cells being principal contributors to the strong paucimannosidic features of the CRC tumor tissues. In line with these speculations, our IHC analysis demonstrated prominent co-localization of paucimannosidic epitopes with anti-inflammatory (CD163+) macrophages known to originate from tumor-infiltrating monocytes (Figure 2I). The observation of paucimannose-rich anti- inflammatory macrophages is particularly interesting as this macrophage sub-population is regarded to support tumor growth and spread^50^. However, we note that other cellular sources including pro-inflammatory macrophages and other cells in the CRC TME (e.g. tumor-infiltrating neutrophils, DCs, T-, NK-, B-cells and fibroblasts) may also contribute to the paucimannosidic signatures in the CRC tumor tissues, details that may now be explored using new spatial -omics approaches (e.g. MALDI-MS imaging), advanced microscopy (e.g. FRET) or cell sorting followed by ‘single-cell’ glyco(proteo)mics analysis^51–53^.

Recapitulating a newly discovered catabolic route for *N*-glycoprotein biosynthesis reported for human neutrophils^31^, in this study, we used immunocytochemistry and an established pharmacological Hex inhibitor (M31850) on a patient-derived CRC line (LIM2405) to document *in vitro* that a Hex-mediated truncation pathway catalyzing the formation of paucimannosidic proteins also exist in CRC cells. In parallel efforts, we have recently used *HEXA* and *HEXB* knock- out lines to demonstrate the existence of this non-canonical pathway in other cancer cells indicating that Hex-mediated paucimannosidic protein formation may be a biosynthetic pathway generally active across various cancer cell types (thus not unique to CRC) (unpublished observation, *under consideration*). In this present study, mass spectrometry and enzyme activity assays showed that Hex subunit β forming the Hex B isoenzyme form (ββ) dominates in the investigated biospecimens suggesting a negligible influence of the Hex A (αβ) and Hex S (αα) isoenzyme forms in CRC.

Importantly, using established functional assays, we documented that the adhesion, migration, and invasion potentials of the LIM2405 cells were dramatically reduced upon M31850-based Hex inhibition (Figure 3J), illustrating that the Hex enzyme and/or its paucimannosidic protein products are intimately involved in processes underpinning CRC metastasis. Longitudinal co-cultures of LIM2405 cells with anti-inflammatory macrophages supported the pro-tumorigenic role of Hex in CRC and suggested that the immune component is important for the Hex><CRC axis. While these *in vitro* observations are exciting as they indicate a functional relationship between Hex and/or paucimannosidic proteins with CRC progression and dissemination, future works are required to confirm this relationship *in vivo*, for example, by studying the effect of Hex disruption on cancer growth and spread in an orthotopic model in which *HEXB*-KO and wild-type CRC cells are xenografted in mice. Moreover, efforts are warranted to understand how Hex disruption and/or depletion of paucimannosidic proteins mechanistically reduce tumorigenic processes in CRC. Given that paucimannosidic proteins carry functional epitopes known to be recognized by a range of immuno-modulatory glycan-binding proteins (lectins) i.e. DC-SIGN, mannose receptor, and langerin^54,55^, it is tempting to speculate that aberrations of such glycan-lectin interactions may contribute to the reduced metastatic potential of the Hex inhibited CRC cells. In line with this hypothesis, macrophages, which were found to impact the proliferation of Hex inhibited CRC cells (Figure 3K), abundantly express such immuno-modulatory glycan-binding proteins^54,55^.

Alternatively, the Hex enzyme may directly influence the metastatic potential either by altering hydrolytic processes and pathways within the lysosomes of cells or upon release to the extracellular environment where Hex amongst other lysosomal hydrolases are regarded to assist the enzymatic degradation of the ECM important for cancer growth and EMT^56^. We also note that Hex inhibition led to considerable alterations within the cellular proteome with ∼200 proteins under-expressed in Hex inhibited versus untreated LIM2405 cells many of which known to be involved in cell-cell adhesion processes (Figure 3I). Thus, Hex may also indirectly contribute to pro-tumorigenic functions. Although unlikely to play a significant role in CRC cells that express limited amounts of gangliosides (relative to glycosphingolipid-rich neurons)^57^, we cannot at this stage rule out that the reduced metastatic potential of Hex inhibited LIM2405 cells may relate to an altered Hex-mediated elimination of gangliosides sent for degradation in the lysosomes as observed in Tay-Sachs and Sandhoff disease featuring a damaging build-up of incompletely degraded gangliosides (GM2 gangliosidosis) in neurons^58^. Regardless of the underpinning mechanisms and effector molecules, the discovery of pro-tumorigenic roles of Hex in CRC is interesting as it may open new avenues for therapeutic intervention in key disease-promoting processes.

Aberrations of the expression and activity of Hex in CRC are also interesting as they point to Hex as a potential molecular marker for key disease traits. In line with similar findings from other studies reporting that Hex is overexpressed in CRC cell lines and tissues^20,38,59^, we found that the Hex enzyme activity is elevated in PMBCs and plasma from CRC patients with advanced stages of the disease relative to matched controls suggesting associations to disease stage and severity. An even stronger link was found between the plasma Hex activity and patient survival (Figure 4E-F). By applying a simple Hex activity assay on neat plasma (2 µl/sample) from a sizeable CRC patient and donor cohort, we demonstrated that high plasma Hex activity correlates well with an increased disease risk as determined by a substantially lower five-year survival relative to CRC patients with lower plasma Hex activity levels. Importantly, the Hex><mortality relationship remained highly significant after correcting for age as a confounding factor for patient survival. Together with the fact that the MUG substrate-based Hex activity assay is simple, sensitive, and robust (Supplementary Figure S3), the close association between enzyme activity and mortality, suggest that Hex is a promising easy-to-assay marker candidate for disease risk stratification in CRC.

Serum and urine Hex were previously proposed as a putative marker for colon cancer^60–62^. However, the observation that several lysosomal glycoside hydrolases including Hex appear raised across a variety of conditions (thus not specific to CRC) limits the potential use of Hex as a diagnostic marker for CRC^63^. Given the observed link to patient survival as reported herein, we instead propose that the plasma Hex activity may have prognostic value in terms of disease risk stratification after the CRC diagnosis have been made and preferably at an early stage of the disease where treatment options are plentiful, and intervention impactful.

In conclusion, we have used new untargeted -omics methods and access to valuable patient samples to identify that paucimannosylation are unconventional glycosylation features closely associated with CRC progression. By targeting the paucimannose-producing enzyme, we then showed that Hex inhibition dramatically reduces the tumorigenic potential of CRC cells indicating new functional links between Hex, paucimannosylation and CRC. Importantly, the plasma Hex activity correlated with both disease stage and patient risk as measured by their five-year survival. Collectively, the findings open new avenues for effective prognostication and therapeutic intervention strategies in CRC.

## Acknowledgments

RK was supported by the Cancer Institute of New South Wales (ECF181259). NB and PD are supported by International Macquarie University Research Excellence Scholarships funded by Macquarie University. THC is supported by an International Research Training Program Scholarship funded by the Australian Government. SBA is supported by the Cancer Council New South Wales (RG23-06). MTA is the recipient of an Australian Research Council Future Fellowship (FT210100455).

## Author contributions

RK and MTA designed experiments. RK, LK, NB, and PD conducted experiments. RK, LK, NB, PD, SBA established laboratory methodologies. RK, LK, NB, PD, THC, BLW and MTA analyzed the data. RK, SBA, and MTA provided reagents for this study. RK and MTA wrote the manuscript. RK and MTA supervised this study and acquired funding. All authors have reviewed and approved the manuscript.

## STAR Methods

### CRC and donor biospecimens

The study was approved by the Human Research Ethics Committee (Medical Sciences) at Macquarie University, Sydney, Australia (Protocol 5201800073). All biospecimens including snap frozen and formalin-fixed paraffin-embedded (FFPE) tissues, PBMCs, and plasma were sourced from the Victorian Cancer Biobank (Melbourne, Australia). Matched snap frozen tissues, PBMCs and plasma were from 28 patients who were clinically diagnosed with CRC and their disease progression staged (I-IV) according to the TNM staging system (n = 7/stage). Additionally, seven paired adjacent normal tissues and one unpaired adjacent normal tissue (all snap frozen from CRC patients in stage I) were used as controls (n = 8). Furthermore, paired tumor and adjacent normal formalin-fixed paraffin-embedded (FFPE) tissue sections (thickness: 5 μm/slide, size: 2 cm x 2 cm) were obtained from one CRC patient in stage II. Finally, EDTA-plasma from 78 healthy donors and 302 CRC patients for whom five-year survival outcome and other key metadata were known were obtained from the Victorian Cancer Biobank with consent from all relevant parties. All biospecimens and technical approaches are summarized in **Supplementary Table S1**.

### LIM2405 cell cultures

The CRC patient-derived cell line LIM2405 (CVCL_4437, adenocarcinoma of the caecum of a male CRC patient) was cultured in RPMI-HEPES with 2 mM glutamine (Sigma) supplemented with 10% (v/v) fetal bovine serum (FBS, Invitrogen), 1% (v/v) penicillin streptomycin (Thermo), 1 ug/ml hydrocortisone (Sigma), 0.025 U/ml insulin (Sigma), and 0.01 µg/ml thioglycerol (Sigma) at 37°C in the humidified atmosphere containing 5% (v/v) CO2. LIM2405 cells were seeded (10^5^ cells/ml) in RPMI media into six-well plates (Corning) and incubated for two days at 37°C in 5% (v/v) CO2. After cells reached confluency, RPMI media was added containing different concentrations of M31850 in DMSO (0 µM, 0.24 µM, 7.81 µM, 15.62 µM, 31.25 µM) or different concentrations of DMSO alone (0 µM, 0.24 µM, 7.81 µM, 15.62 µM, 31.25 µM). All cells were incubated with M31850 or vehicle (DMSO) for 24 h, 37°C.

### Protein extraction

Snap frozen tissues were thawed. Tissue lysis and homogenization were performed in a lysis buffer containing 8 M urea in 50 mM triethylammonium bicarbonate (TEAB) (pH 8.5) and a protease inhibitor cocktail (Roche) using zirconium beads (3 mm diameter, Sigma) in a TissueLyser (30 Hz, 2 min, Qiagen). Protein concentrations were determined by bicinchoninic acid (BCA) (Thermo). PBMCs were thawed in serum-free RPMI medium, pelleted (500 g, 8 min, 20°C), washed with 1 ml PBS and centrifuged as above. The cell pellet was resuspended in 200 µl RIPA buffer containing PBS, 1% (v/v) Triton, 0.5% (w/v) sodium deoxycholate, 0.1% (w/v) SDS and a protease inhibitor cocktail (Roche). Cells were lysed on ice for 30 min followed by sonication using a Branson 450 Digital Sonifier (two cycles of 5–10 s, 30% output, on ice). Cell debris was pelleted (10,000 rpm, 10 min, 4°C) and supernatants containing the protein extract were transferred to new microtubes. Protein concentrations were determined by BCA (Thermo) ahead of protein precipitation using four volumes of acetone (16 h, −30°C). Proteins were pelleted (14,000 rpm, 10 min, 4°C), and resuspended in 8 M urea in 50 mM TEAB.

LIM2405 cells with or without Hex inhibition were detached with trypsin-EDTA (Sigma), washed with PBS and lysed with RIPA buffer. Proteins were quantified by BCA. Proteins (50 µg/sample) were precipitated using cold acetone for 16 h at −30°C. Samples were clarified by centrifugation (14,000 rpm, 10 min, 4°C). The pellet was resuspended in 8 M urea and 50 mM TEAB.

### Protein preparation for glycomics and glyco/proteomics

In preparation for the multi-omics experiments, protein extracts from tissues, PBMCs and LIM2405 cell line samples (50 µg/sample) were reduced using 10 mM DTT (30 min, 30°C) and alkylated using 40 mM iodoacetamide (final concentrations, 30 min, in the dark, 20°C). Alkylation reactions were quenched using excess DTT. Samples were split setting aside 15 µg protein extract for glycomics (see below) and 35 µg protein extract for glyco/proteomics. For the latter, samples were digested using sequencing grade porcine trypsin (1:50, w/w; 12 h, 37°C, Promega). Proteolysis was stopped by acidification using 1% (v/v) trifluoroacetic acid (TFA, final concentration). Peptides were desalted using primed Oligo R3 reversed phase solid phase extraction (SPE) micro-columns as described^64^ and dried.

### Glycan preparation for glycomics

In preparation for glycomics, proteins (15 µg/sample) were immobilized on a primed 0.45 μm polyvinylidene fluoride membrane (Merck Millipore) and handled as described^65^. Briefly, *N*- glycans were released using 10 U recombinant *Elizabethkingia miricola N*-glycosidase F (10 U/μl, 16 h, 37°C, Promega). Detached glycans were reduced with 1 M sodium borohydride in 50 mM potassium hydroxide (3 h, 50°C). Reactions were stopped using glacial acetic acid, and glycans were desalted using strong cation exchange/C18 and porous graphitized carbon (PGC) SPE micro- columns.

### TMT labelling of peptides

Peptides (35 µg/sample) were labelled with tandem mass tags (TMT). Separate peptide reference pools were generated for the tissue and PBMC samples by pooling peptides from all samples to enable quantitative comparisons across multiple TMT-10plex experiments. The individual peptide samples were then randomly combined across four TMT-10plex sets for both the tissue and PBMC samples, see **Supplementary Table S12** for design. Each set included at least one sample from each CRC stage and a control sample. The reference sample was consistently labelled with the 126 Da reporter ion channel. For both the tissue and PBMC experiments, a total of 36 peptide samples were labelled with TMT including seven samples from each CRC stage and 8 controls.

For each sample, peptides from 25 μg protein extract were dissolved in 100 μl 100 mM TEAB and labelled with TMT10plex tags (0.23 mg in 41 μl neat anhydrous ACN, 1 h, 20°C, Thermo). Labelling reactions were quenched using 8 μl 5% (v/v) hydroxylamine (15 min, 20°C). Labelled peptides were mixed 1:1 (w:w) and desalted using hydrophilic–lipophilic balance SPE cartridges (Waters). A small aliquot containing 10 µg peptide mixture was set aside for direct LC-MS/MS analysis (unenriched fraction) and dried. Remaining samples were dried for glycopeptide enrichment.

### Glycopeptide enrichment

TMT-labelled peptide mixtures (∼240 μg) were reconstituted in 50 μl 80% ACN in 1% (both v/v) TFA and loaded onto primed custom-made hydrophilic interaction liquid chromatography (HILIC) SPE micro-columns packed with zwitterionic HILIC resin (10 μm particle size, 200 Å pore size, kindly provided by Merck Millipore) onto supporting C8 disks (Empore) in p10 pipette tips as described^46,64^. Briefly, the flow-through/wash fractions containing non-glycosylated peptides were collected for separate downstream analysis, and the retained glycopeptides were eluted over three rounds with 0.1% (v/v) TFA, 25 mM ammonium bicarbonate, and then 50% (v/v) ACN. Peptide and glycopeptide fractions were separately dried, desalted on primed Oligo R3 reversed phase SPE micro-columns, aliquoted, and dried.

### High pH prefractionation of glyco/peptides

The peptide and glycopeptide fractions were resuspended in 50 μl 25 mM ammonium bicarbonate and separately loaded onto primed Oligo R2 reversed-phase SPE micro-columns packed on supporting C18 discs (Empore) in p10 pipette tips for high pH prefractionation as described^46,64^.

Briefly, following thorough washing steps, the glyco/peptides were sequentially eluted with 25 mM ammonium bicarbonate in 10% ACN (fraction 1), 25 mM ammonium bicarbonate in 20% ACN (fraction 2), and 25 mM ammonium bicarbonate in 60% ACN (fraction 3) and dried. The three fractions were resuspended in 0.1% (v/v) formic acid (FA) for separate LC–MS/MS analysis.

### LC-MS/MS

For glycomics, the released *N*-glycans were separated on an UltiMate 3000 HPLC system (Dionex) interfaced with a LTQ Velos Pro Linear Ion Trap mass spectrometer (Thermo). The glycans were loaded on a PGC HPLC capillary column (Hypercarb KAPPA, 5 μm particle size, 200 Å pore size, 180 μm inner diameter × 100 mm length, Thermo) operated at 50°C with a constant flow rate (4 μl/min) supplemented with a post-column make-up flow supplying pure ACN at 4 μl/min delivered by the HPLC system^66^. The mobile phases were ammonium bicarbonate (10 mM), pH 8.0 (solvent A) and 10 mM ammonium bicarbonate in 70% (v/v) ACN (solvent B) employing a 86 min-gradient: 8 min at 2.6% B, 2.6-13.5% B over 2 min, 13.5-37.3% B over 55 min, 37-64% B over 10 min, 64-98% B over 1 min, 5 min at 98% B, 98-2.6% B over 1 min, and 4 min at 2.6% B. The electrospray ionization source was operated in negative ion polarity mode with a source potential of 3.6 kV. Full MS1 scans (*m/z* 570-2,000) were acquired using one micro-scan, *m/z* 0.25 full width half maximum (FWHM) resolution, 5 × 10^4^ automatic gain control (AGC), and 50 ms maximum accumulation time. MS/MS data were acquired using *m/z* 0.35 FWHM resolution, 2 × 10^4^ AGC, 300 ms maximum accumulation time, and 2 *m/z* precursor ion isolation window. The five most abundant precursors were selected from each MS1 full scan for collision-induced dissociation–based MS/MS using a normalized collision energy (NCE) of 33% with an activation Q of 0.250 and 10 ms activation time.

For glyco/proteomics, TMT-labelled peptide mixtures were loaded on a trap column (2 cm × 100 μm inner diameter) custom packed with ReproSil-Pur C18 AQ 5 μm resin (Dr Maisch, Ammerbuch-Entringen, Germany) and separated on an analytical column (Reprosil-Pur C18-Aq; 25 cm × 75 μm, 3 μm ID, Dr Maisch) at 300 nl/min provided by an UltiMate 3000 RSLCnano HPLC system. The mobile phases were 99.9% ACN in 0.1% (both v/v) FA (solvent B) and 0.1% (v/v) FA (solvent A). The gradient was 2-30% B over 100 min, 30-50% B over 18 min, 50-95% B over 1 min, and 9 min at 95% B. The nanoLC was connected to a Q-Exactive HF-X Hybrid Quadrupole-Orbitrap mass spectrometer (Thermo) operating in positive ion polarity mode. The Orbitrap acquired full MS1 scans (*m/z* 350-1,800, AGC: 3 × 10^6^, 50 ms maximum accumulation, 60,000 FWHM resolution at *m/z* 200). Employing data-dependent acquisition, the 20 most abundant precursor ions from each MS1 full scan were isolated and fragmented utilizing higher-energy collision-induced dissociation (HCD, NCE 35%). Only multicharged precursors (Z ≥ 2) were selected for fragmentation. Fragment spectra were acquired in the Orbitrap (45,000 resolution, AGC: 1 × 10^5^, 90 ms maximum accumulation, *m/z* 1.0 precursor isolation window, 30 s dynamic exclusion after a single isolation/fragmentation of a given precursor).

### LC-MS/MS data processing and analysis

For glycomics, the glycans were manually identified and quantified as described^46,67,68^. Xcalibur (v2.2, Thermo) was used to browse and annotate the raw LC–MS/MS data. Briefly, glycan precursor ions were extracted using RawMeat (v2.1, Vast Scientific), common contaminants/redundant precursors removed, and generic monosaccharide compositions (Hex, HexNAc, dHex, NeuAc) established using GlycoMod (http://www.expasy.ch/tools/glycomod). The glycan fine structures were manually elucidated using monoisotopic mass, PGC LC elution time and MS/MS fragment patterns. Byos (v5.5.39) was used to annotate glycan fragment spectra and generate cartoons of the identified glycan structures. Glycan quantification was performed using extracted ion chromatograms in Skyline (v9.1) as described^68^.

For glycoproteomics, HCD-MS/MS data of glycopeptides were searched with Byonic (v2.6.46, Protein Metrics) using 10/20 ppm as the precursor/product ion mass tolerance, respectively. Cys carbamidomethylation (+57.021 Da) and N-term/Lys TMT (+229.163 Da) were considered fixed modifications. Fully tryptic peptides were searched with up to two missed cleavages per peptide. The following variable modifications were selected allowing two common and one rare modifications per peptide: Met oxidation (+15.994 Da, common) and *N*-glycosylation of sequon- localized Asn (rare) employing a glycomics-informed *N*-glycan search space^64^. HexNAc1, HexNAc1Fuc1, and HexNAc2 were manually added to the glycan search space. The HCD-MS/MS data were searched against all human proteins (20,300 sequences, UniProtKB, released December 11, 2019). All searches were filtered to <1% false discovery rate (FDR) at the glycoprotein level and 0% FDR at the glycopeptide level using a protein decoy database. Only confident glycopeptide identifications (PEP 2D < 0.001) were considered^46^. Glycopeptides were quantified using the ‘Report Ion Quantifier’ available as a node in Proteome Discoverer (v2.2, Thermo) as described^46^. Glycopeptides were manually grouped by summing the reporter ion intensities from the glycopeptide-to-spectrum matches (glycoPSMs) belonging to the same UniProtKB identifier, same glycosylation site within the protein, and same glycan composition. The abundances of the unique glycopeptides from each channel were first normalized by dividing each unique glycopeptide reporter ion intensity by the reference reporter intensity from the peptide reference pool (see above) within that specific experiment and further normalized by the total sum intensity of each channel to correct for any inter-sample variation introduced during labelling and mixing.

For proteomics, HCD-MS/MS data (from both unenriched peptides and HILIC flow-through fractions) were processed using MaxQuant (v1.6.12.0, https://www.maxquant.org) and searched against all human proteins (20,300 sequences, UniProtKB released December 11, 2019) using the Andromeda search engine^69^ with a tolerance of 4.5 ppm for precursor ions and 20 ppm for product ions. The enzyme specificity was set to trypsin with a maximum of two missed cleavages permitted.

Carbamidomethylation of Cys (+57.021 Da) and N-term/Lys TMT (+229.163 Da) were considered fixed modifications. Met oxidation (+15.994 Da) and protein N-terminal acetylation (+42.010 Da) were considered variable modifications. Both the protein and peptide level identifications were filtered to 1% FDR. Processing and statistical analyses (see below) of the MaxQuant output were performed in Perseus v2.0.7.0. Proteins identified using the reverse database, proteins only identified through modified peptides, and proteins identified from the MaxQuant contaminant database were excluded. By enabling the report fragment ions “10plex TMT” in the quantification settings, proteins were quantified by their reporter ion intensities using at least one razor/unique peptide per protein. The TMT-based protein abundances were normalized as described above.

### Immunohistochemistry (IHC)

IHC was performed on paired FFPE tissue slides (CRC tumor and adjacent normal tissue pairs) from a CRC patient in stage II using a previously published method^70^ with some modifications. Briefly, tissue slides were heated to 50°C for 40 min followed by three cycles of deparaffination in neat xylene (30 min/cycle, 20°C) and rehydration in 70-100% ethanol (3 min/cycle, 20°C). Deparaffinated tissues were washed in Tris-buffered saline (TBS) and antigens were heat-retrieved in a tris-ethylenetriaminetetracetic acid (EDTA) buffer containing 10 mM Tris base, 1 mM EDTA, 0.05% (w/v) Tween20, pH 9.0 (95°C, 30 min). Slides were cooled, washed with TBS, and blocked with 5% (w/v) bovine serum albumin (BSA, 99.9% w/w purity, 1 h, 20°C, Sigma) to limit unspecific antibody binding. Importantly, BSA was pre-treated with 100 mM sodium periodate (4°C, 2 h) and dialyzed overnight in TBS to eliminate glycoepitopes from any contaminating glycoproteins. BSA blocked slides were washed with TBS and then stained with antibody. A fluorescence-conjugated rabbit monoclonal anti-human IgG-based antibody targeting the epithelial cellular adhesion molecule (EpCAM) was used to stain epithelial cells (anti-EpCAM-Alexa Fluor 647 (AF647), 1:100 dilution, 1 h, 20°C, Abcam). To stain anti-inflammatory macrophages known to express CD163^71^, a rabbit monoclonal anti-human CD163 IgG-based antibody (1:100 dilution, 16 h, 4°C, Abcam) was applied, which was then visualized using a fluorophore-labelled goat monoclonal anti-rabbit IgG-Alexa fluor 405 (AF405) IgG (1:400 dilution, 1 h, 20°C, Abcam). As previously reported^24,25^, paucimannosidic glyco-epitopes were detected using a mouse IgM antibody ‘Mannitou’ (kindly provided by Prof Simone Diestel, Univ Bonn, Germany, 1:2 dilution of supernatant collected from cultured hybridomas, 2 h, 20°C). Mannitou was visualized using a monoclonal anti-mouse IgM produced in goat (1:400 dilution, 1 h, 20°C, Thermo) conjugated with Alexa fluor 488 (AF488). The tissue slides were washed four times between each of the antibody staining steps with warm TBS with 0.5% (w/v) Tween20 (TBS-T, 5 min/washing step) followed by a final TBS rinse for 5 min. To preserve fluorescence, coverslips were mounted to the tissue sections using antifade mounting media (Thermo). The tissue stains were visualized, and co- localization qualitatively assessed using an OlympusFV-3000 confocal microscope with 40x and 60x magnification using oil immersion objectives. Image data were handled using cellSens imaging software (Olympus, Japan).

### Cell viability assay

LIM2405 cells (5 × 10^4^ cells/ml) were seeded into 96-well microplates (Greiner 655160, tissue culture grade, flat bottom) in 100 μl culture media without Hex inhibitor (DMSO vehicle only) or with Hex inhibitors (M31850 in DMSO) spanning an extended concentration range (0.24, 7.81, 15.62, 31.25, 62.50, 125). Cells were incubated for 24 h at 37°C in 5% (v/v) CO2 and 10 μl 3-(4,5- dimethylthiazol-2-yl)-2,5-diphenyltetrazolium bromide (MTT) reagent were then added in the final concentration of 0.5 mg/ml to each well. Cells were incubated for another 3 h at 37°C, 5% (v/v) CO2. After removing the media, 100 µl solubilization reagent containing 40% (v/v) dimethylformamide, 2% (v/v) glacial acetic acid, 16% (w/v) SDS (pH 4.7) was added to each well and the plate incubated for 15 min at 20°C under gentle agitation. Absorbance of the formazan product was measured with Fluostar Galaxy (BMG) at 450 nm. The viability of Hex inhibitor treated cells was determined in technical triplicates based on absorbance readings relative to untreated (DMSO only) cells after subtracting the absorbance measured in blank samples.

### Cell adhesion assay

Cell adhesion experiments were performed as previously described^72^ but with minor modifications. LIM2405 cells (3 × 10^5^ cells/ml) were plated into six-well plates and incubated in 1 ml media for 24 h with different concentrations of M31850 (0 µM, 0.24 µM, 7.81 µM, 15.62 µM, 31.25 µM). Cells not exposed to Hex inhibitor were treated with different concentrations of DMSO (0 µM, 0.24 µM, 7.81 µM, 15.62 µM, 31.25 µM). All cells were incubated for 16 h at 37°C and 5% (v/v) CO2. Meanwhile, a 96-well plate was coated with Matrigel™ (2 μg/well; Corning) for 16 h at 4°C, washed three times with PBS and blocked with 3% (w/v) BSA (Sigma) for 2 h, 20°C. Cells were then washed with PBS, trypsinized and seeded in the pre-made Matrigel-coated plate (10^4^ cells/well). Cells were incubated for 1 h at 20°C in serum-free medium supplemented with 3% (w/v) BSA, washed 3 times in PBS and fixed with 10% (v/v) formaldehyde. Fixed cells were washed with PBS and stained with 1% (w/v) toluidine blue containing 1% (w/v) borax for 5 min, 20°C. The dye was solubilized in 100 μl 1% (w/v) SDS for 5 min, 20°C. Absorbance (in effect, cell adhesion) was measured at 620 nm in technical duplicates. Three independent experiments were performed for cells treated for 24 h with 31.25 µM M31850 in DMSO or with 31.25 µM DMSO only.

### Cell migration assay

LIM2405 cells (10^5^ cells/ml) were seeded in RPMI media into six-well plates (Corning, Australia) and incubated for two days at 37°C in 5% (v/v) CO2. Upon reaching confluency, RPMI media was added containing different concentrations of Hex inhibitor M31850 (0 µM, 0.24 µM, 7.81 µM, 15.62 µM, or 31.25 µM). As vehicle controls, different concentrations of DMSO (0 µM, 0.24 µM, 7.81 µM, 15.62 µM, or 31.25 µM) were added to cells prepared in other wells. All cells were incubated for 24 h, 37°C. Cells were detached with trypsin-EDTA (Sigma), washed three times with PBS and plated (10^4^ cells/ml) in serum-free RPMI media in the upper chambers of 8 mm pore Transwell (HTS Transwell 96-well plate, Corning). 150 µl RPMI media containing 1% (v/v) FBS was added to the bottom chamber of the Transwell plate and plates were incubated at 37°C at 5% (v/v) CO2 for 16 h. After incubation, the non-migrated cells in upper Transwell chamber were washed with PBS and removed using cotton swabs. Migrated cells under the 0.8 mesh membrane in Transwell chamber were fixed in 10% (v/v) formaldehyde in PBS for 10 min. Cells were washed three times with PBS and stained for 5 min in 1% (w/v) toluidine blue in PBS. Excess toluide blue was removed by MilliQ washing and the wells were further cleaned with cotton swabs. Transwell chambers were placed into 96 well plates containing 200 µl 1% (v/v) SDS and incubated for 15 min at 37°C. The released toluidine blue (in effect measuring migrated cells) was measured twice (technical duplicates) at 595 nm with Fluostar Galaxy. The dose-dependent migration experiments were performed without replicates (n = 1), while four independent migration experiments (n = 4) were performed for cells treated with 31.25 µM M31850 in DMSO or with 31.25 µM DMSO only.

### Cell invasion assay

LIM2405 cells (10^5^ cells/ml) were seeded in RPMI media into six-well plates (Corning, Australia) and incubated for two days at 37°C and 5% (v/v) CO2. After cells reached confluency, RPMI media was added containing different concentrations of M31850 in DMSO (0 µM, 0.24 µM, 7.81 µM, 15.62 µM, 31.25 µM) or different concentrations of DMSO alone (0 µM, 0.24 µM, 7.81 µM, 15.62 µM, 31.25 µM). All cells were incubated for 24 h, 37°C. Cells were detached with trypsin-EDTA (Sigma), washed three times with PBS and plated (10^4^ cells/ml) in serum free RPMI media in the upper chambers of 8 mm pore Transwell (HTS Transwell 96-Well Plate, Corning) pre-coated with Matrigel™ (2 μg/well; Corning). Following the same protocol for the migration assay, cells were allowed to invade the Matrigel for 16 h at 37°C in 5% (v/v) CO2, washed, fixed and stained. Toluidine blue was released from cells with 1% (v/v) SDS and the intensity of the stain (in effect measuring invading cells) was measured twice at 595 nm with Fluostar Galaxy. Five independent experiments (n = 5) were performed for cells treated with 7.81 µM M31850 in DMSO or with 7.81 µM DMSO only.

### Isolation of human monocytes, macrophage differentiation and co-cultures with LIM2405

PBMCs were isolated from buffy coat of de-identified healthy donors obtained from Australian Red Cross (Sydney, Australia) with approval from Macquarie University Human Research Ethics Committee (#52021780424399). PBMCs were isolated using a Lymphoprep density gradient (StemCell Technologies #07801) and centrifugation at 1,200 x *g* for 20 min at room temperature. The intermediate buffy layer was collected and washed with PBS containing 2 mM EDTA and 1% (w/v) BSA. Monocytes were isolated through CD14 positive selection using a Pan monocyte isolation kit (Miltenyi Biotec, #130096537) as described^73^. The monocyte purity (typically >95%) was validated by flow cytometry.

For macrophages differentiation, 10 ng/mL granulocyte-macrophage colony-stimulating factor (GM-CSF; Miltenyi Biotec, #130093865) was added to isolated monocytes and cells were incubated for 10 days in RPMI 1640 culture medium (Gibco #11875093) supplemented with 10% (v/v) heat-inactivated FBS (Gibco), 1% sodium pyruvate, 1% penicillin/streptomycin, 1% (all w/v) non-essential amino acid (Thermo). Polarization into anti-inflammatory macrophages was performed using 20 ng/mL IL-4 and 20 ng/ml IL-13 (Miltenyi Biotec) as described^74^ and confirmed with CD206 marker expression through flow cytometry.

For LIM2405 and macrophage co-cultures, LIM2405 cells were pre-incubated for 24 h with 7.8 μM M31850 or DMSO as a vehicle control, before the media was removed, the cells washed and then transferred to the primary anti-inflammatory macrophages in a 1:10 (mol/mol) ratio of LIM2405 cells to CD163+ macrophages. The above-mentioned culture medium was used to culture both LIM2405 cells alone and with the anti-inflammatory macrophages for up to 96 h.

### Brightfield microscopy of co-culture cells

LIM2405 cell growth was monitored every 24 h and images captured using an Olympus CKX41 inverted microscope, equipped with Olympus DP21 camera. The light source was a 6 V, 30 W halogen lamp. The microscope was configured for brightfield imaging using Koehler illumination and images were acquired and handled with the Olympus DP2-BSW software. The average cell numbers across two fields of view were manually determined for all conditions. For each condition, the same two fields were longitudinally monitored.

### Immunocytochemistry

LIM2405 cells were seeded (10^5^ cells/ml) into six-well plates (Corning) containing sterile poly-L- lysine coated coverslips on the bottom of the well and grown for 24 h at 37°C in 5% (v/v) CO2. Cells were washed with PBS and fixed with 4% (v/v) paraformaldehyde in PBS (pH 7.4) for 20 min, 20°C. After three washes with PBS, coverslips were treated with 0.2% (v/v) Triton X-100 in PBS with 5% (v/v) horse serum for 20 min, 20°C. Cells were incubated with anti-Hex B antibody (#PAA637Hu01, 1:500, Clod-Clone Corp., TX, USA) for 30 min, 20°C. Cells were washed three times with PBS and incubated with secondary antibody anti-rabbit Alexa-647 (1:1,000 dilution, Invitrogen) for 30 min in the dark. In the same coverslips, undiluted Mannitou (see above) was added and incubated for 30 min, 20°C. Cells were washed with PBS and incubated for another 30 min with anti-mouse IgM Alexa Fluor 488-conjugated as a secondary antibody (1:500 dilution, Thermo). After washing the cells with PBS, coverslips were mounted using with ProLong™ Gold Antifade Mountant media with DAPI (Invitrogen) to preserve fluorescence. Cells were visualized and co-localization qualitatively assessed using a fluorescence microscope Olympus FluoView FV 3000RS IX83 (Olympus, Japan) with a UPLSAPO60XS objective. Measurements were performed in technical duplicates.

### N-acetyl-β-D-hexosaminidase (Hex) activity assay

Hex enzyme activity was established in LIM2405 (2 µg protein extract), PBMCs (1.5 µg protein extract) and in neat plasma (2 µl/sample). Samples were incubated with 30 μl prewarmed 3 mM 4- methylumbelliferyl-2-acetamido-2-deoxy-β-D-glucopyranoside (MUG) or 4-methylumbelliferyl- 2-acetamido-2-deoxy-β-D-glucopyranoside-6-sulfate (MUGS) substrate (Merck Millipore) in phosphate–citrate buffer (pH 5.0, 37°C, 30 min, in the dark) as described^75^. Reactions were stopped by adding 200 µl 0.25 M glycine carbonate buffer (pH 10.0). Hex activity was measured by fluorometric quantitation of the fluorogenic product 4-methylumbelliferone (4-MU) using a plate reader (FLUOstar Optima, BMG Technologies) with excitation at 360 nm and emission at 450 nm. Released 4-MU was quantified using a standard curve made from known amounts of 4-MU (Merck Millipore). The Hex activity was determined in technical triplicates and normalised within each plate assay based on control samples. Three independent experiments (n = 3) were performed for LIM2405 cells treated with 31.25 µM M31850 in DMSO or with 31.25 µM DMSO only. Hex activity was measured across 36 PBMC samples and 380 plasma samples (**Supplementary Table S11**).

### Hex stability test

The stability of Hex was determined using blood from healthy volunteers collected by venipuncture into either empty (for serum) or EDTA- or heparin-containing (for plasma) collection tubes (Vacutainer, Becton Dickinson Co, Australia). Samples were stored for 20 min, 20°C and plasma/serum separated from other blood components by centrifugation (1500 x g, 15 min, 20°C).

Samples were aliquoted and stored for either 4 h, 24 h, or 96 h at 20°C, 4°C, -20°C or -80°C until analyzed for Hex activity as per above. The impact of freeze/thaw cycles was also measured.

## Statistical information

Pathway enrichment analysis was performed using DAVID Bioinformatics^76^ (National Institute of Health) with corrected *p* < 0.05 as significance threshold. Bubble graphs and correlation plots were performed using SRplot^77^. Associations were explored using Pearson correlation coefficient and tested for significance using t distribution tests with *p* < 0.05 as confidence threshold. For the - omics datasets and functional assays, significance was generally assessed by unpaired two-tailed Student’s t-tests or ANOVA corrected for multiple comparisons using Tukey test with significant threshold set to 0.05 (95% confidence interval). Five-year survival analysis was performed using Kaplan-Meier survival plots and log-rank tests (*p* < 0.05). GraphPad Prism (v9.4.1, Dotmatics) or Perseus v2.0.7.0 were used for statistical analysis. Cox proportional-hazards model performed in the R software was used to determine associations between survival, age and Hex activity.

## Data availability

The glycoproteomics and proteomics LC-MS/MS raw data have been deposited to the ProteomeXchange Consortium via the PRIDE^78^ partner repository with the dataset identifiers: PXD051882 (tissues), PXD051907 (PBMCs) and PXD051909 (LIM2405 cell lines). Glycomics LC-MS/MS raw data were deposited to GlycoPOST^79^ with the identifiers: GPST000423 (tissues), GPST000424 (PBMCs) and GPST000425 (LIM2405 cell line). See **Supplementary Table S12** for overview of data files and the accession numbers of the available data.

---

**Supplementary Figure S1.**
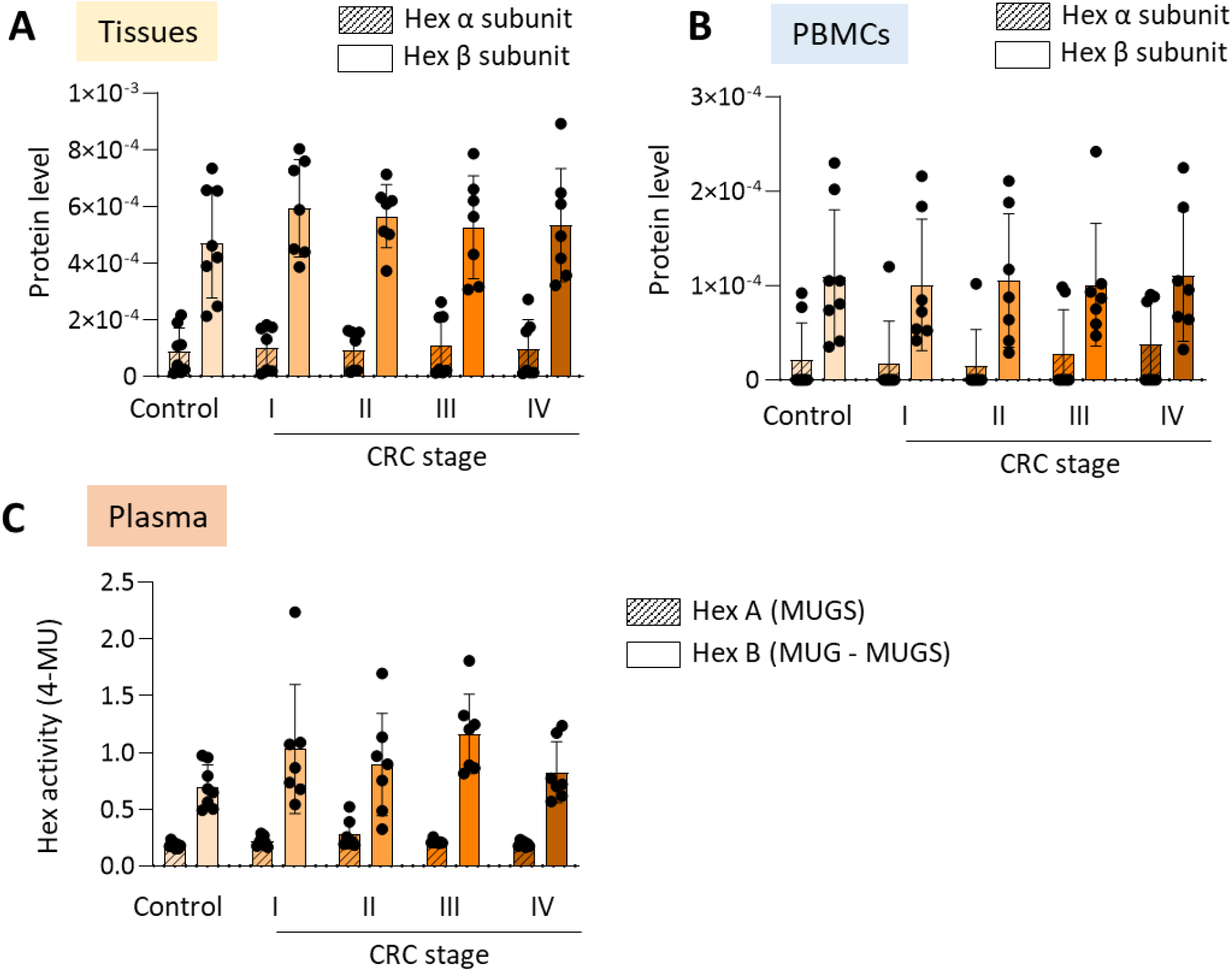
The Hex B (ββ) isoenzyme form dominates in CRC. Relative expression of the Hex α- (crossed) and β- (open) subunits in **A**) tumor tissues and **B**) PBMCs from CRC patients (n = 7/stage) and matching controls (n = 8). **C**) The activity of Hex A (αβ, measured by MUGS) and Hex B (ββ, measured by MUG – MUGS) isoenzyme forms in plasma from the same sample cohort.

**Supplementary Figure S2.**
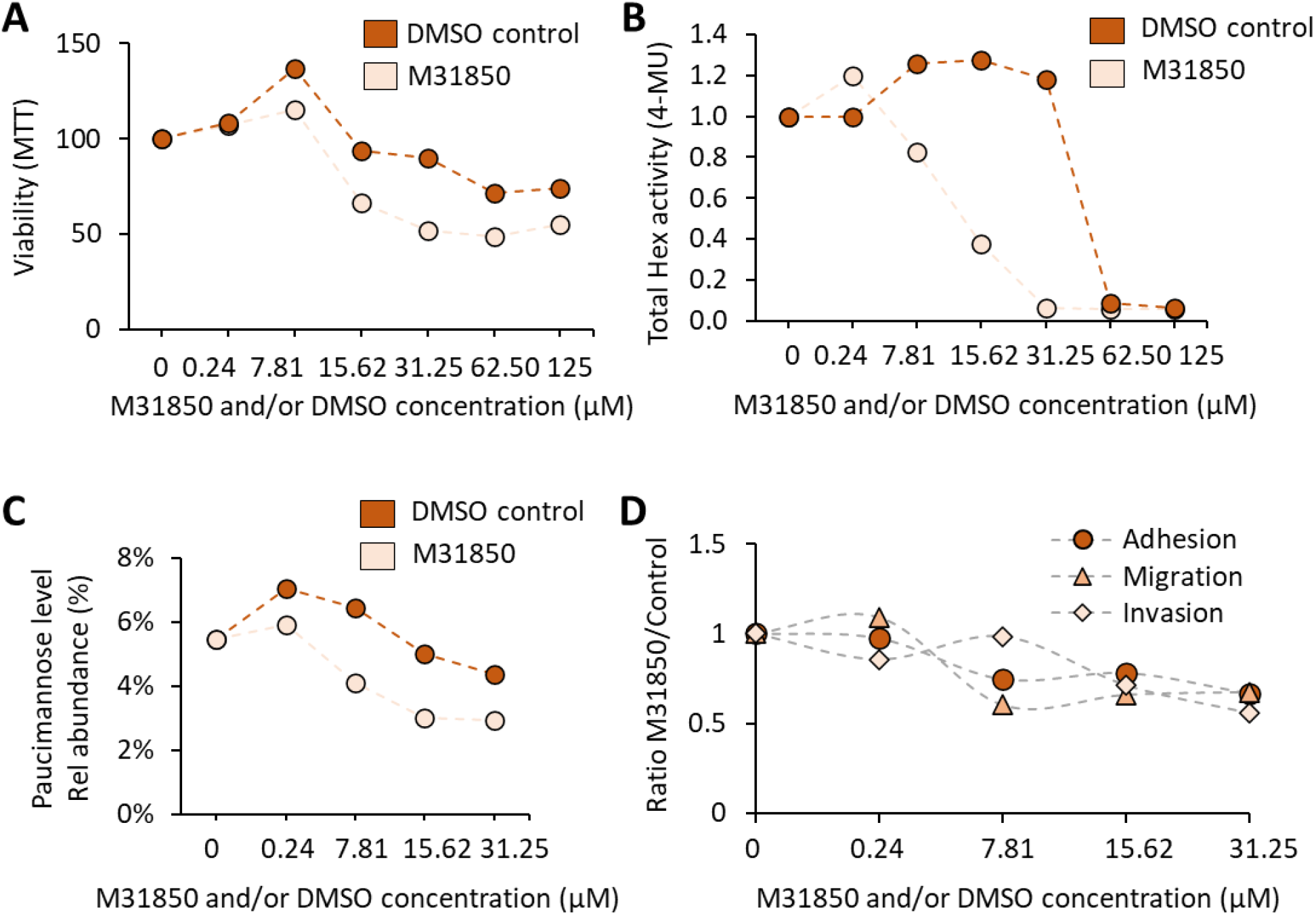
Impact of Hex inhibitor (M31850) and vehicle (DMSO) on LIM2405 phenotype. LIM2405 cells were treated for 24 h with different doses of M31850 in DMSO (0-125 µM, light orange) or with different concentrations of DMSO alone (0.24-125 µM, dark orange) after which several phenotypic features were measured including: **A**) Cell viability measured using an MTT assay. **B**) Total Hex activity measured using a MUG substrate assay. **C**) Total level of paucimannosylation measured by glycomics. **D**) Cell adhesion, migration and invasion measured using established functional assays.

**Supplementary Figure S3.**
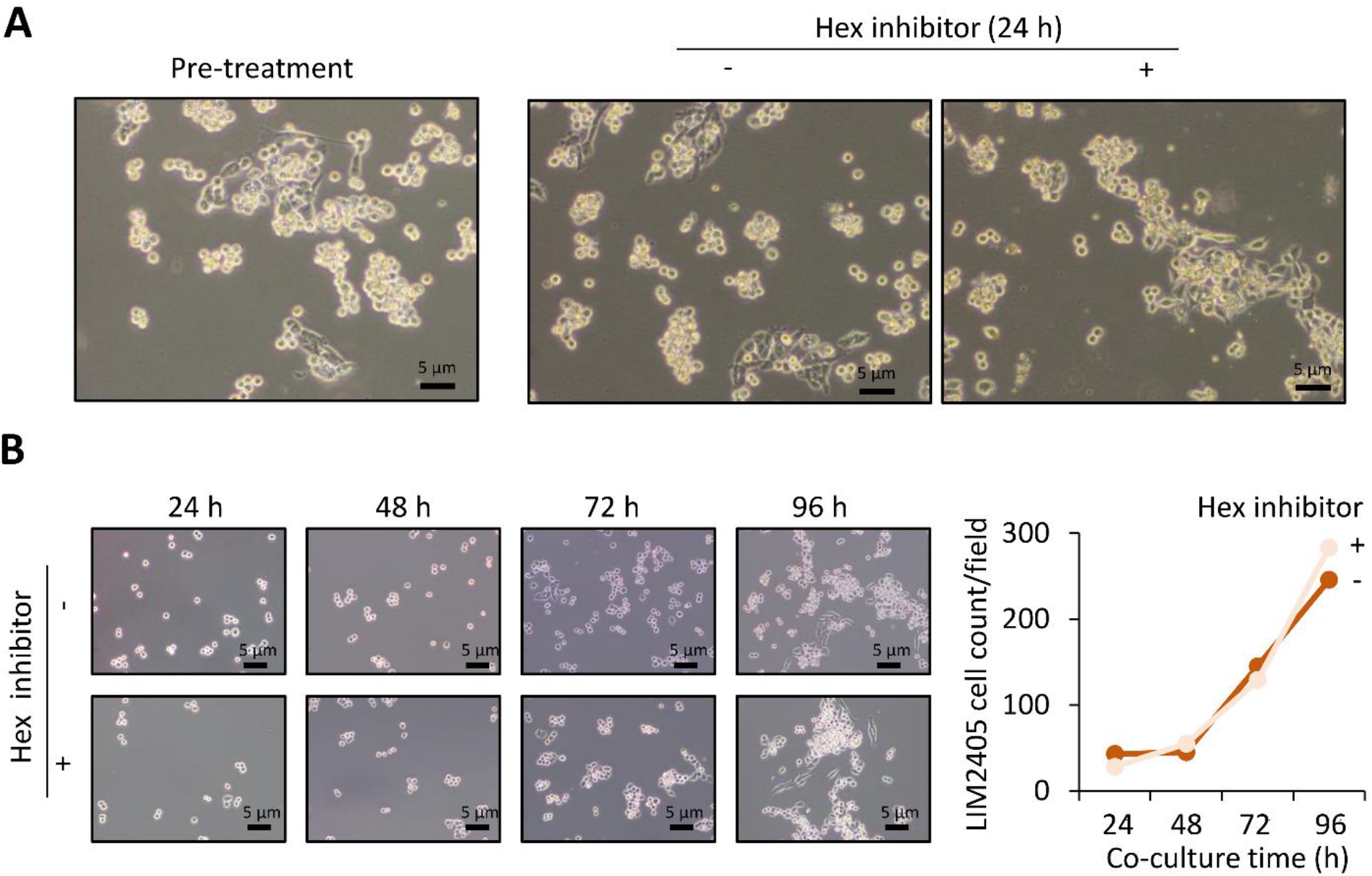
Unaltered LIM2405 cell proliferation upon Hex inhibition in the absence of anti-inflammatory macrophages. **A**) Representative microscopy images of LIM2405 cell line cultured in RPMI media before (Pre-treatment) and after 24 h treatment without (-) or with (+) Hex inhibitor (7.8 µM M31850 in DMSO). **B**) LIM2405 pre-treated for 24 h with or without Hex inhibitor (7.8 µM M31850 in DMSO) was cultured for 96 h in macrophage media (but in the absence of macrophages) to investigate the influence of Hex inhibition on LIM2405 proliferation without the immune component. Counts of LIM2405 cells were plotted as average of two fields. Images were obtained with 5x magnification.

**Supplementary Figure S4.**
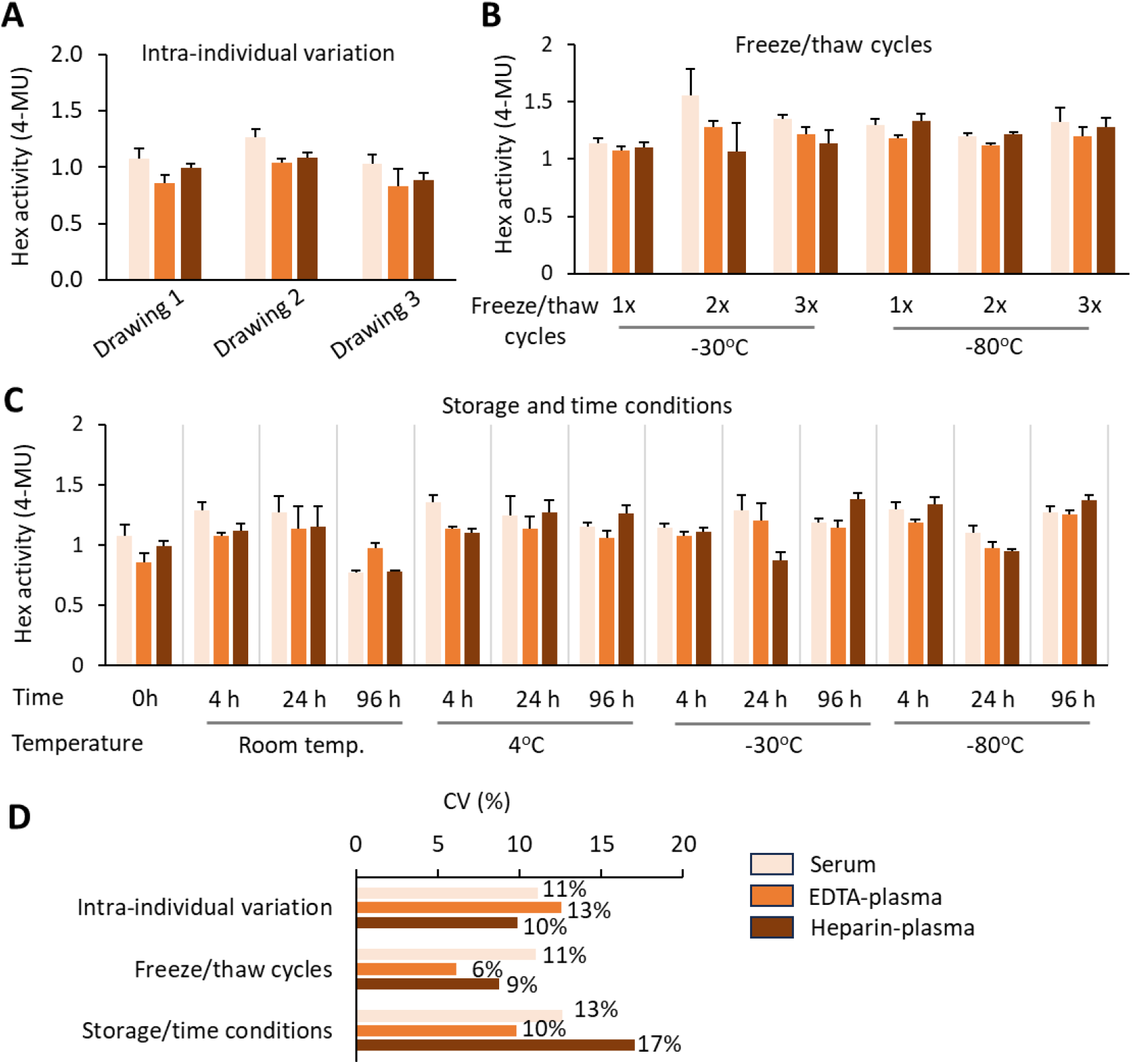
Robustness of the Hex activity assay. **A)** Blood was collected from the same healthy donor over three different days (5-7 days interval). The total Hex activity (MUG) of these samples was determined in technical triplicates using identical conditions. **B**) Hex activity was measured after the sample was subjected to one, two or three freeze/thaw cycles when stored at −30°C or −80°C. **C**) The Hex activity was measured from the same biological sample (processed into serum, EDTA- and heparin-plasma) immediately after collection, or after 4 h, 24 h or 96 h of storage. Different storage conditions were compared including room temperature, 4°C, −20°C, and −80°C. **D**) Coefficient of variation (CV) across multiple assays and blood collection methods.

**Supplementary Table S1.**
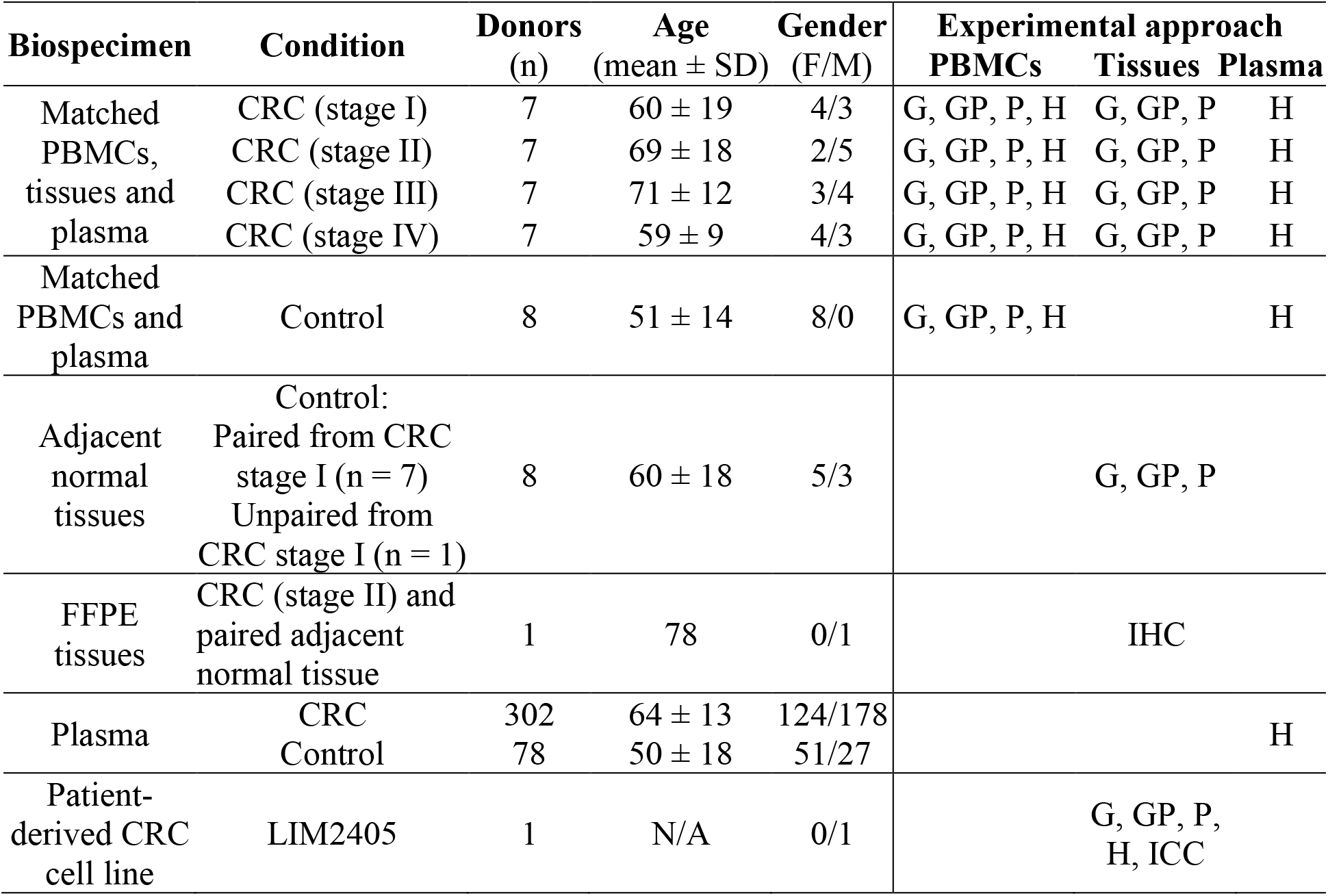
Sample cohort and experimental approaches used in this study. Gender (F: female, M: male). Experimental approaches (G: glycomics, GP: glycoproteomics, P: proteomics, H: Hex enzyme activity, IHC: immunohistochemistry, ICC: immunocytochemistry).

**Supplementary Table S2.** N-glycomics data of CRC and adjacent normal tissues.

**Supplementary Table S3.** N-glycomics data of CRC and normal PBMCs.

**Supplementary Table S4.** N-glycoproteomics data of CRC and adjacent normal tissues.

**Supplementary Table S5.** N-glycoproteomics data of CRC and normal PBMCs.

**Supplementary Table S6.** N-glycoproteomics data of LIM2405 with and without Hex inhibition.

**Supplementary Table S7.** Proteomics data of CRC and adjacent normal tissues.

**Supplementary Table S8.** Proteomics data of CRC and normal PBMCs.

**Supplementary Table S9.** N-glycome data of LIM2405 cells with and without Hex inhibition.

**Supplementary Table S10.** Proteomics data of LIM2405 with and without Hex inhibition.

**Supplementary Table S11.** Plasma cohort and total Hex activity.

**Supplementary Table S12.** Overview of experiments, obtained raw files and search parameters.

**Supplementary Data S1** (PDF). Glycan spectral evidence.

