## Supplementary Data for "Tumor- and immune-derived *N*-acetyl-β-D-hexosaminidase drive colorectal cancer and stratify patient risk"

#### Annotation and Fragmentation Key

- Fucose (Fuc)
- Galactose (Gal)
- Glucose (Glc)
- Mannose (Man)
- *N*-acetylglucosamine (GlcNAc)
- *N*-acetylgalactosamine (GalNAc)
- *N*-acetylneuraminic acid (NeuAc, sialic acid)

- Indicates mostly Y ions
- Indicates mostly Z ions
- Reduced reducing end
- -Ac Indicates potential loss of acetyl group

### 01 (HexNAc)2 (Deoxyhexose)1

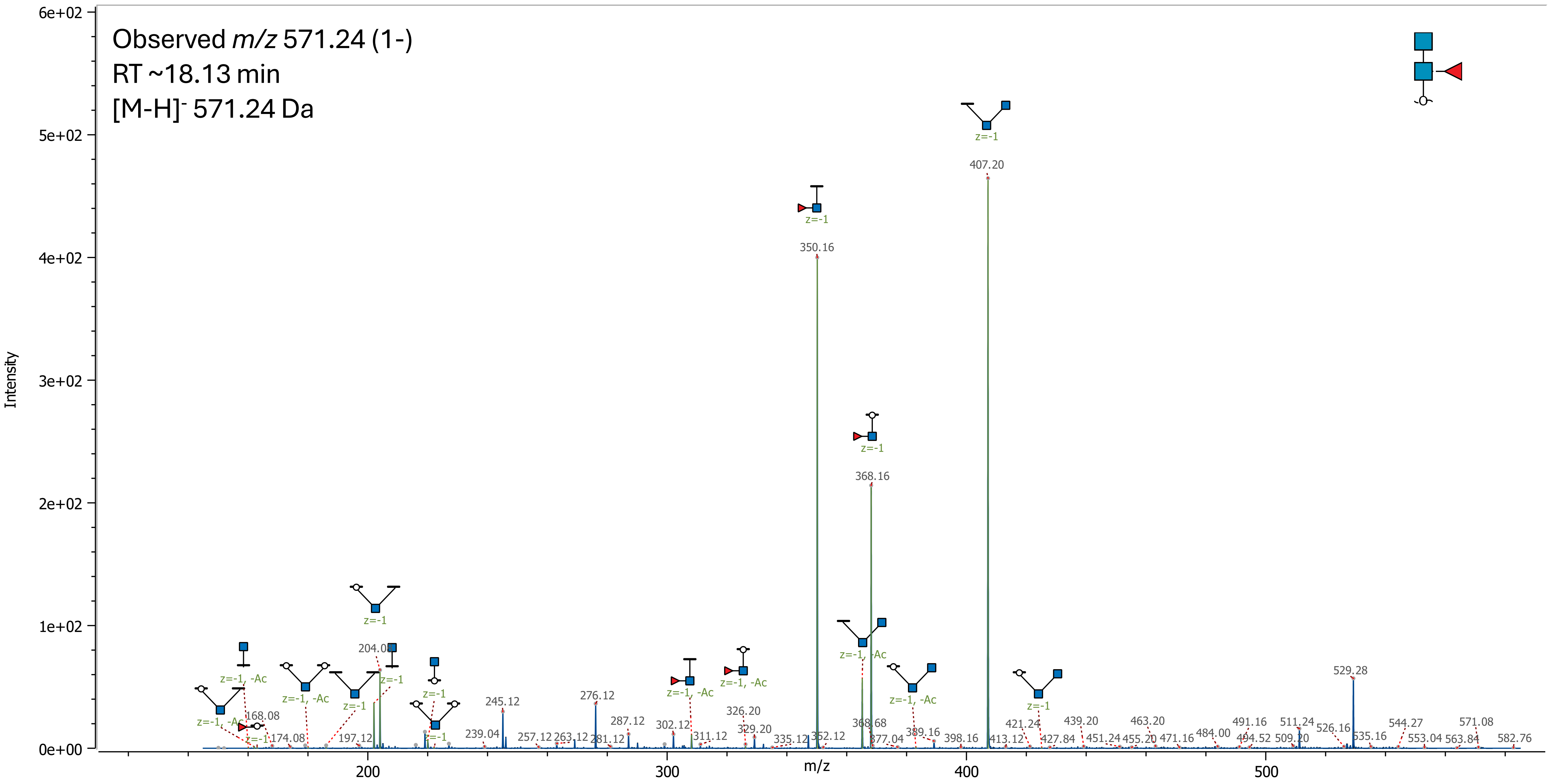

02 (Hex)1 (HexNAc)2

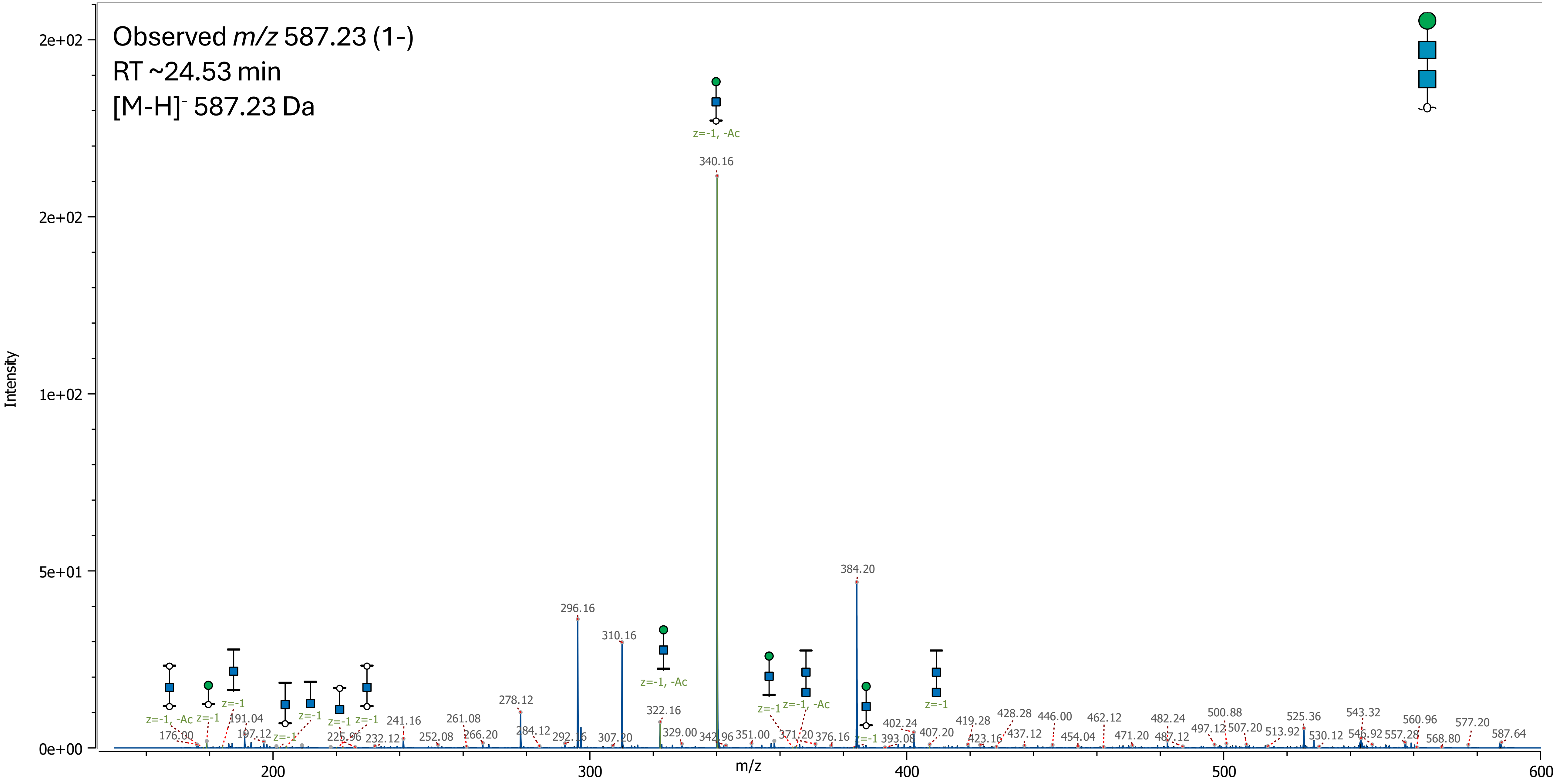

03 (Hex)1 (HexNAc)2 (Deoxyhexose)1

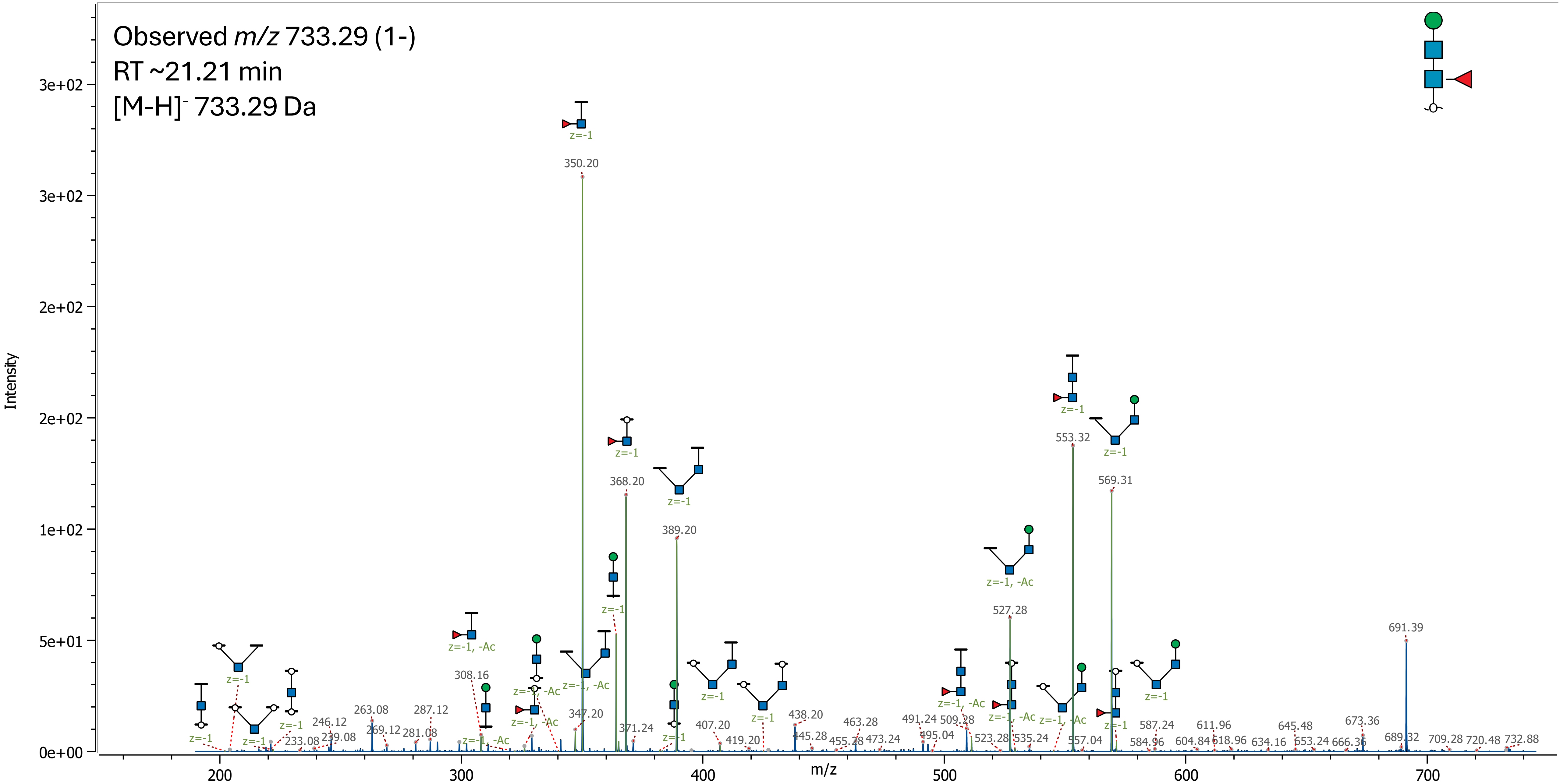

04 (Hex)2 (HexNAc)2

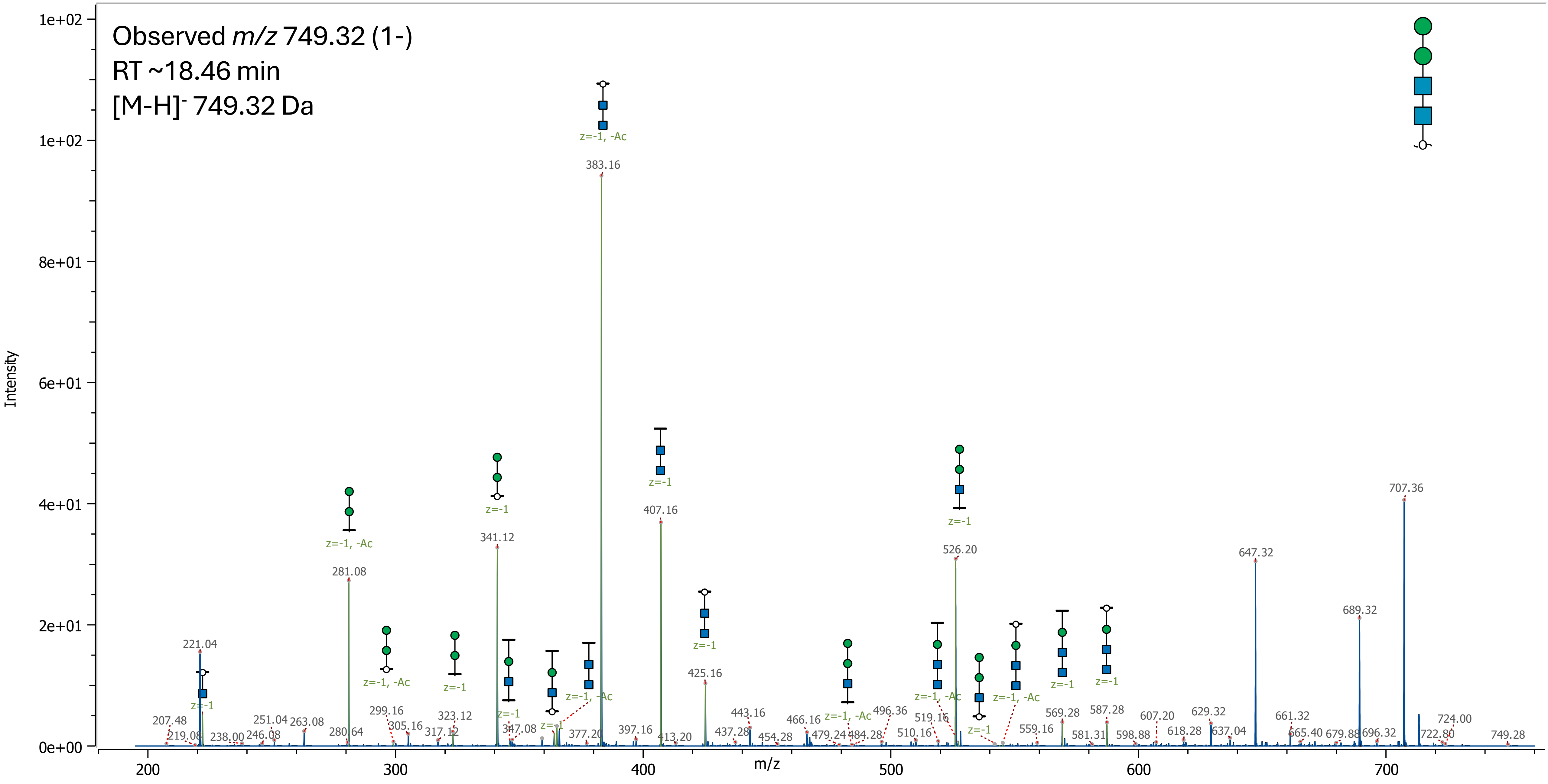

05 (Hex)2 (HexNAc)2 (Deoxyhexose)1

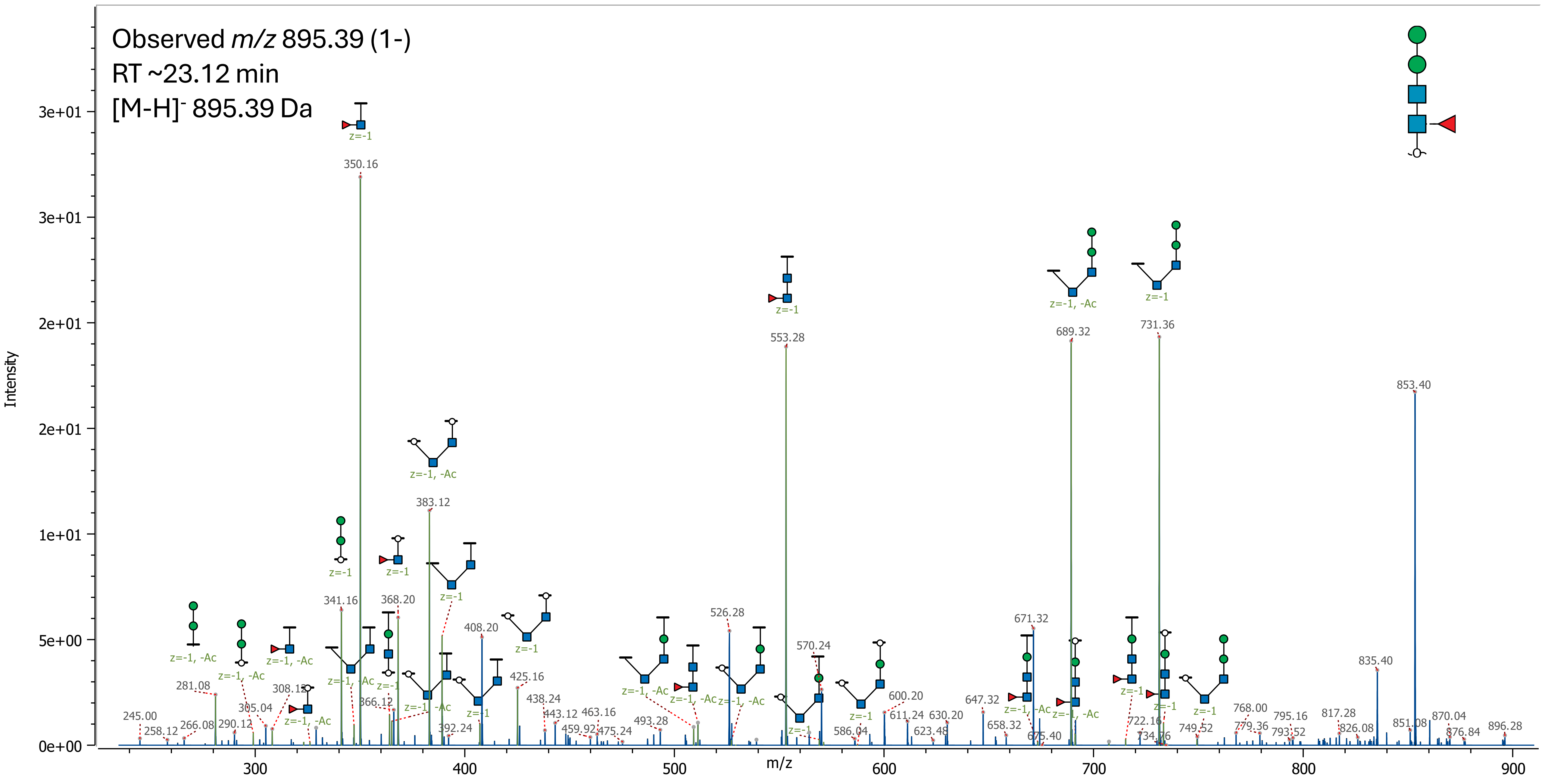

06 (Hex)3 (HexNAc)2

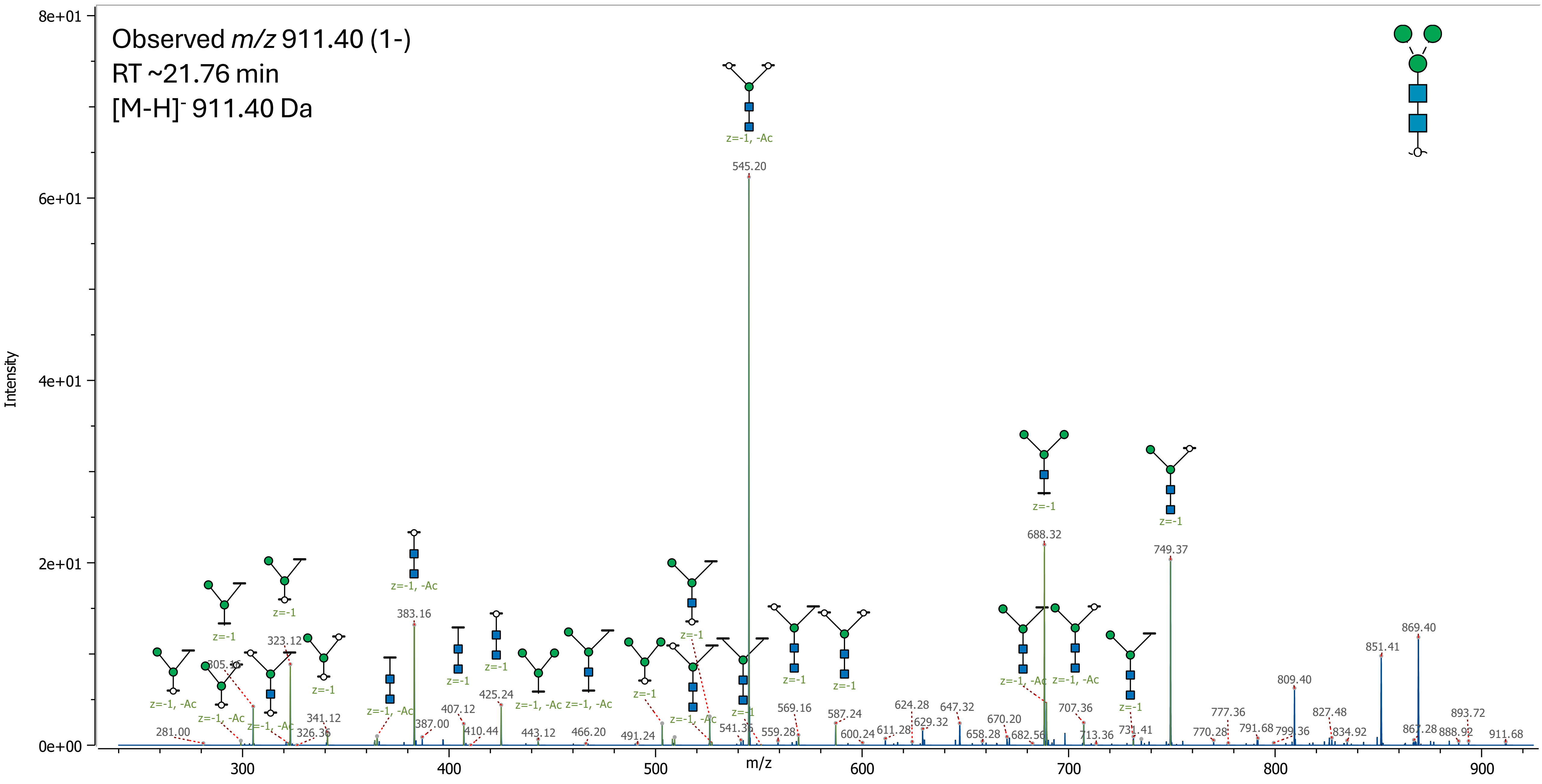

### 07 (Hex)3 (HexNAc)2 (Deoxyhexose)1

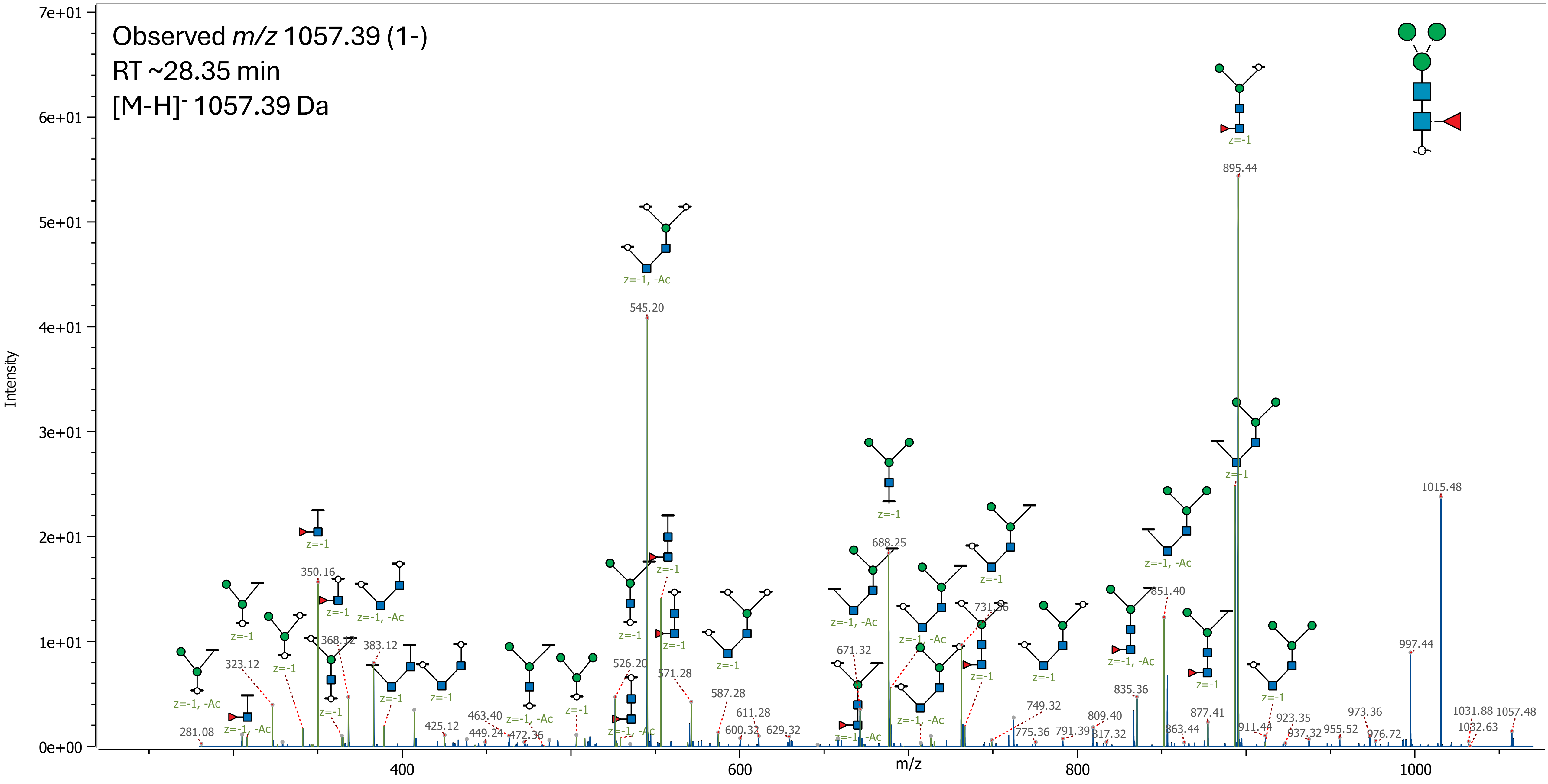

08 (Hex)4 (HexNAc)2

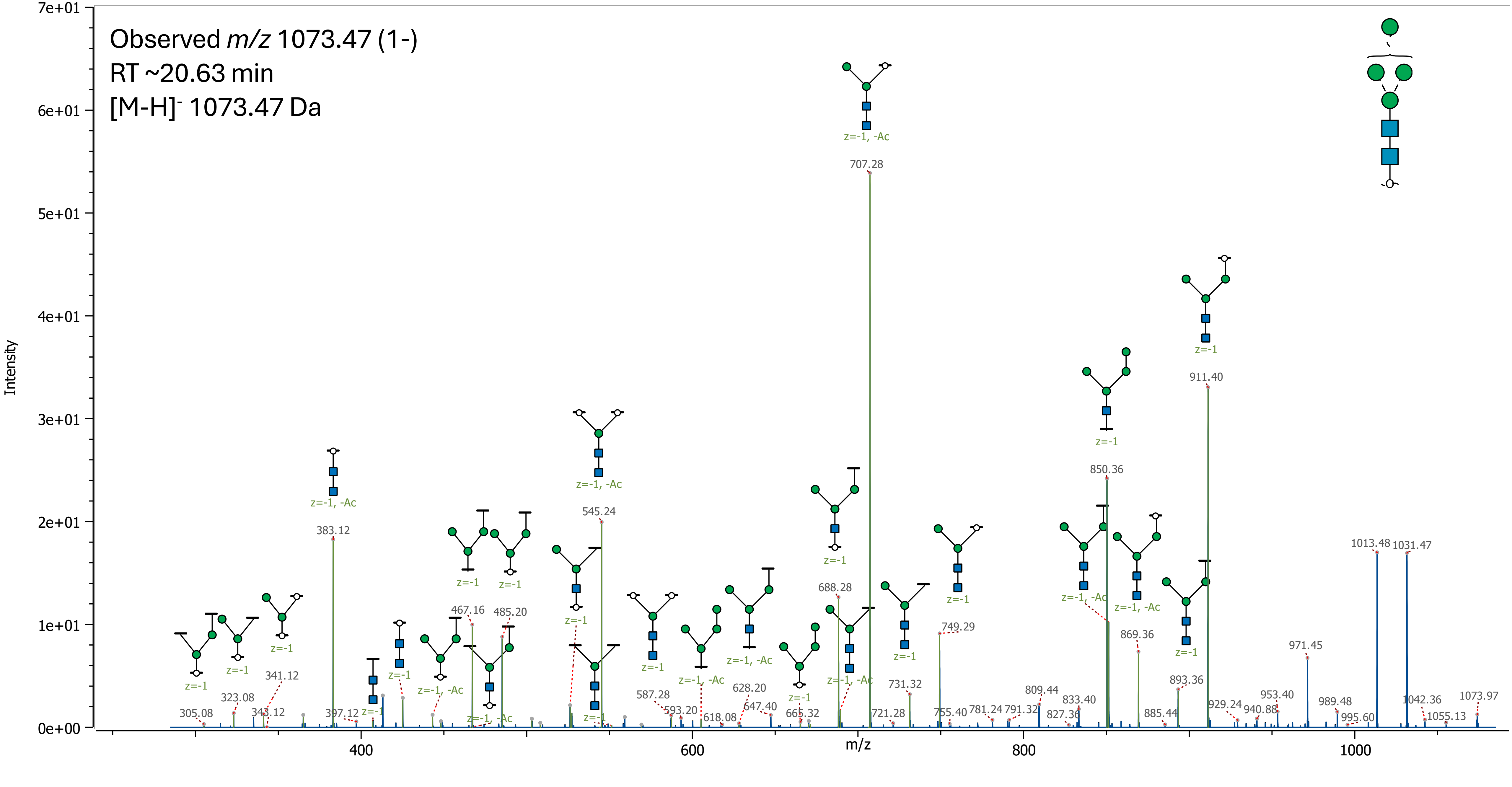

09 (Hex)4 (HexNAc)2 (Deoxyhexose)1

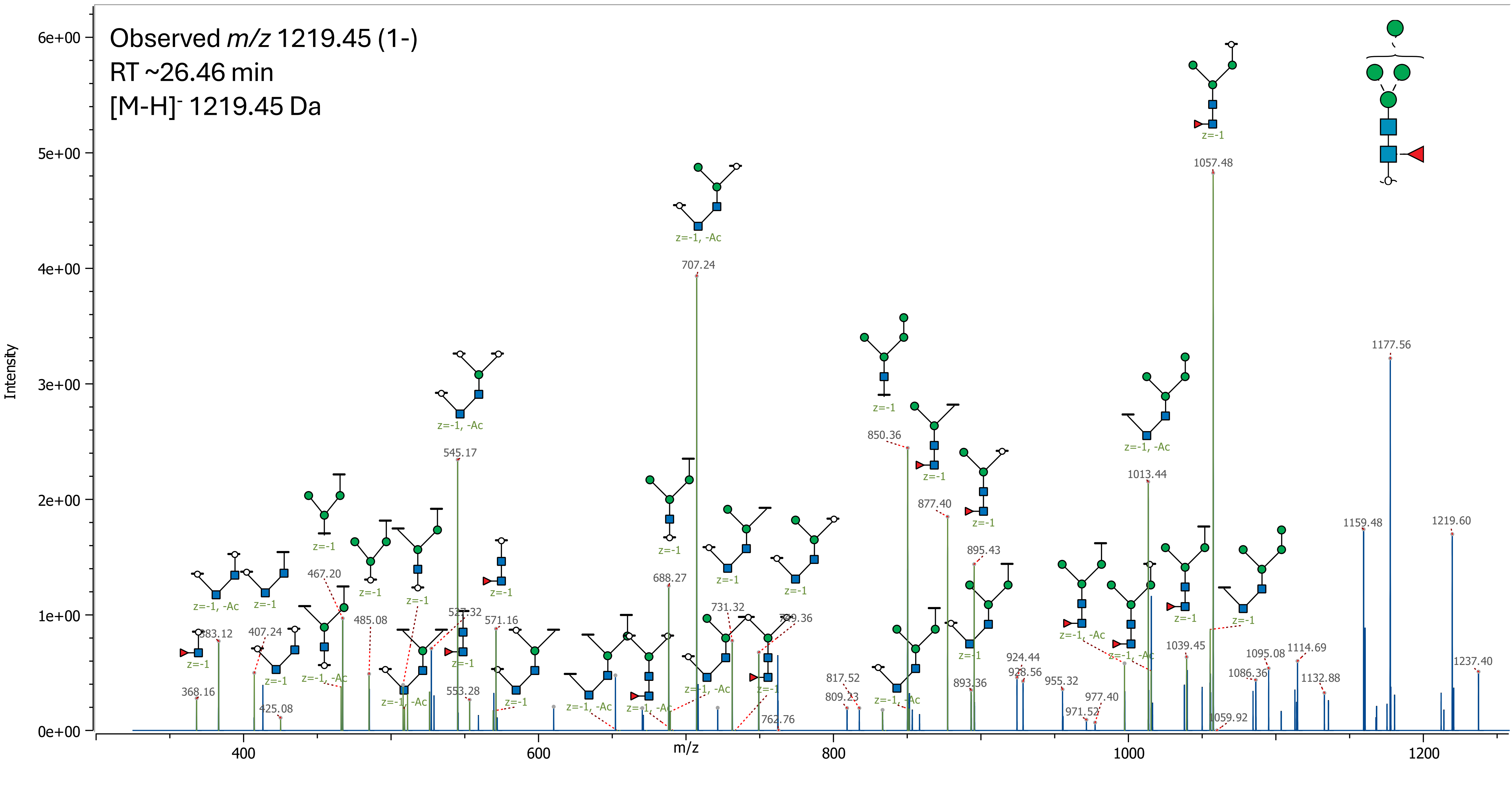

10 (Hex)2 + (Man)3(GlcNAc)2

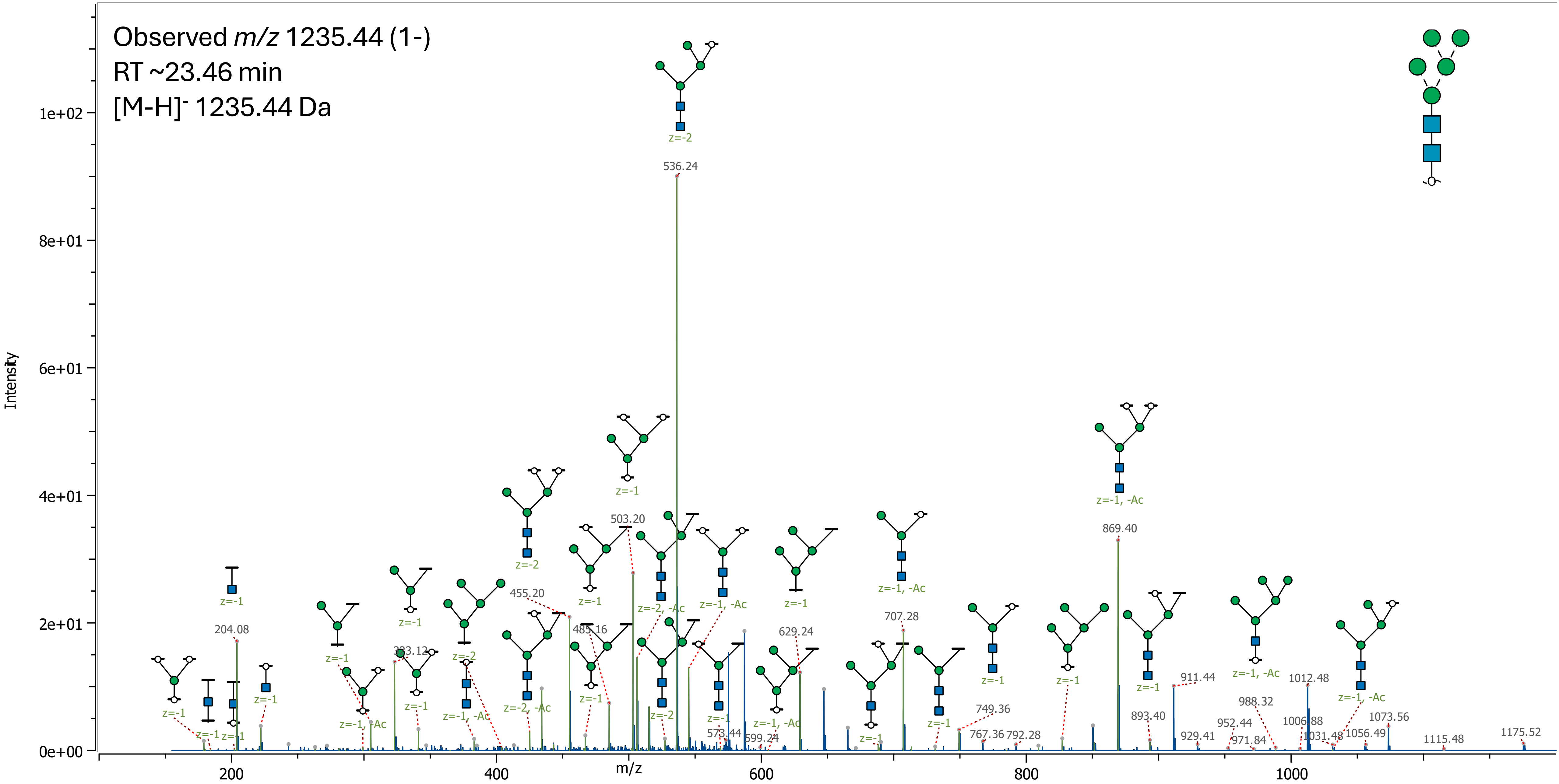

11 (Hex)2 (Deoxyhexose)1 + (Man)3(GlcNAc)2

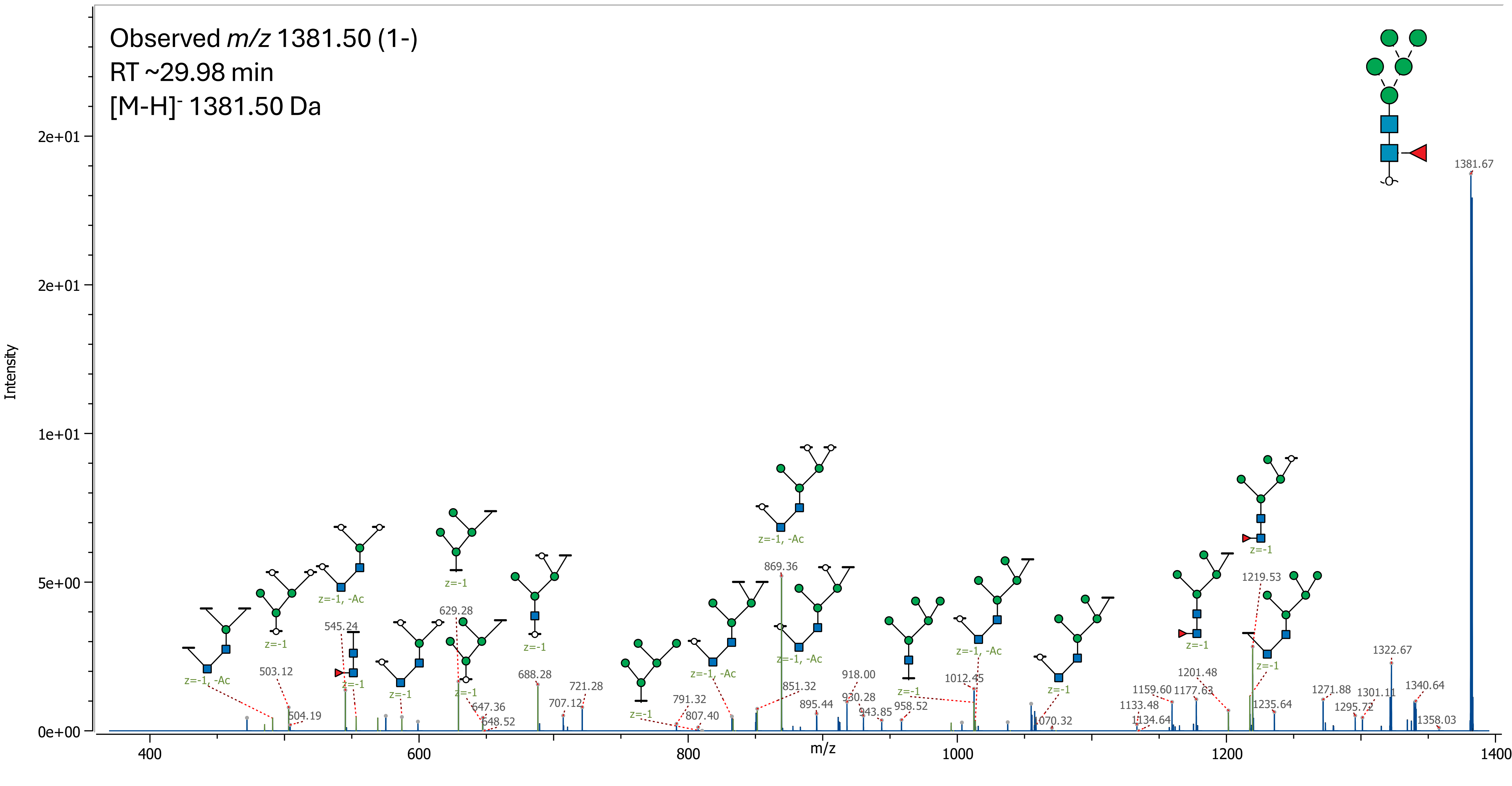

12 (Hex)3 + (Man)3(GlcNAc)2

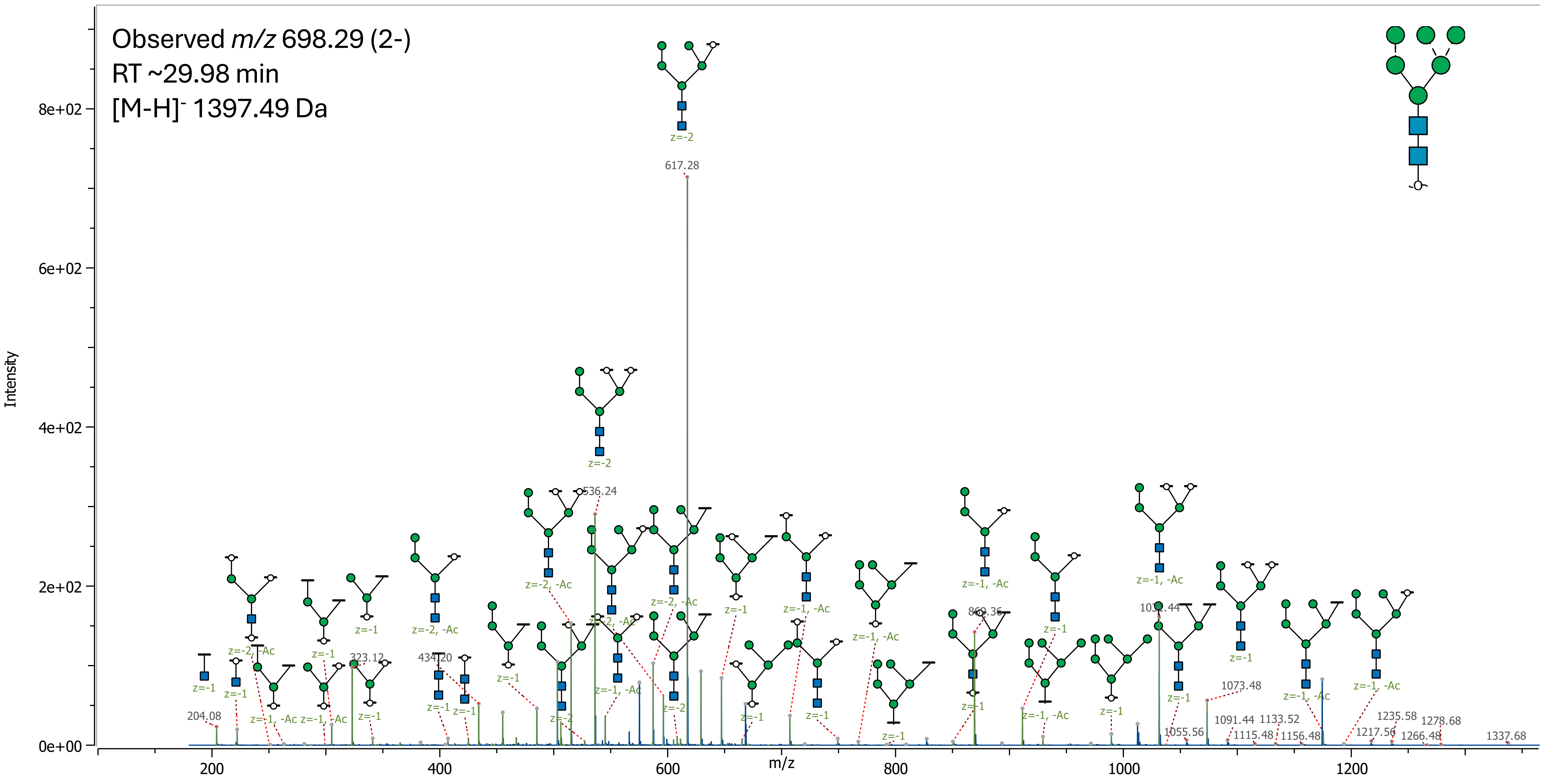

13 (Hex)4 + (Man)3(GlcNAc)2

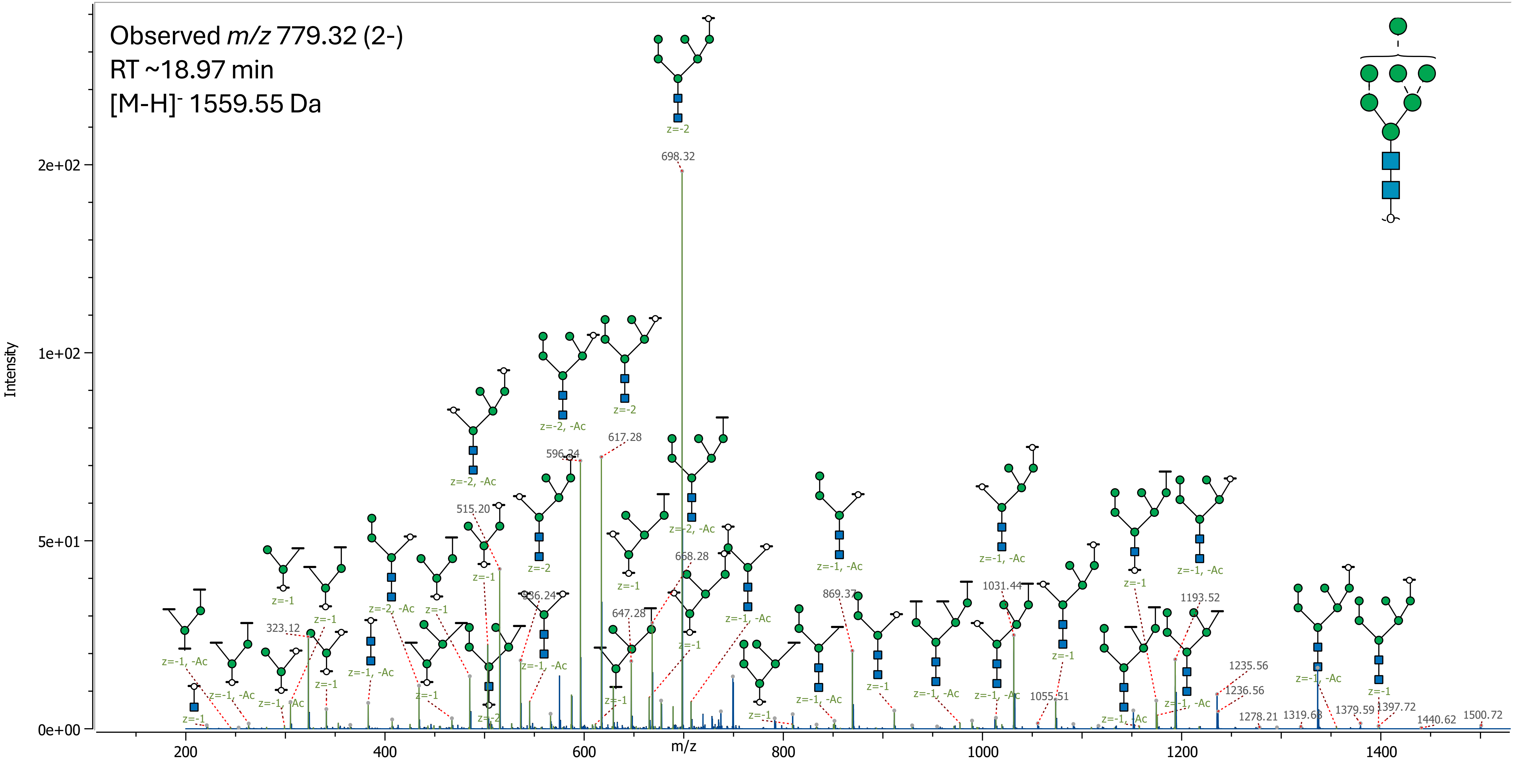

### 14 (Hex)5 + (Man)3(GlcNAc)2

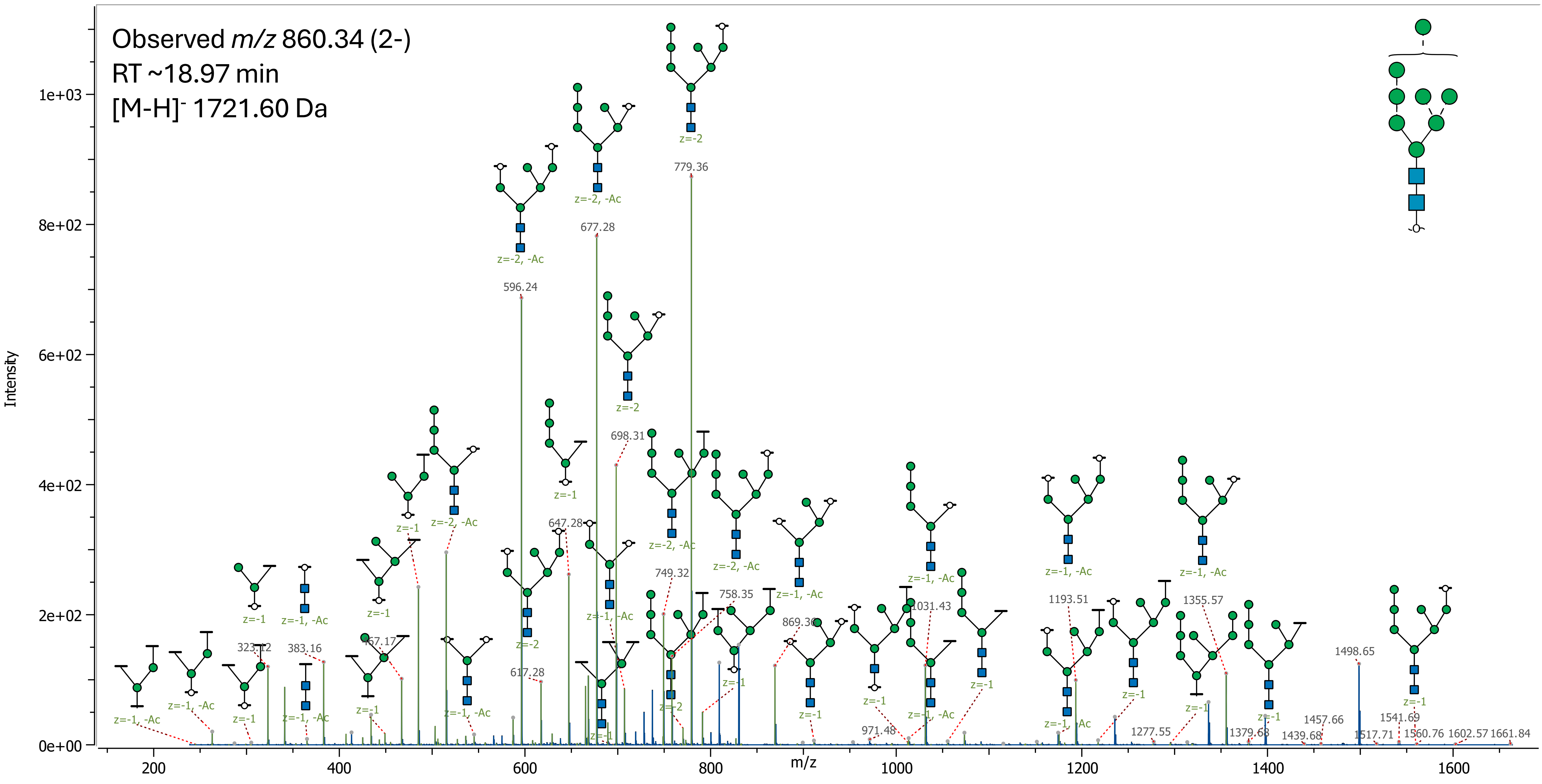

15 (Hex)6 + (Man)3(GlcNAc)2

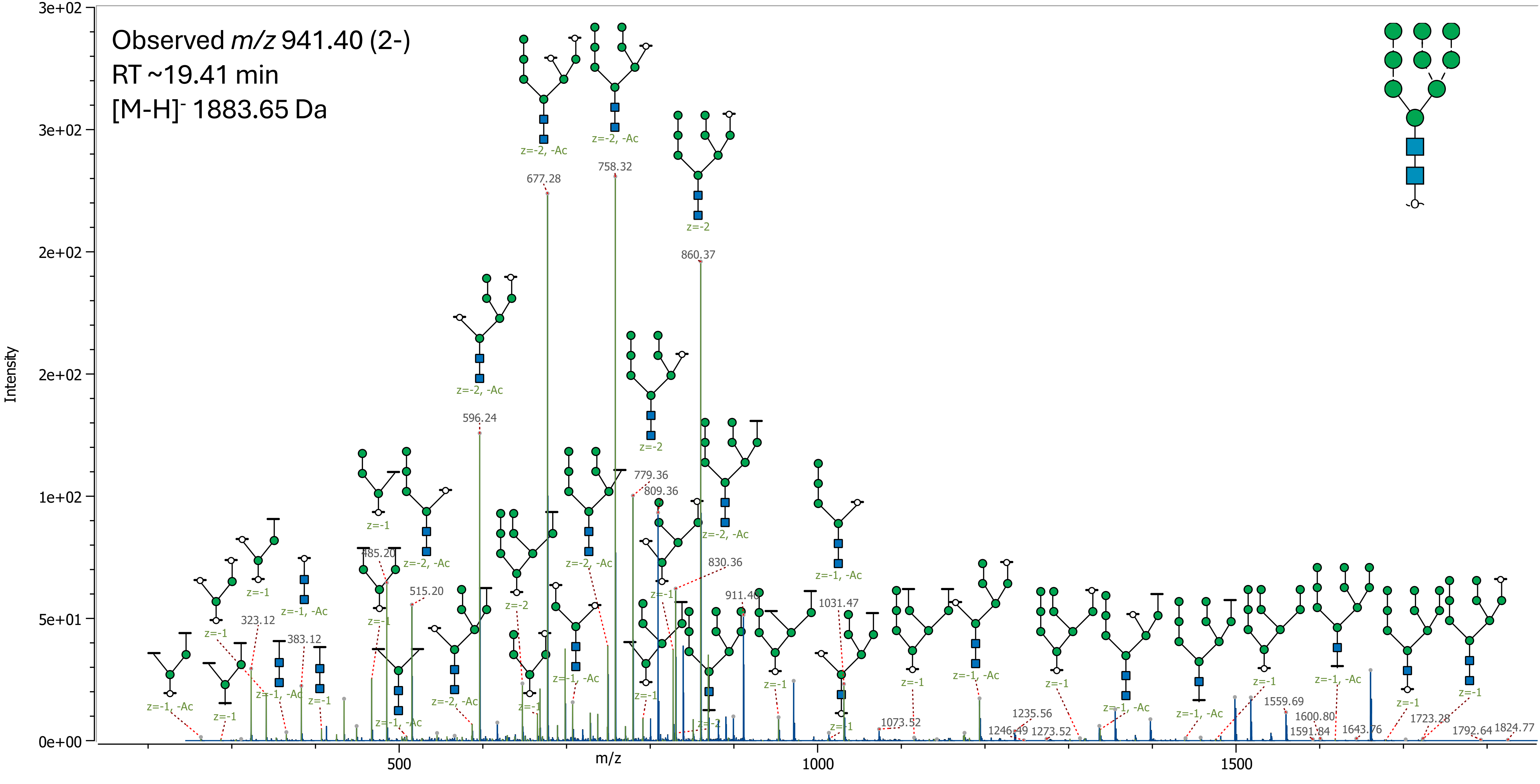

### 16 (Hex)7 + (Man)3(GlcNAc)2

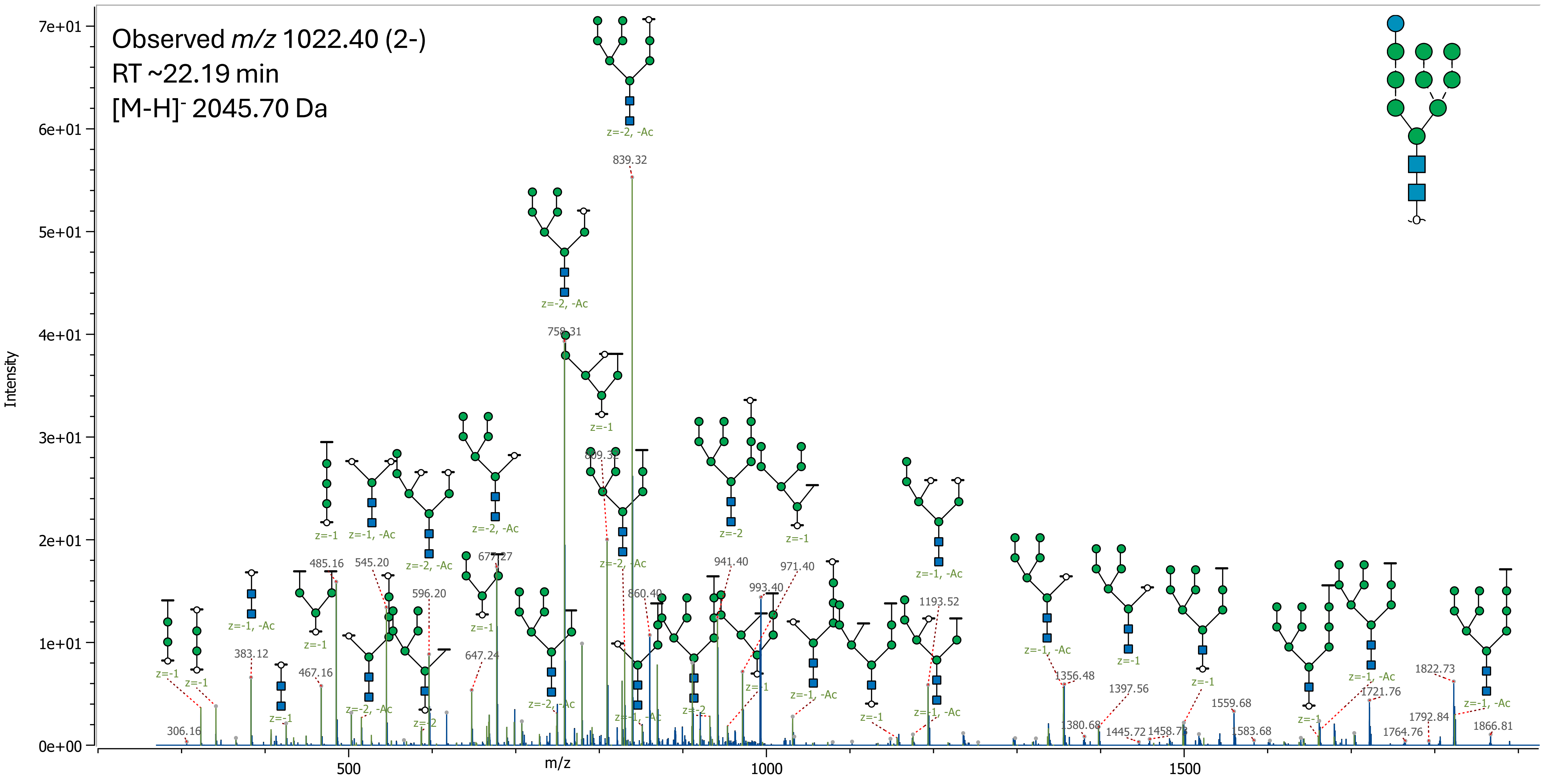

17 (HexNAc)1 + (Man)3(GlcNAc)2

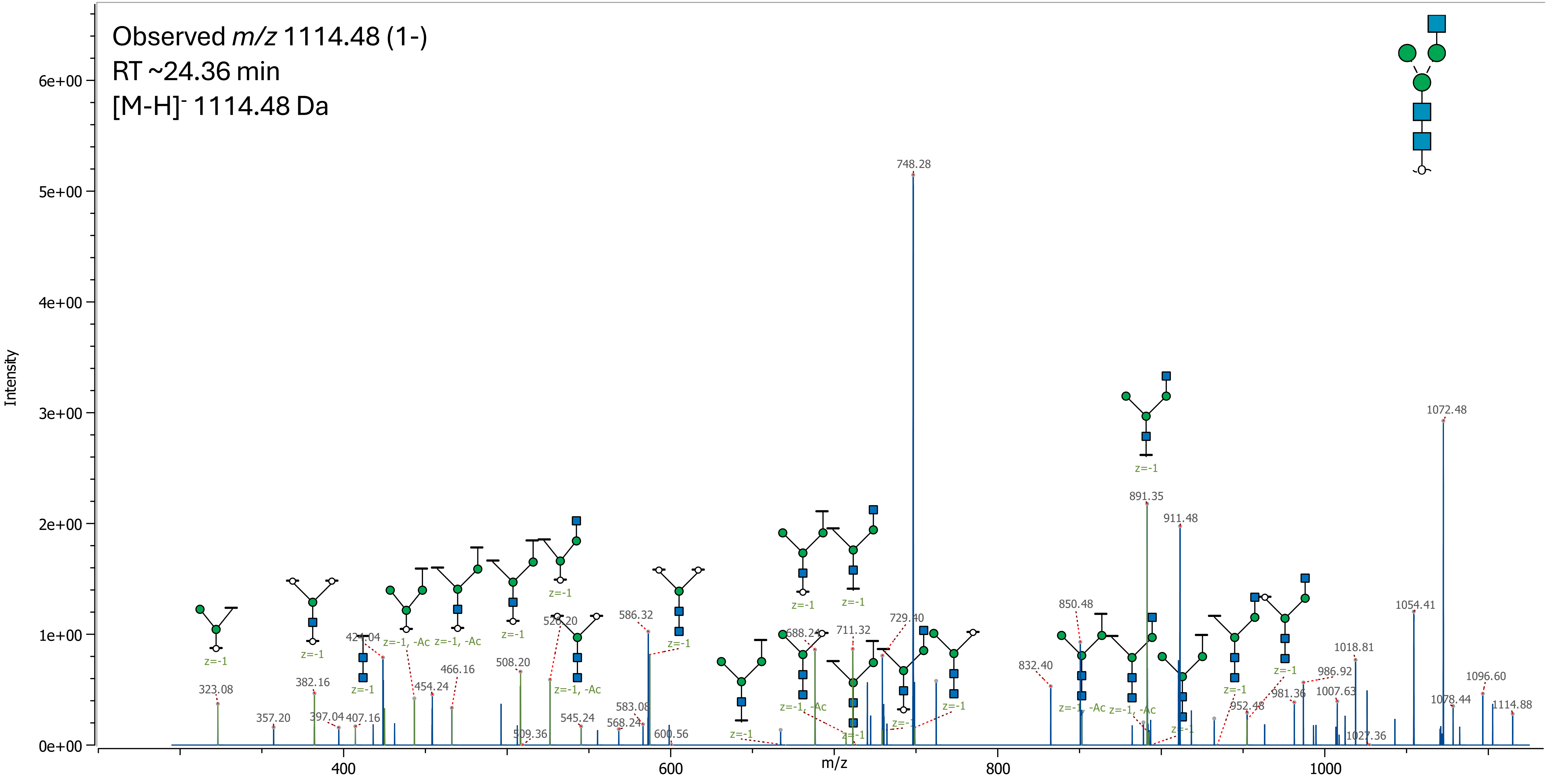

18 (HexNAc)1 (Deoxyhexose)1 + (Man)3(GlcNAc)2

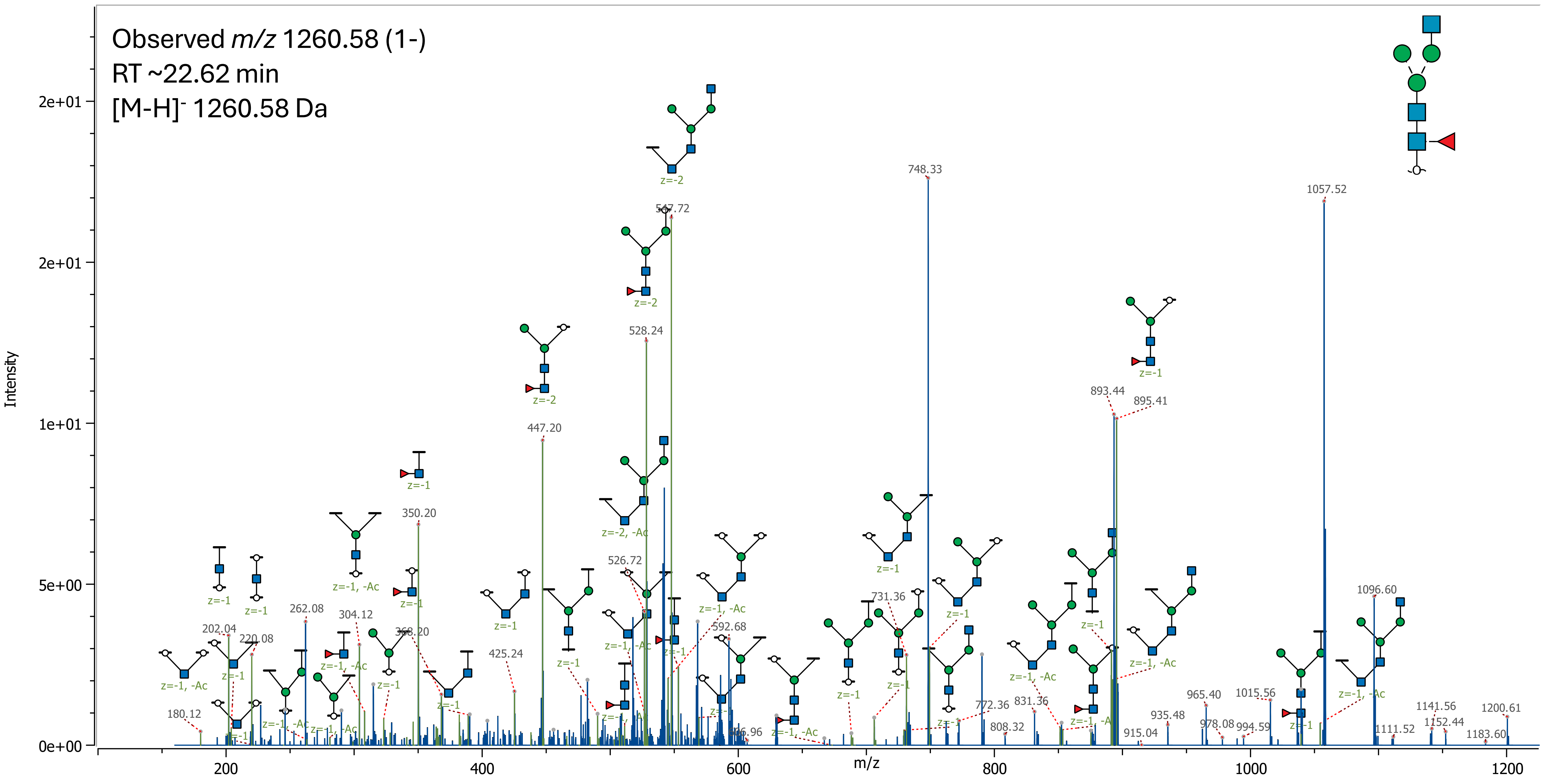

19 (Hex)1 (HexNAc)1 + (Man)3(GlcNAc)2

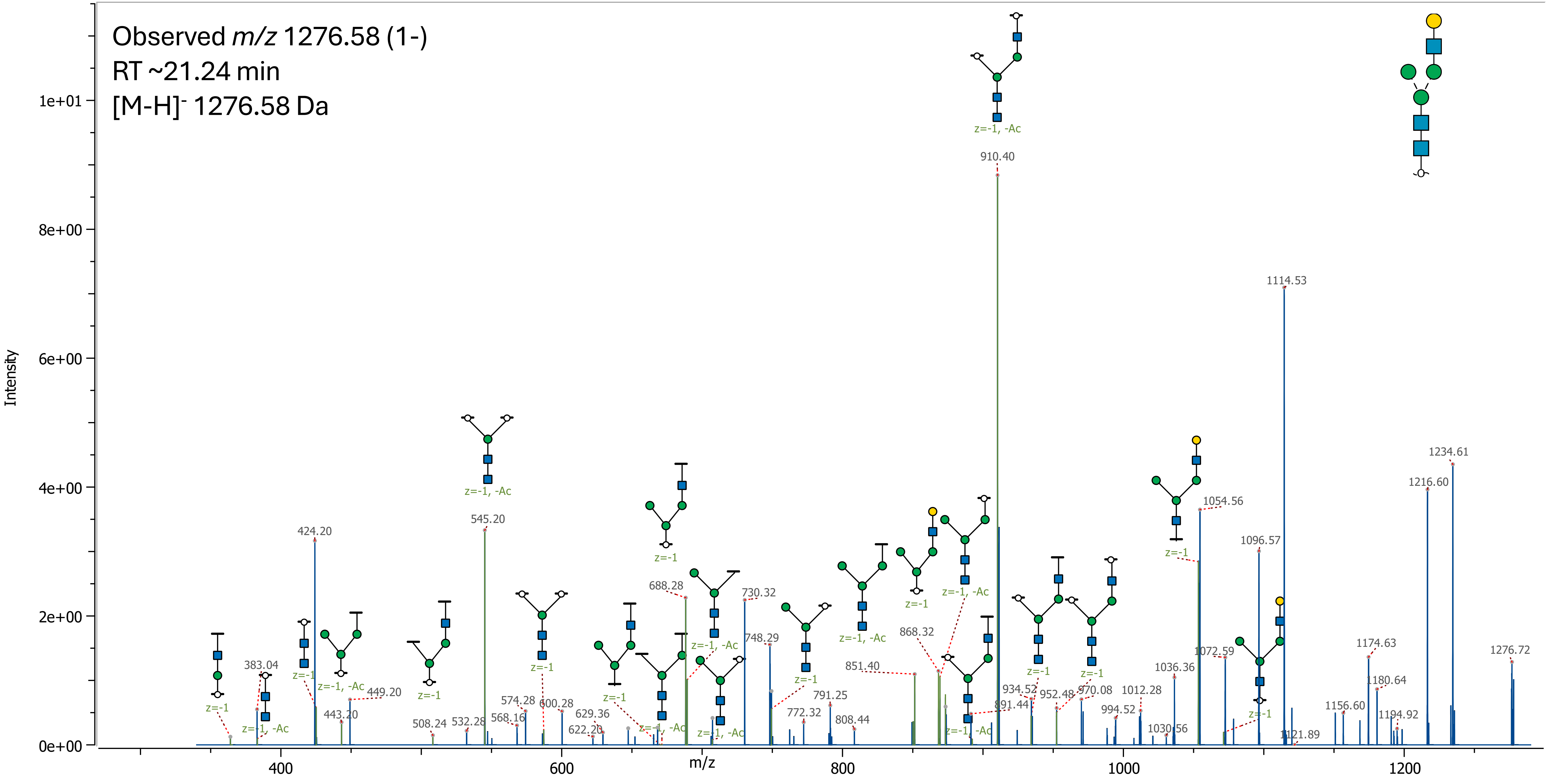

### 20 (HexNAc)2 + (Man)3(GlcNAc)2

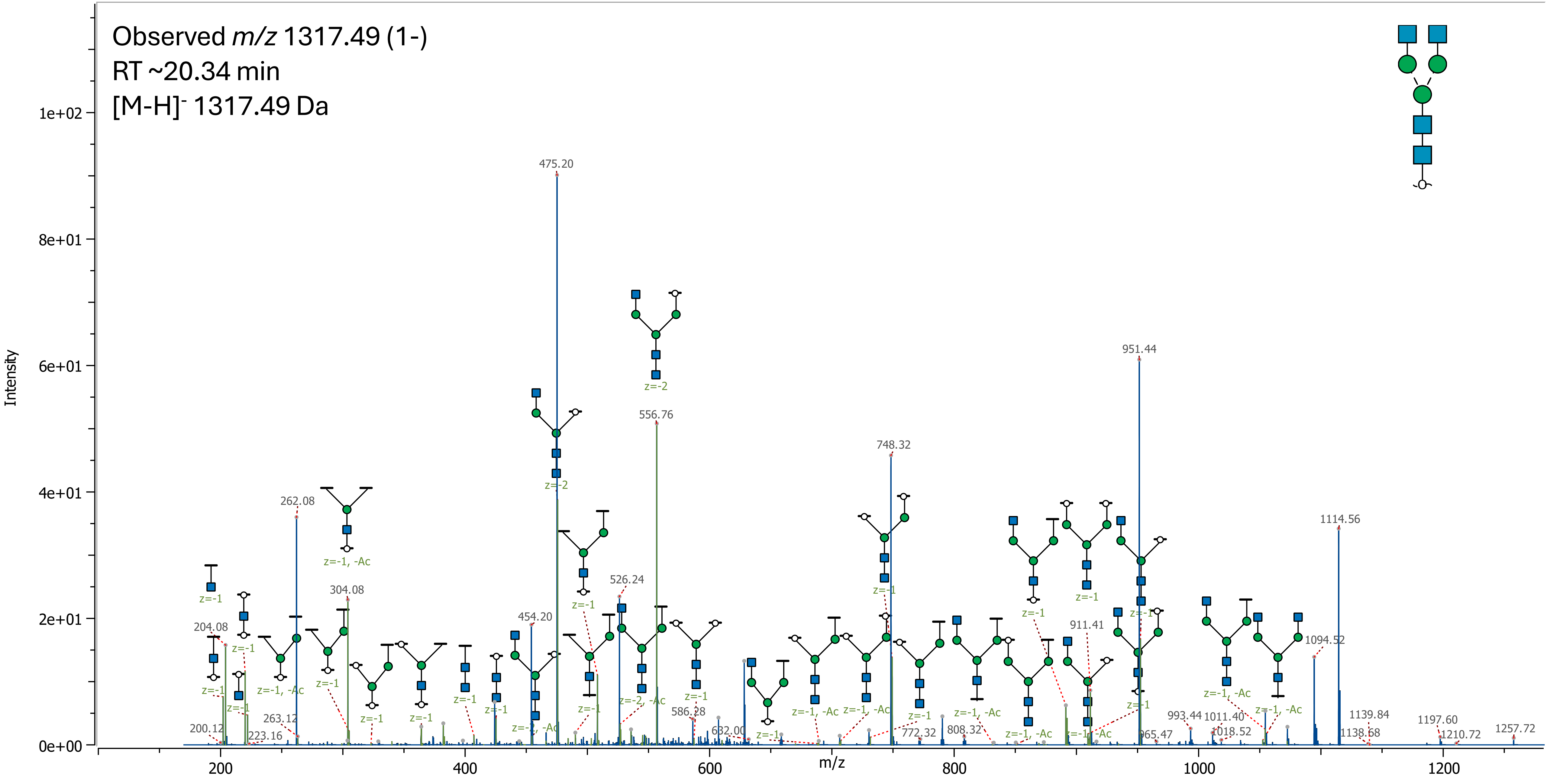

21 (Hex)1 (HexNAc)1 (Deoxyhexose)1 + (Man)3(GlcNAc)2

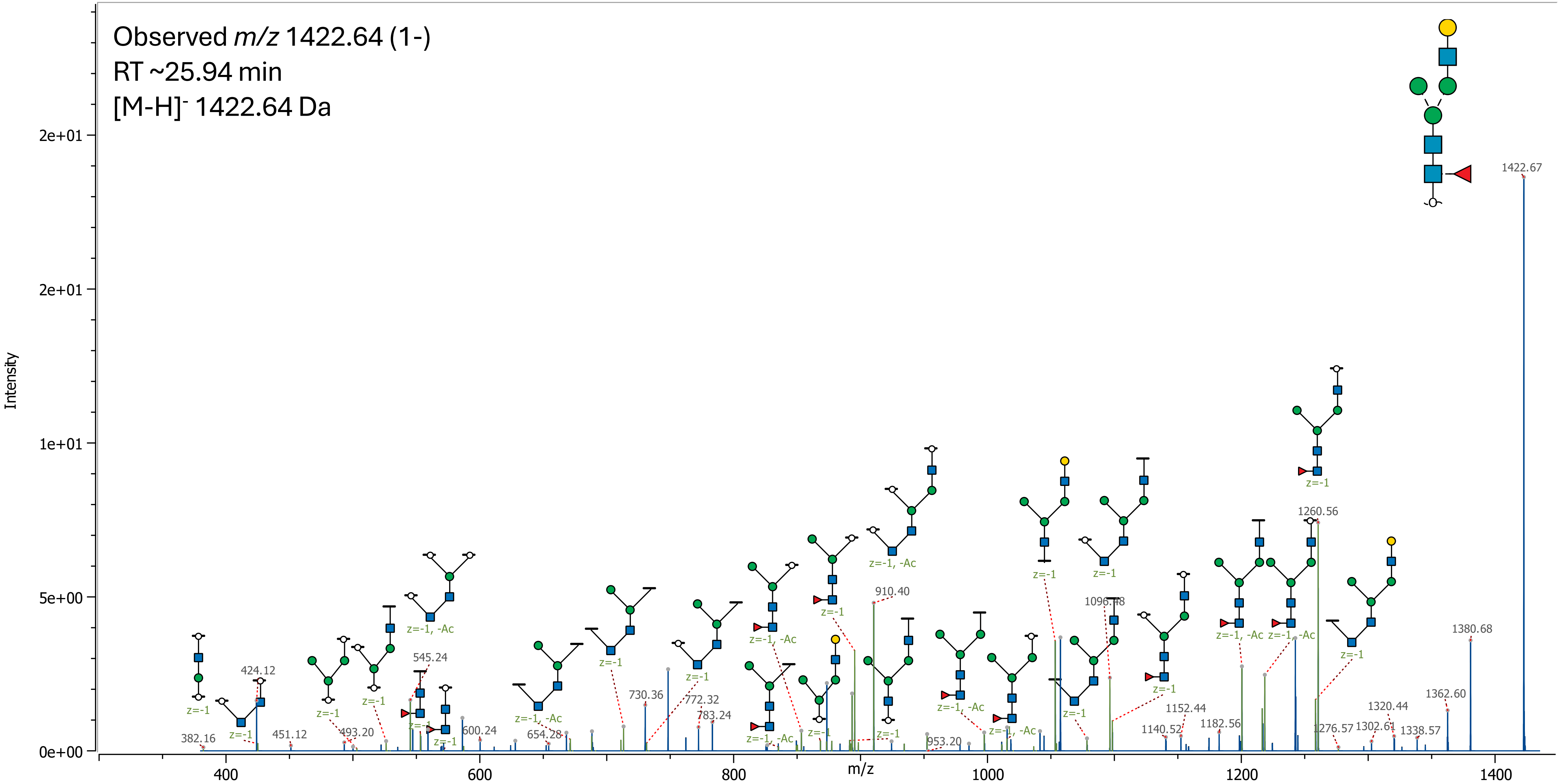

22 (Hex)2 (HexNAc)1 + (Man)3(GlcNAc)2

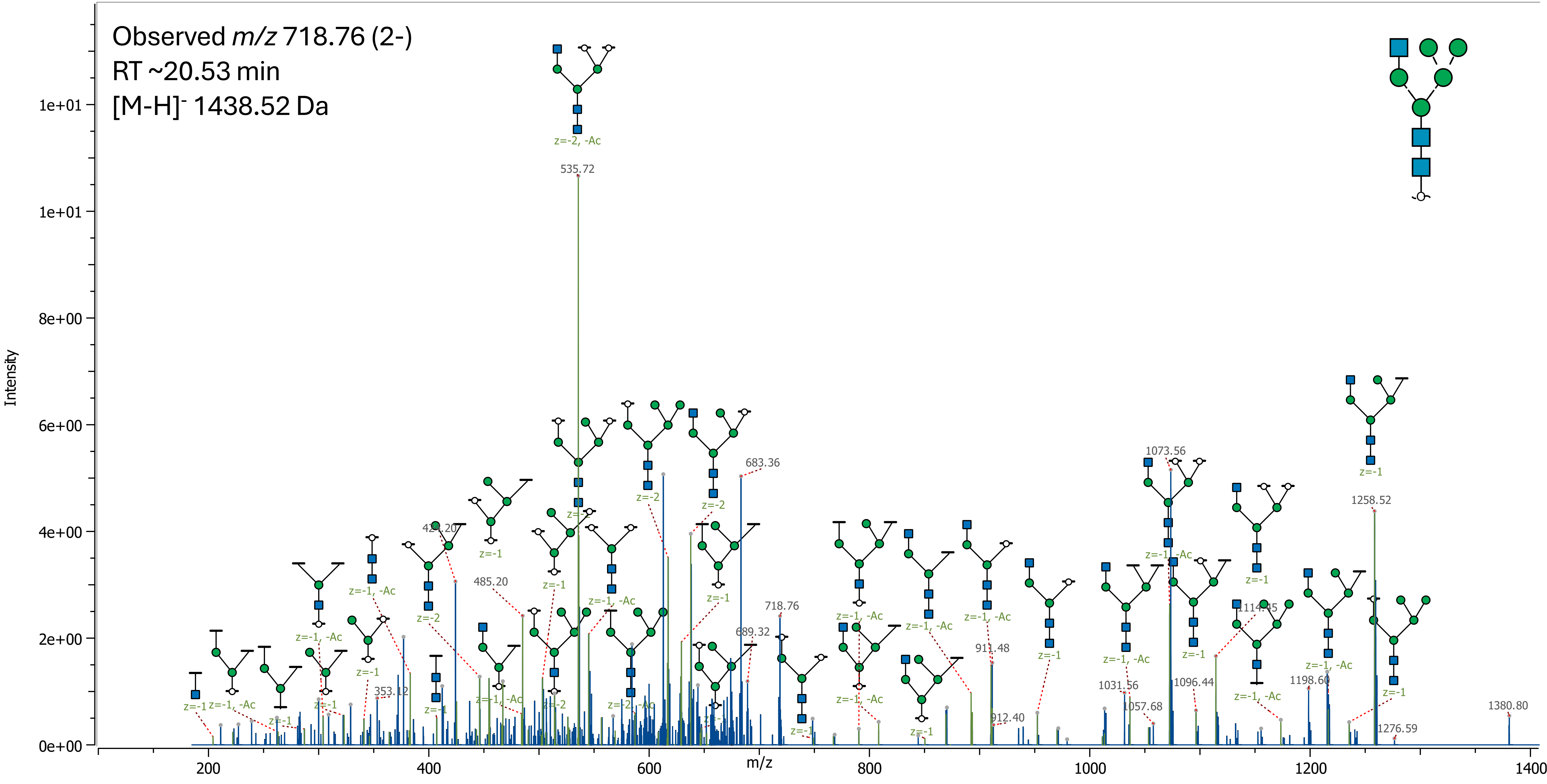

### 23 (HexNAc)2 (Deoxyhexose)1 + (Man)3(GlcNAc)2

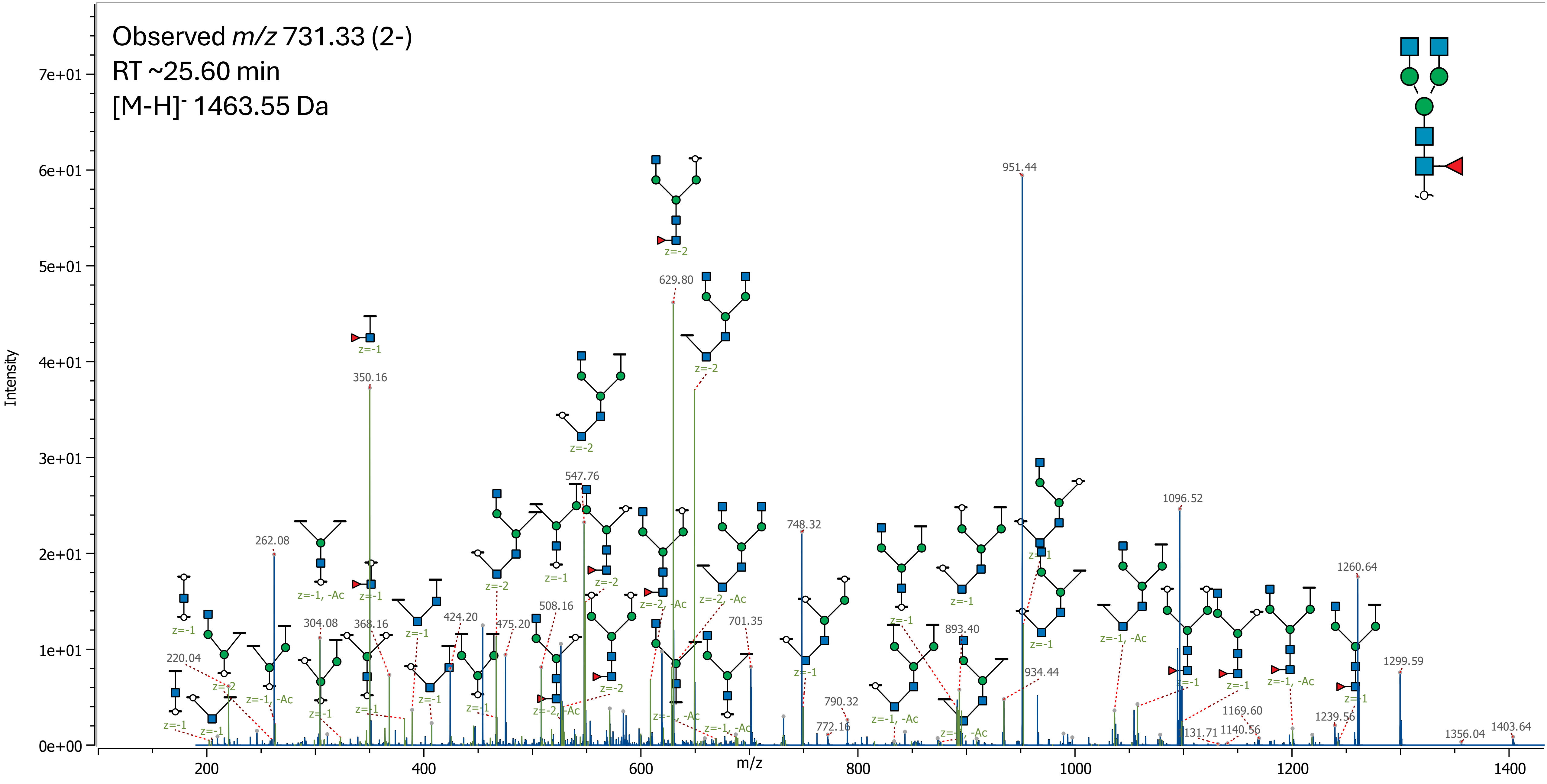

### 24 (Hex)1 (HexNAc)2 + (Man)3(GlcNAc)2

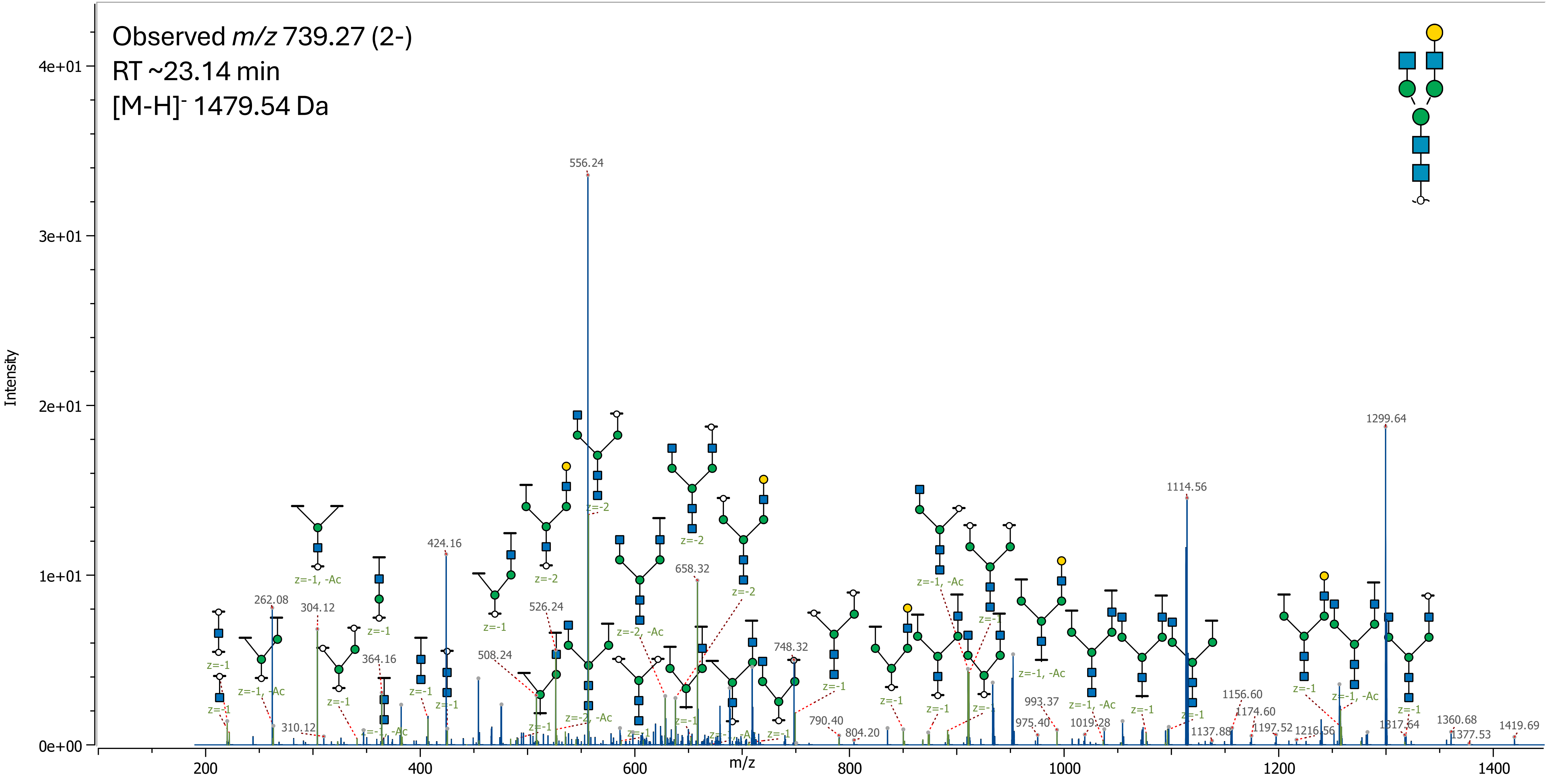

25 (HexNAc)3 + (Man)3(GlcNAc)2

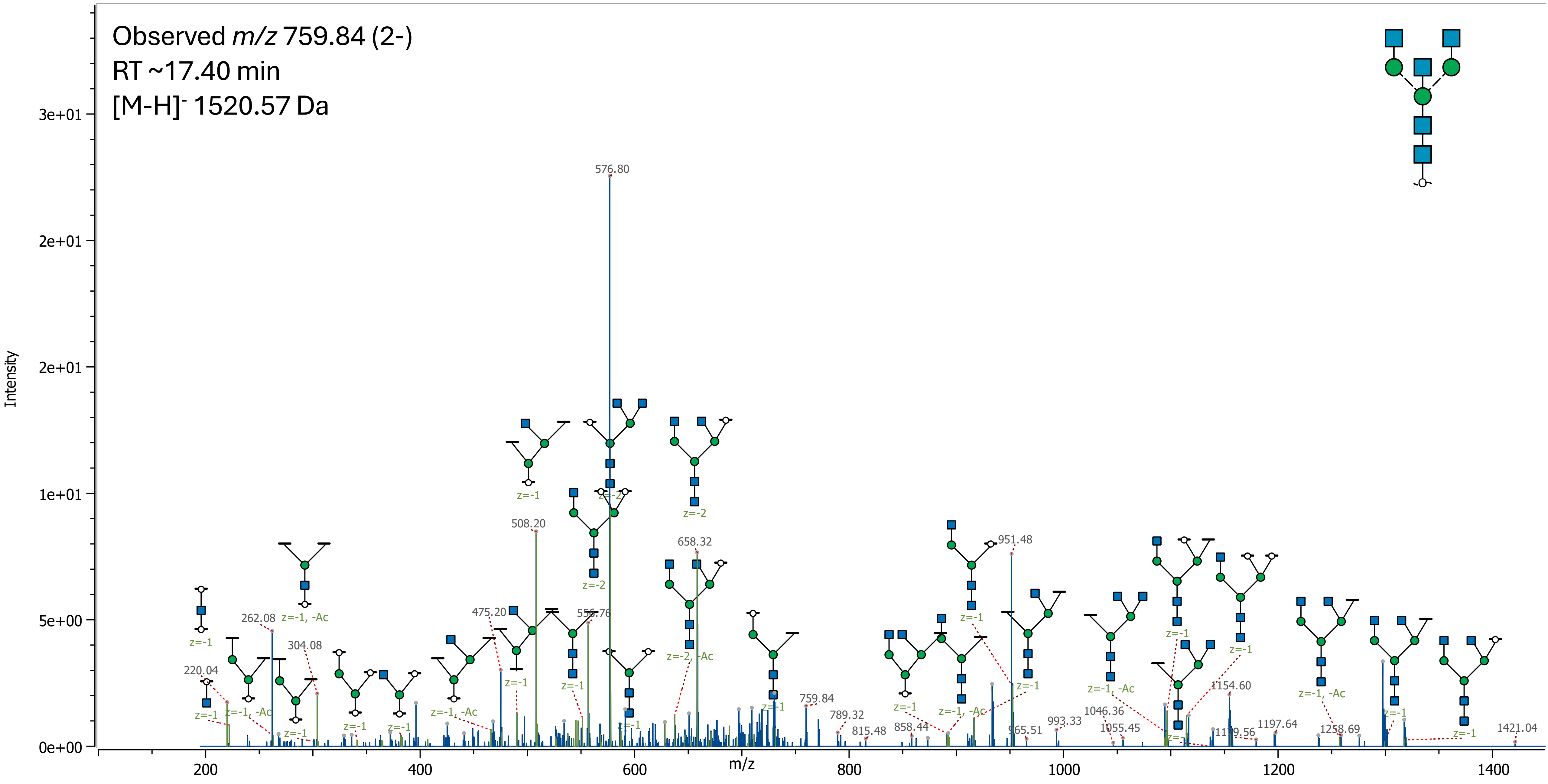

### 26 (Hex)1 (HexNAc)1 (NeuAc)1 + (Man)3(GlcNAc)2

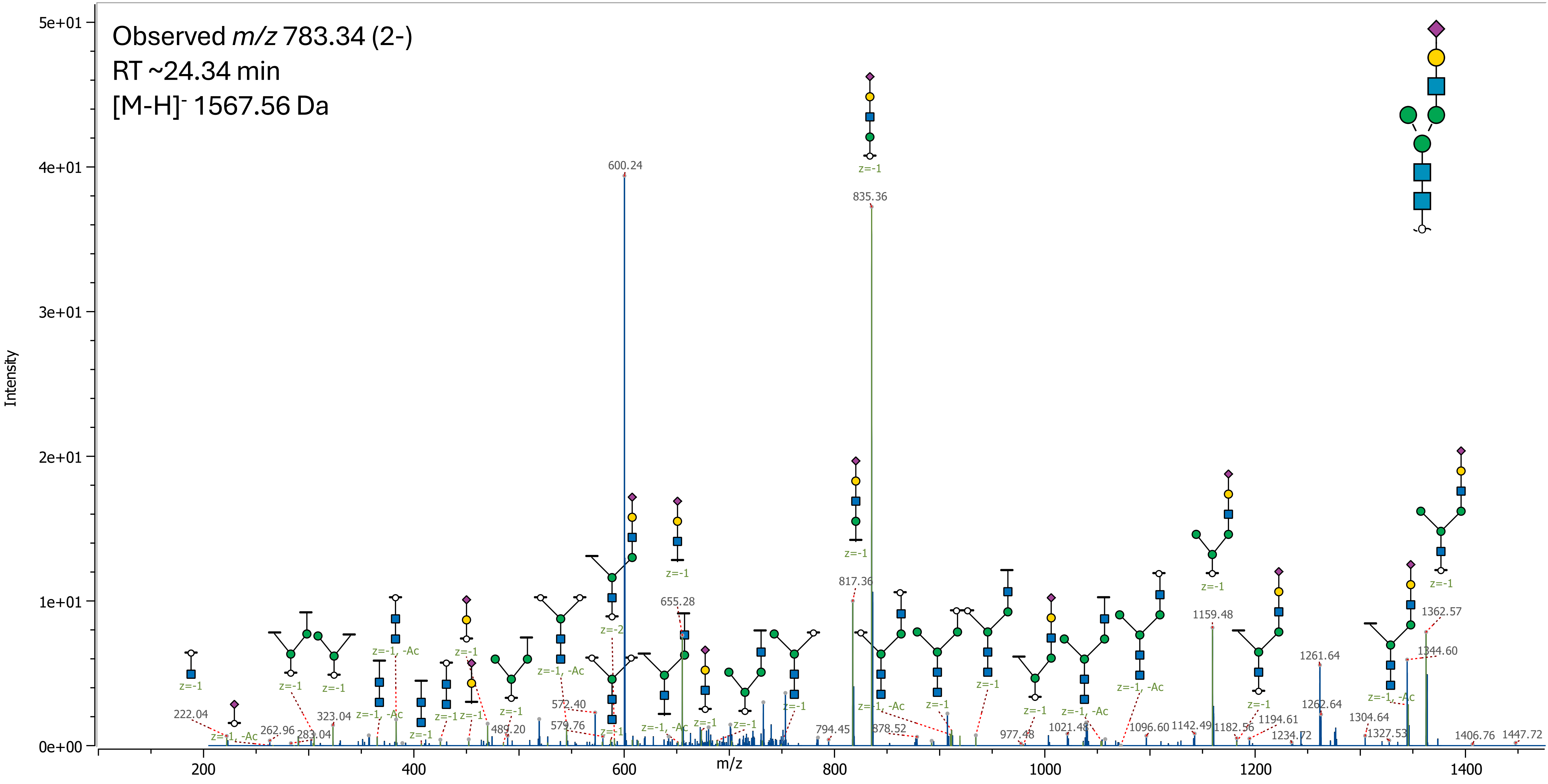

27 (Hex)2 (HexNAc)1 (Deoxyhexose)1 + (Man)3(GlcNAc)2

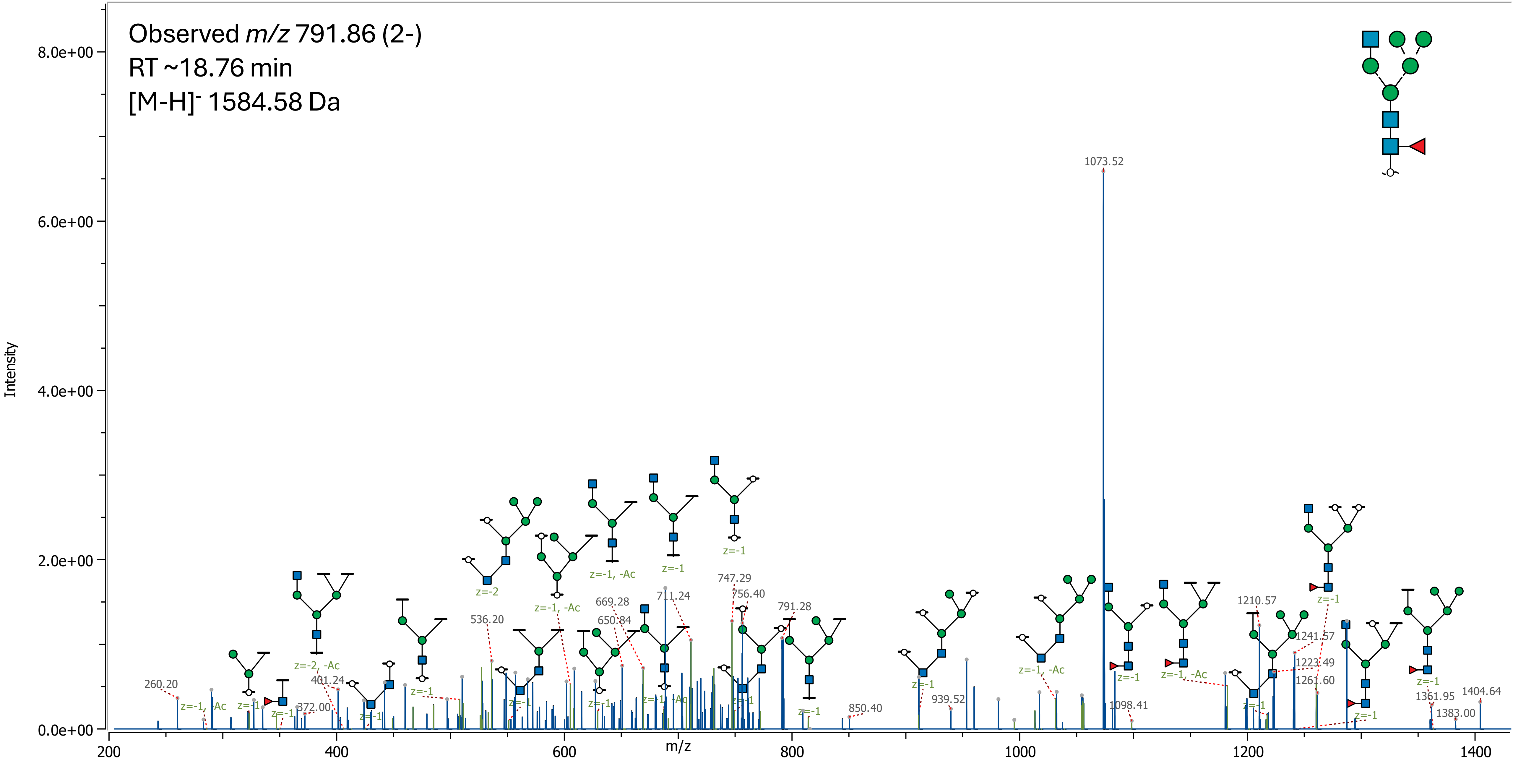

### 28 (Hex)3 (HexNAc)1 + (Man)3(GlcNAc)2

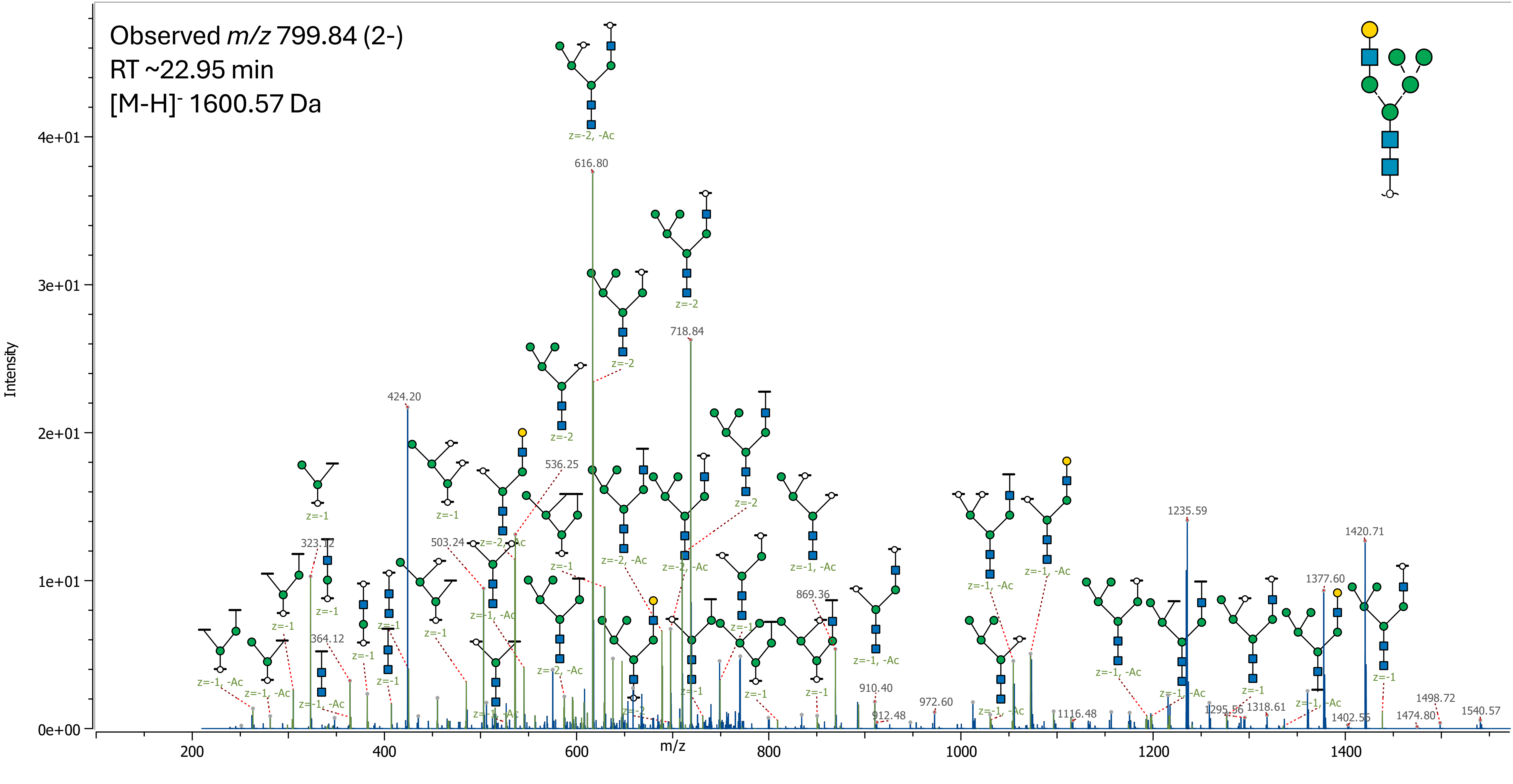

29 (HexNAc)2 (Deoxyhexose)2 + (Man)3(GlcNAc)2

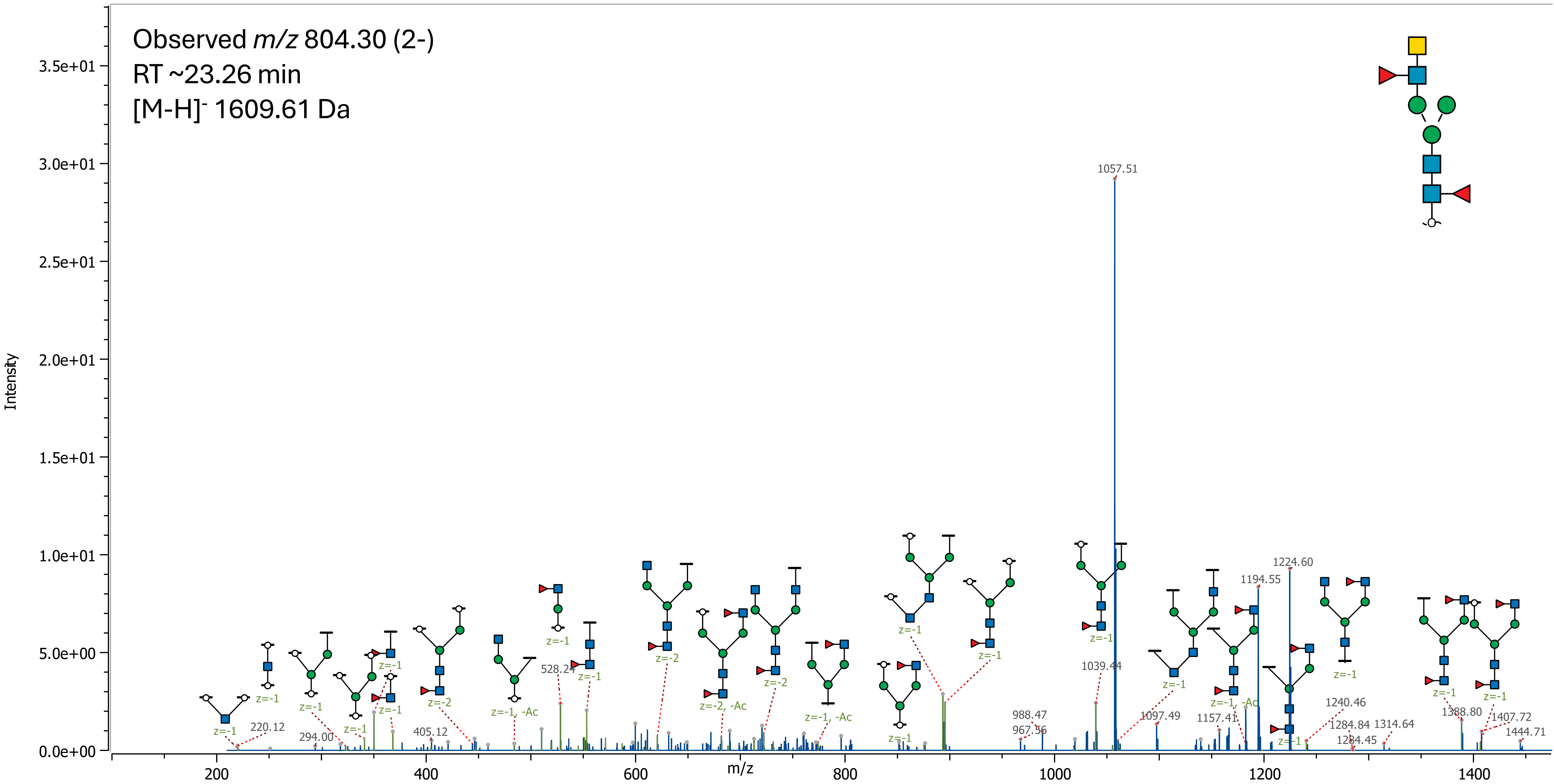

30 (Hex)1 (HexNAc)2 (Deoxyhexose)1 + (Man)3(GlcNAc)2

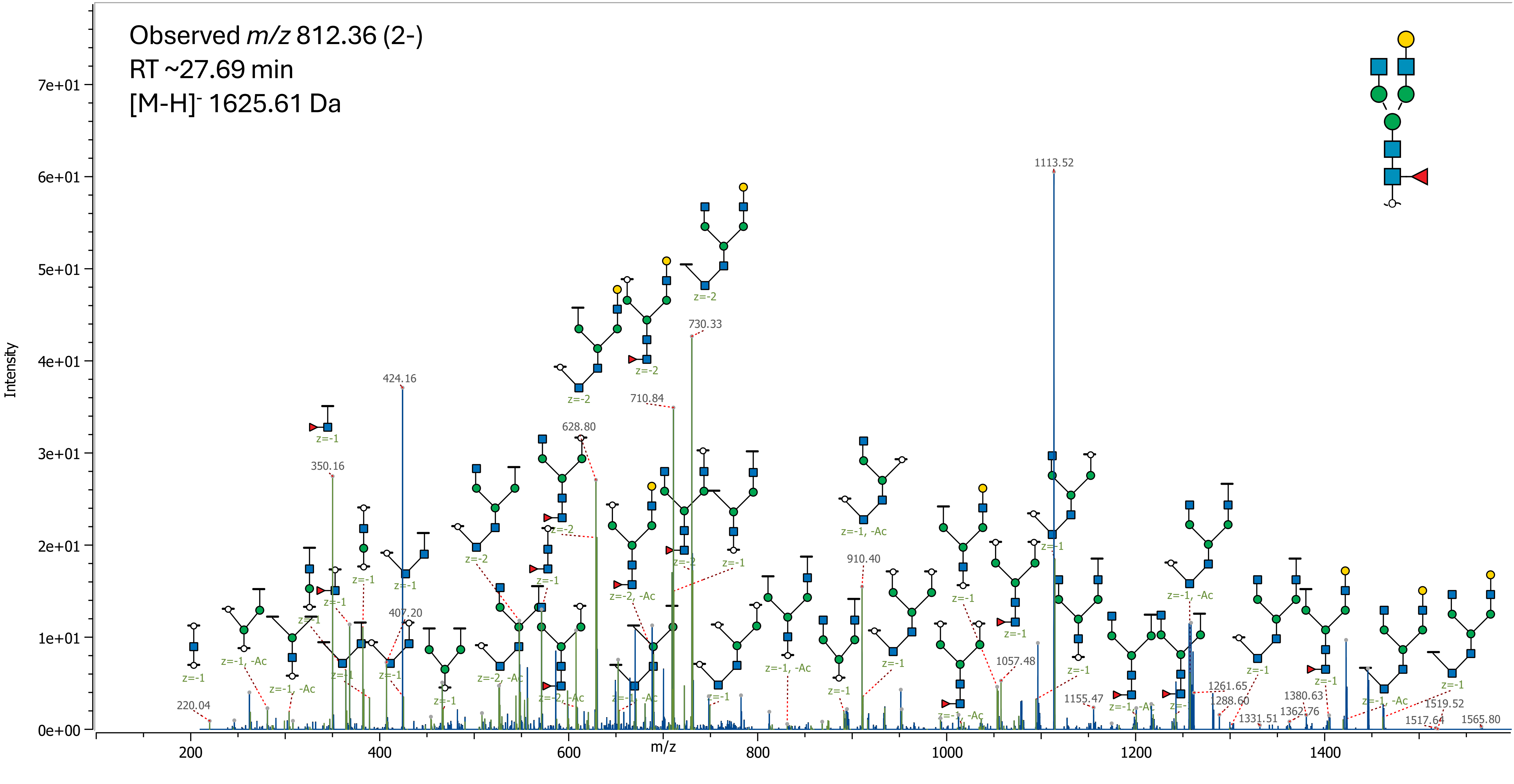

31 (Hex)2 (HexNAc)2 + (Man)3(GlcNAc)2

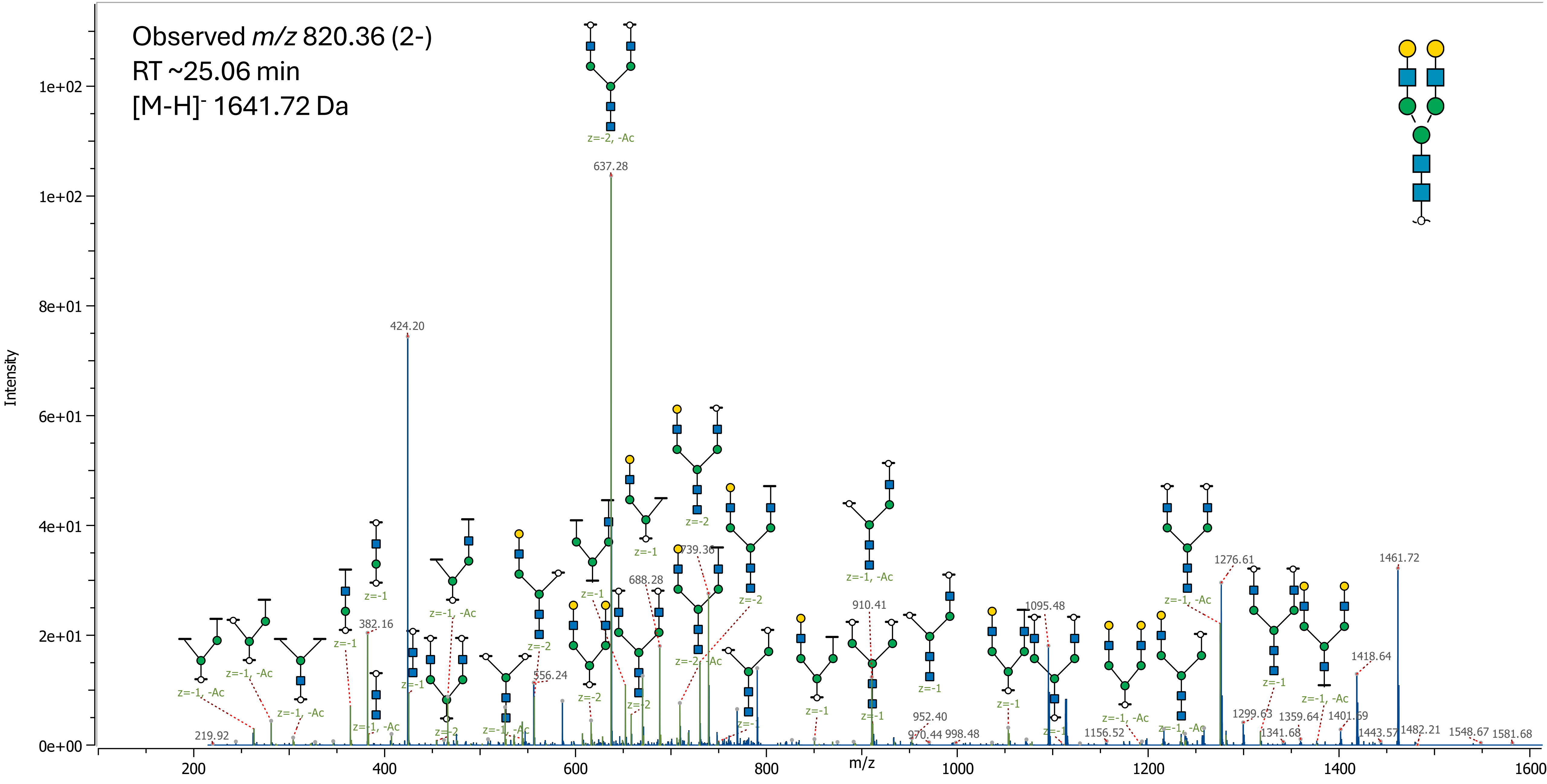

32 (HexNAc)3 (Deoxyhexose)1 + (Man)3(GlcNAc)2

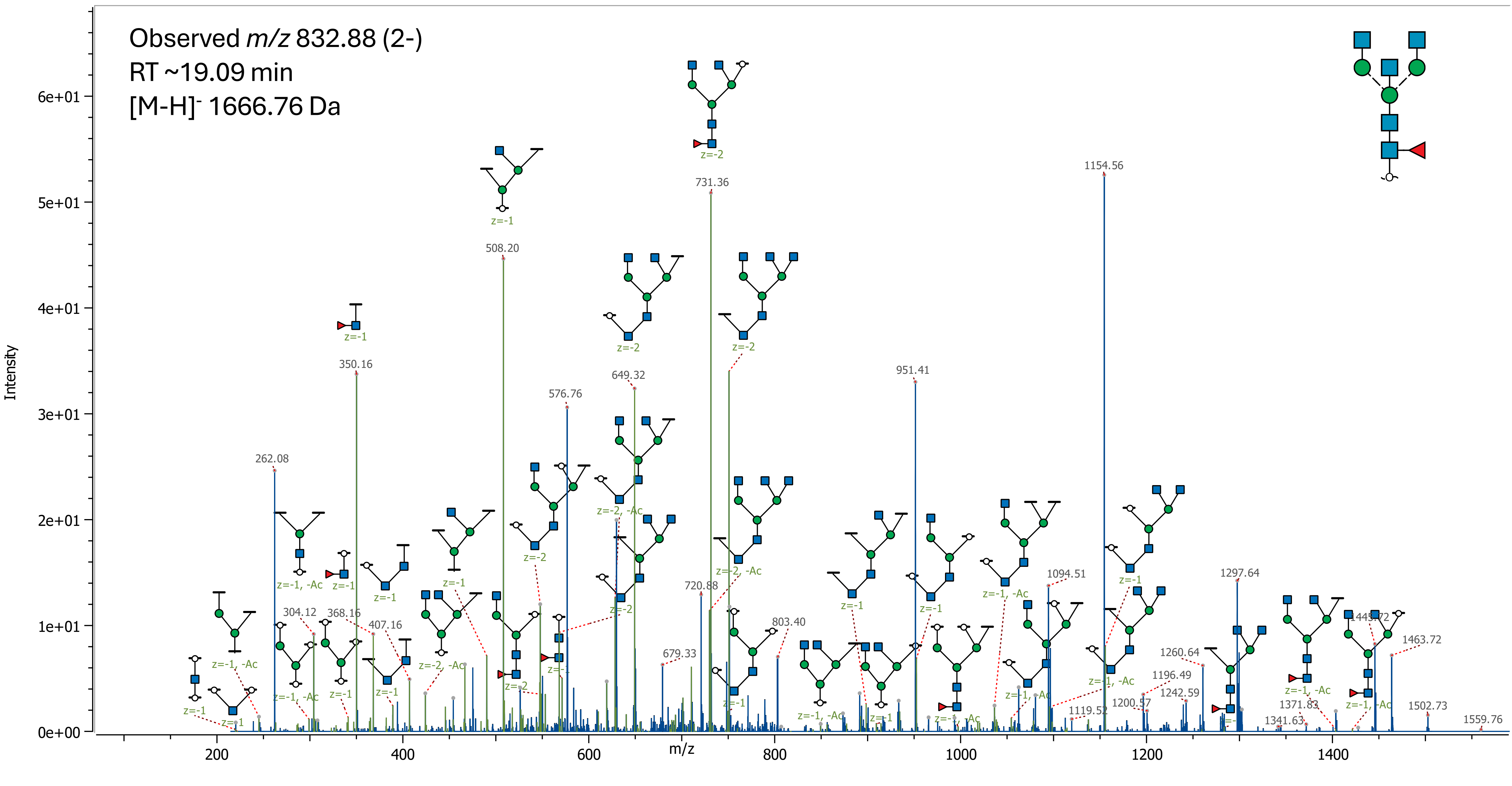

33 (Hex)1 (HexNAc)3 + (Man)3(GlcNAc)2

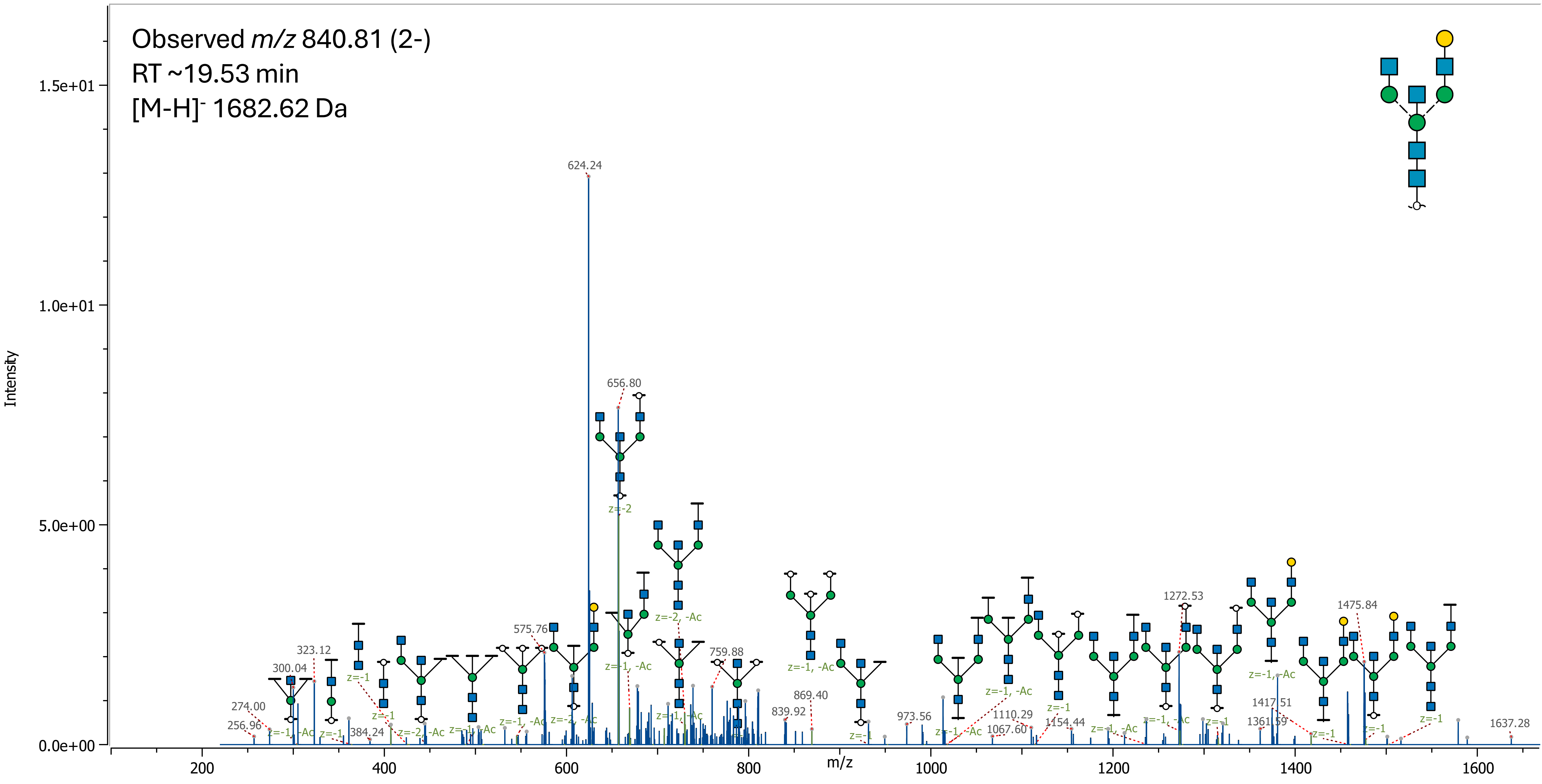

### 34 (Hex)1 (HexNAc)1 (Deoxyhexose)1 (NeuAc)1 + (Man)3(GlcNAc)2

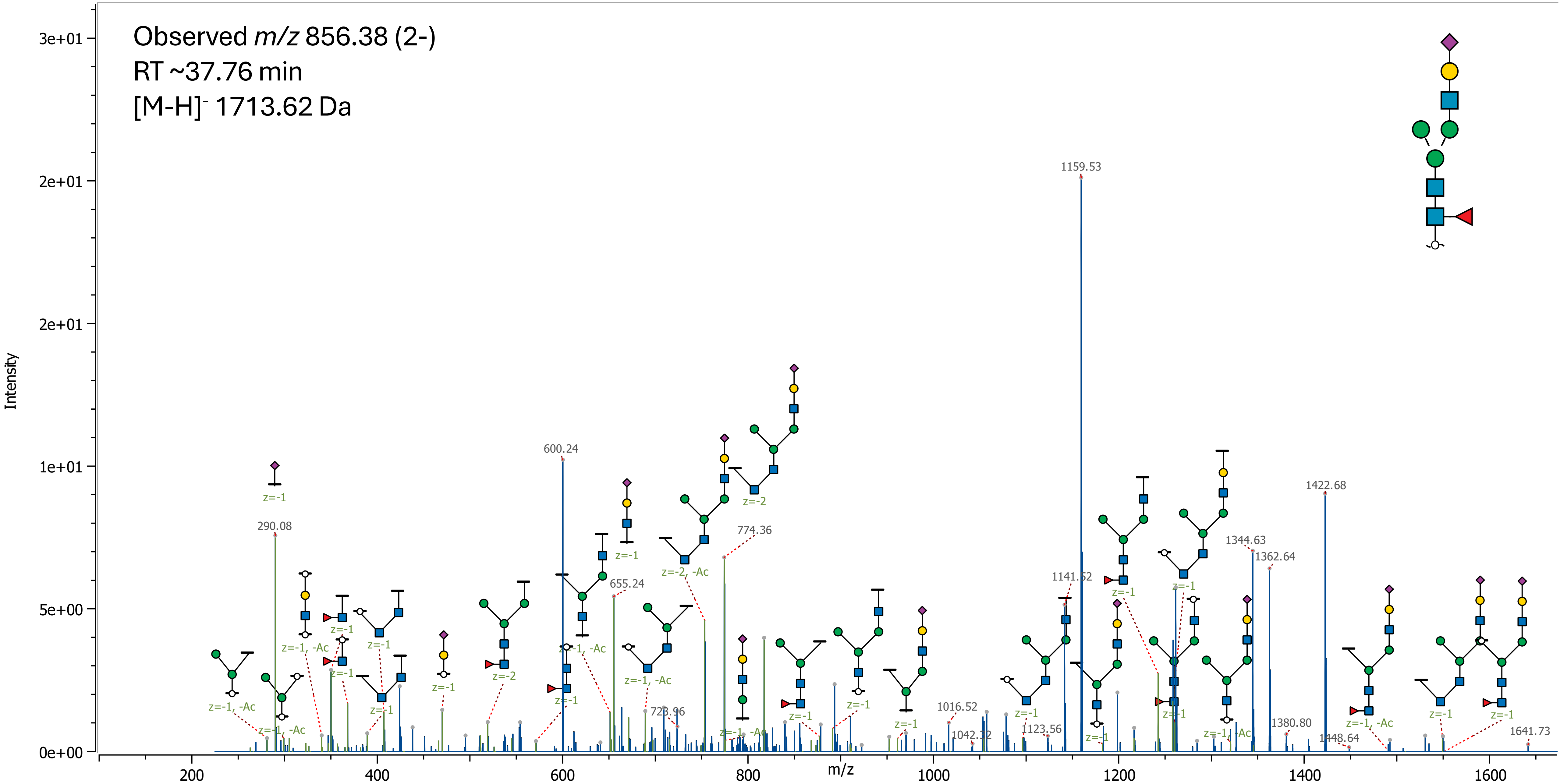

35 (HexNAc)4 + (Man)3(GlcNAc)2

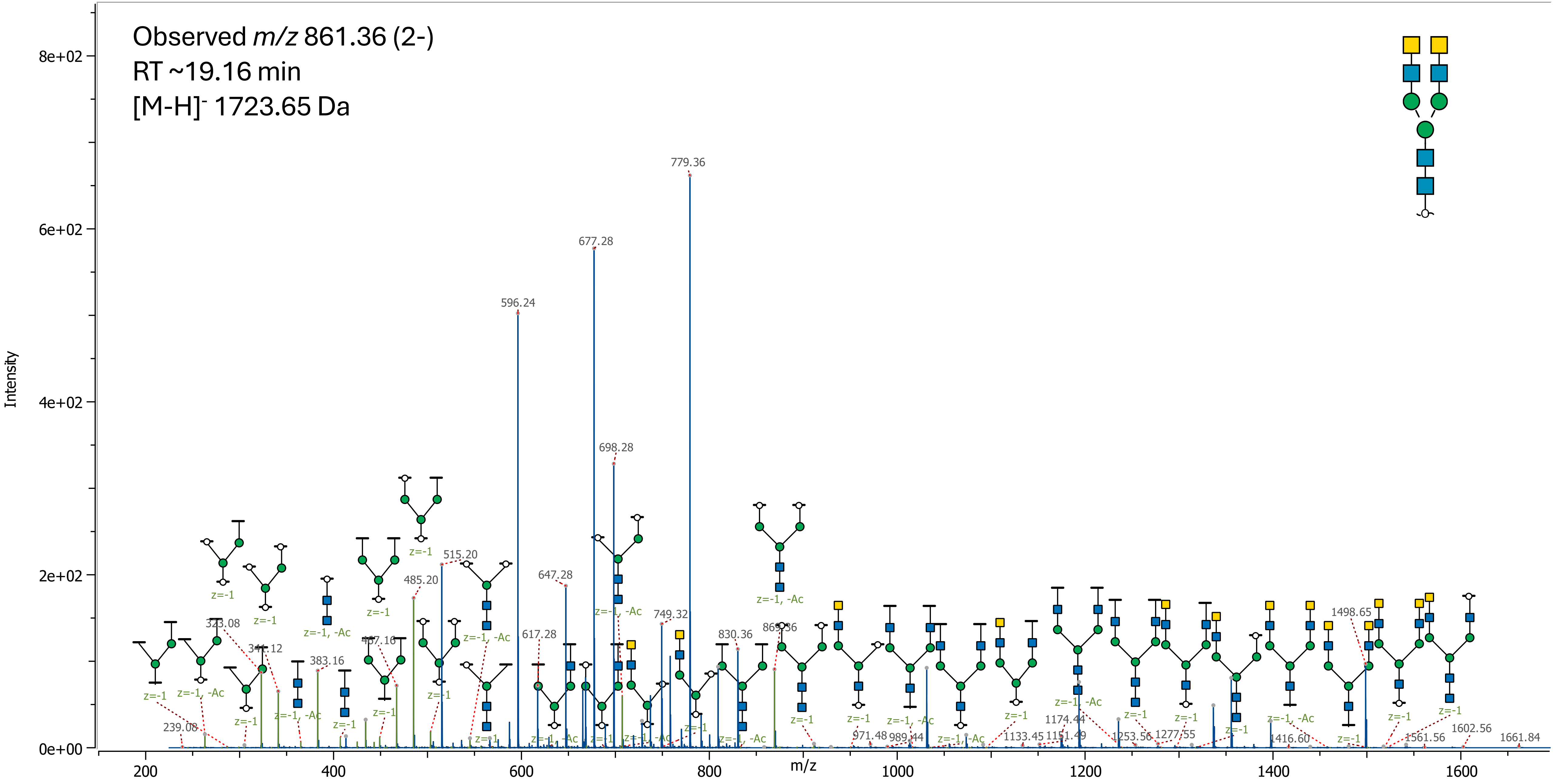

36 (Hex)2 (HexNAc)1 (NeuAc)1 + (Man)3(GlcNAc)2

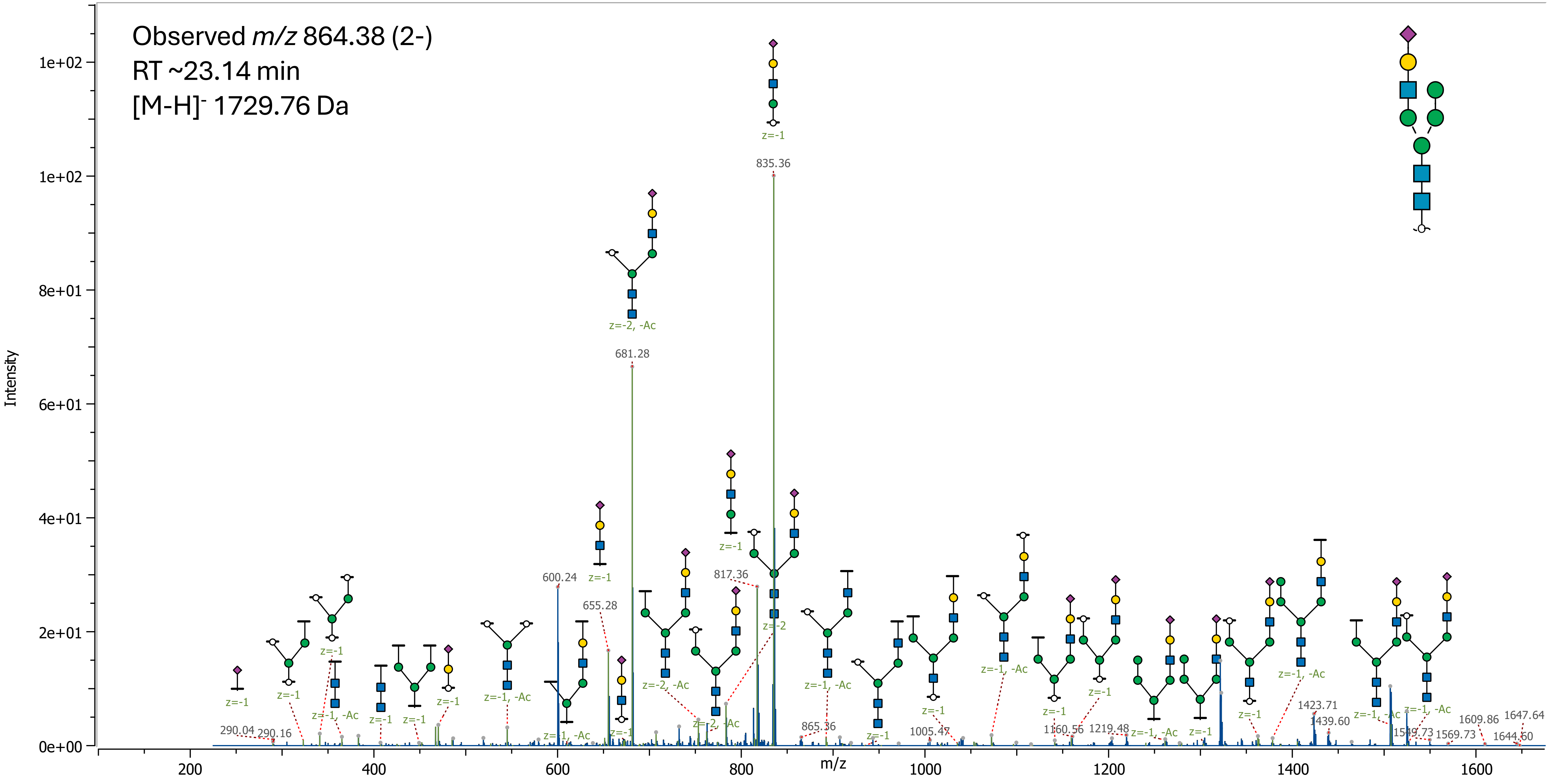

### 37 (Hex)3 (HexNAc)1 (Deoxyhexose)1 + (Man)3(GlcNAc)2

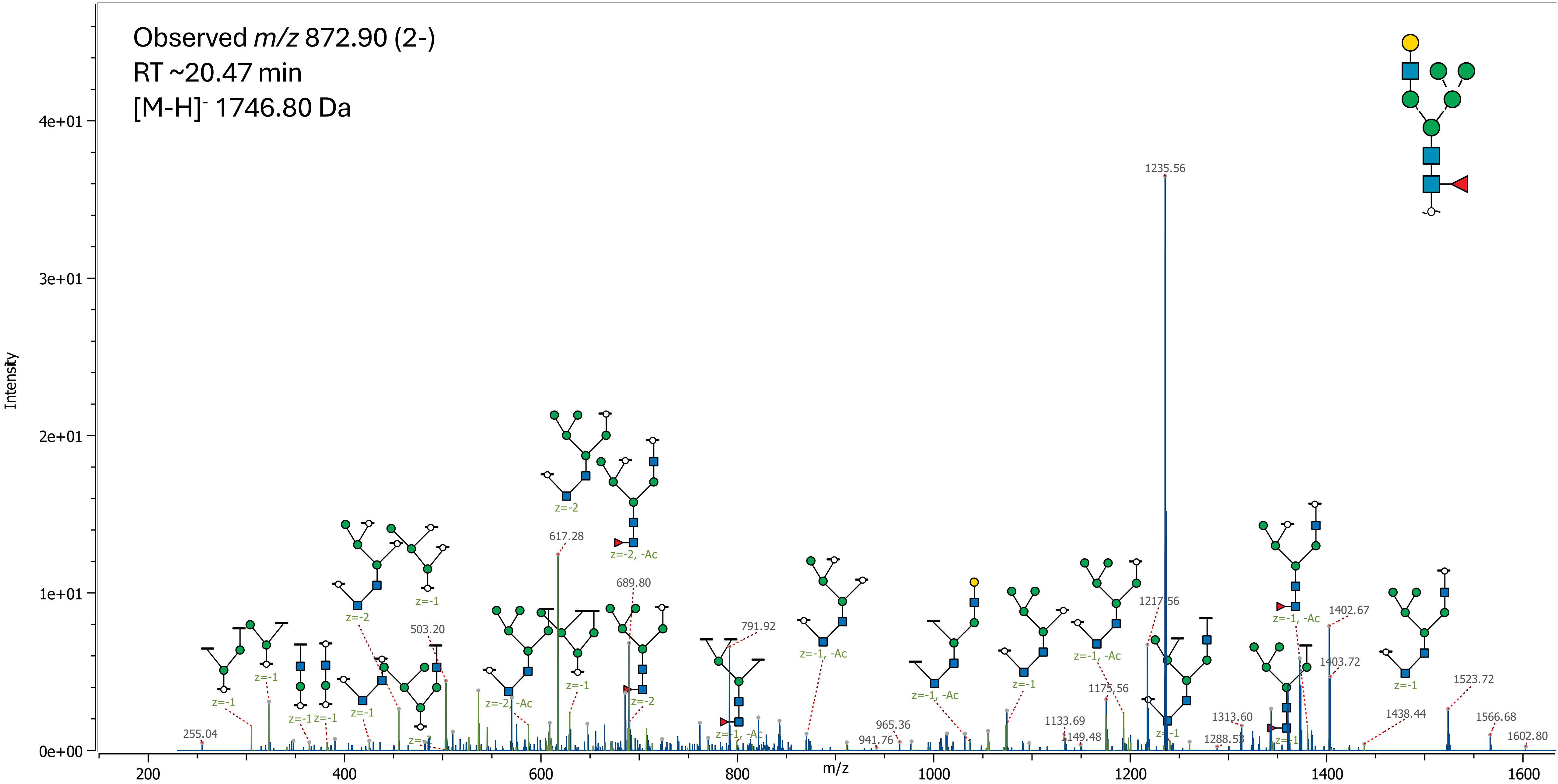

### 38 (Hex)1 (HexNAc)2 (NeuAc)1 + (Man)3(GlcNAc)2

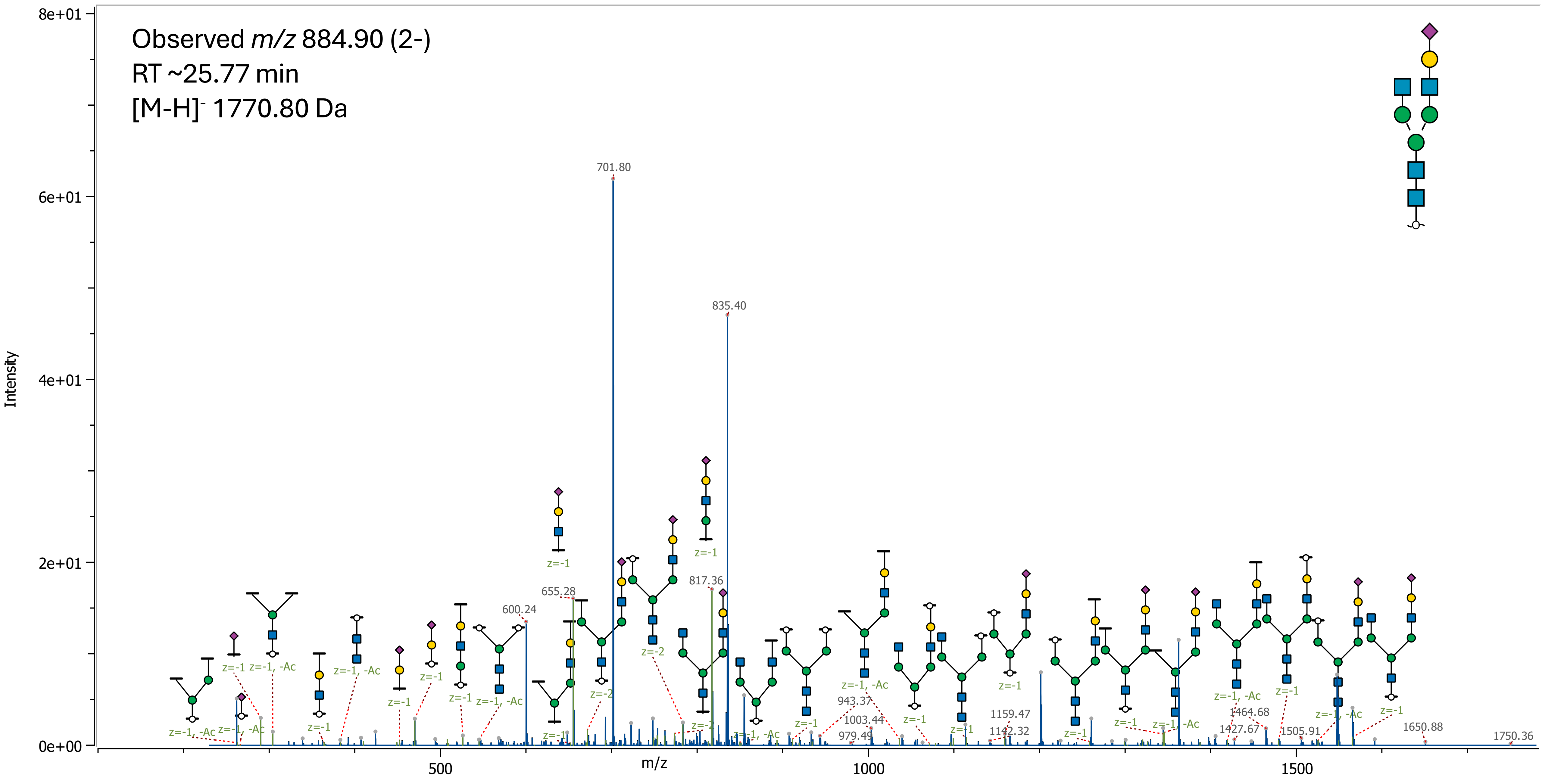

39 (Hex)2 (HexNAc)2 (Deoxyhexose)1 + (Man)3(GlcNAc)2

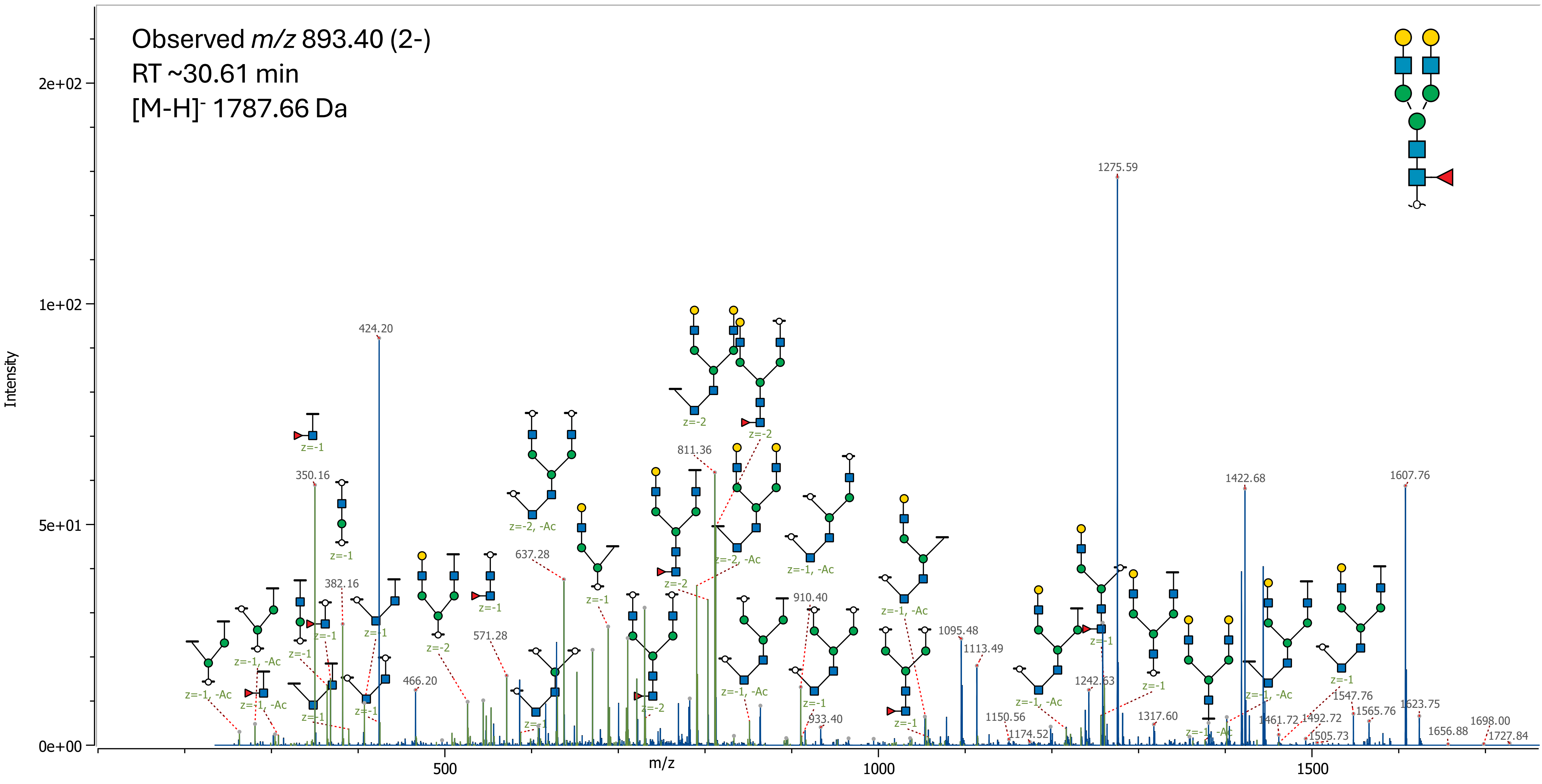

### 40 (HexNAc)3 (Deoxyhexose)2 + (Man)3(GlcNAc)2

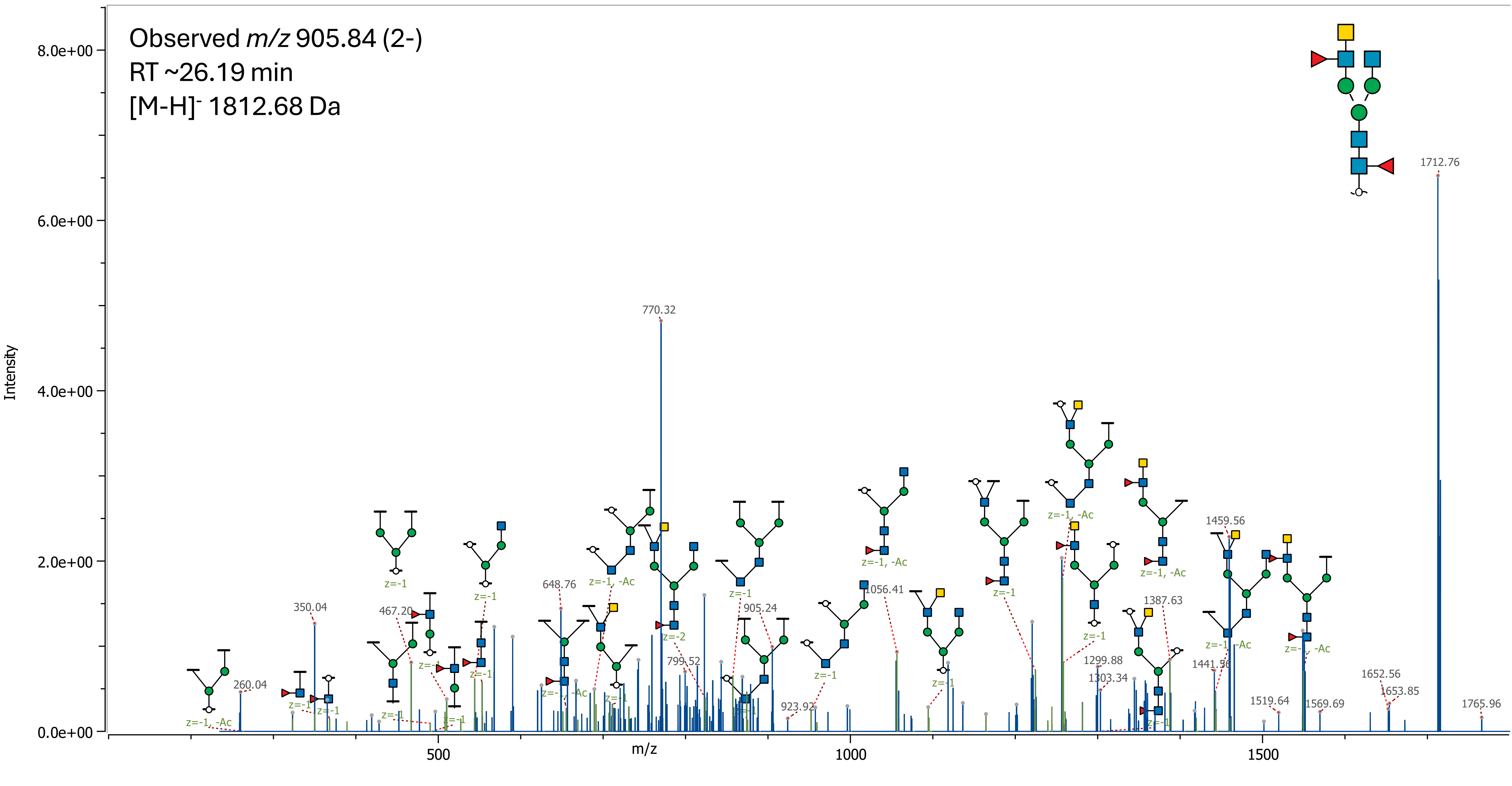

41(Hex)1 (HexNAc)3 (Deoxyhexose)1 + (Man)3(GlcNAc)2

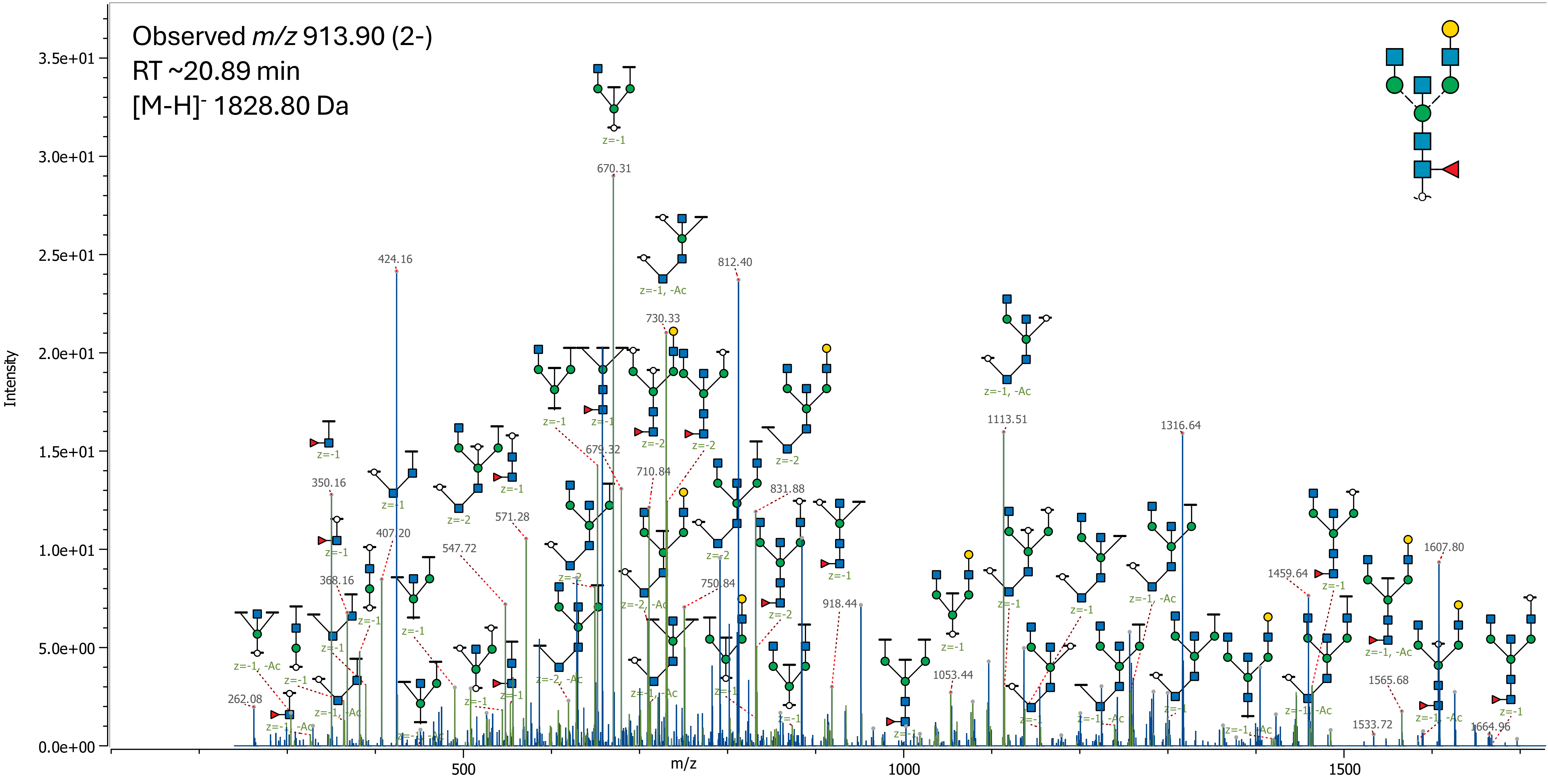

### 42 (Hex)2 (HexNAc)3 + (Man)3(GlcNAc)2

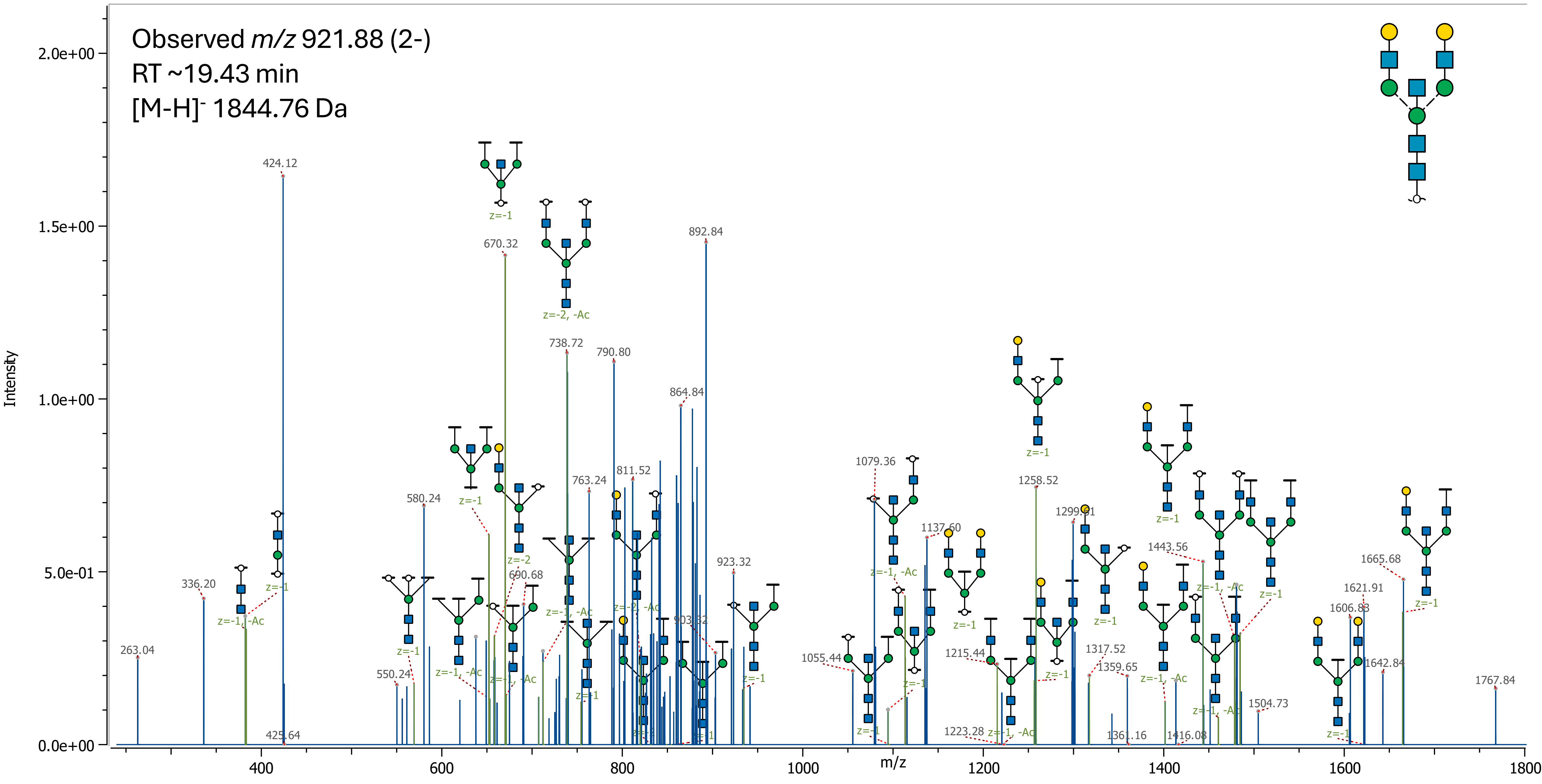

43 (HexNAc)4 (Deoxyhexose)1 + (Man)3(GlcNAc)2

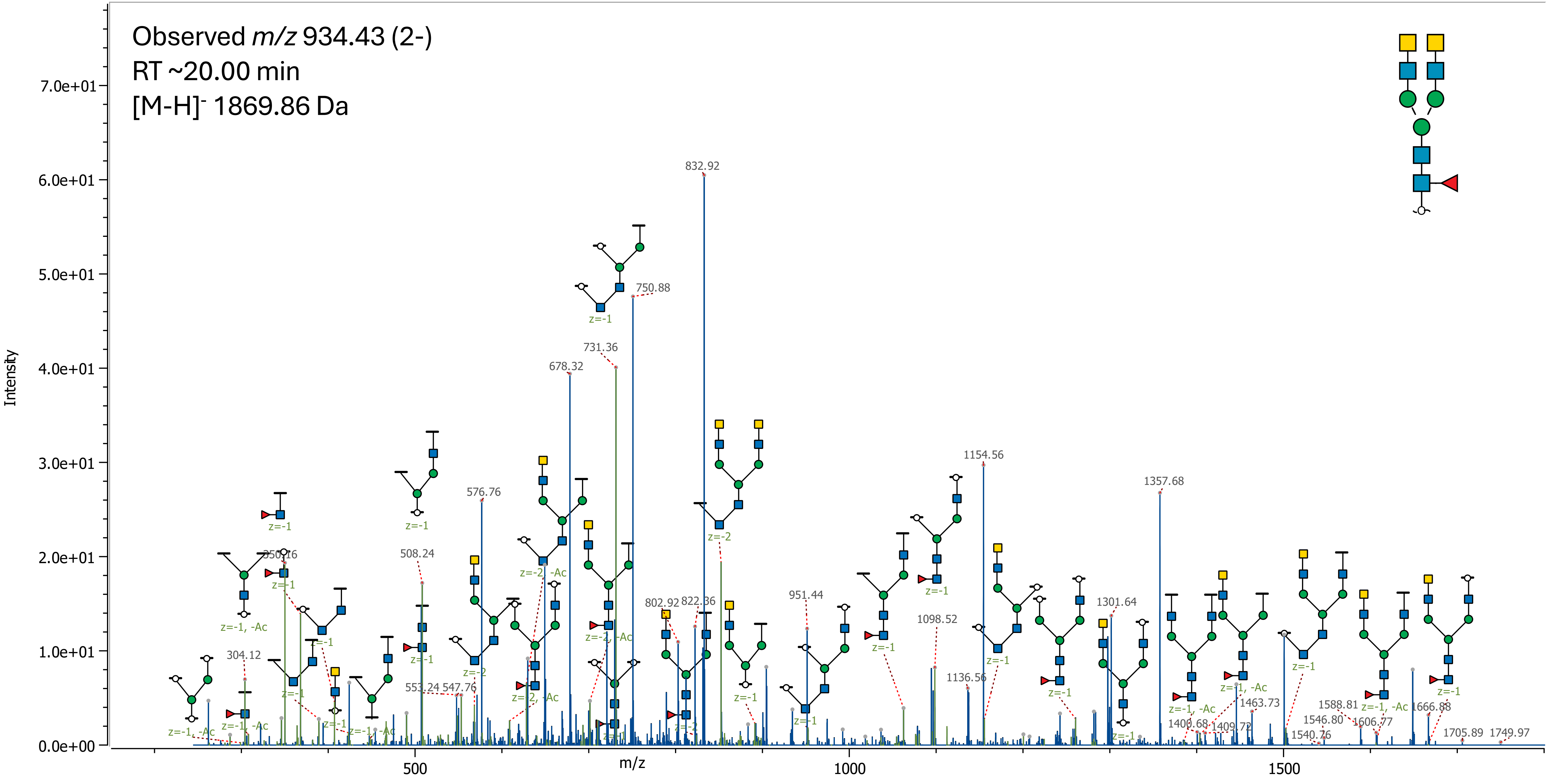

44 (Hex)2 (HexNAc)1 (Deoxyhexose)1 (NeuAc)1 + (Man)3(GlcNAc)2

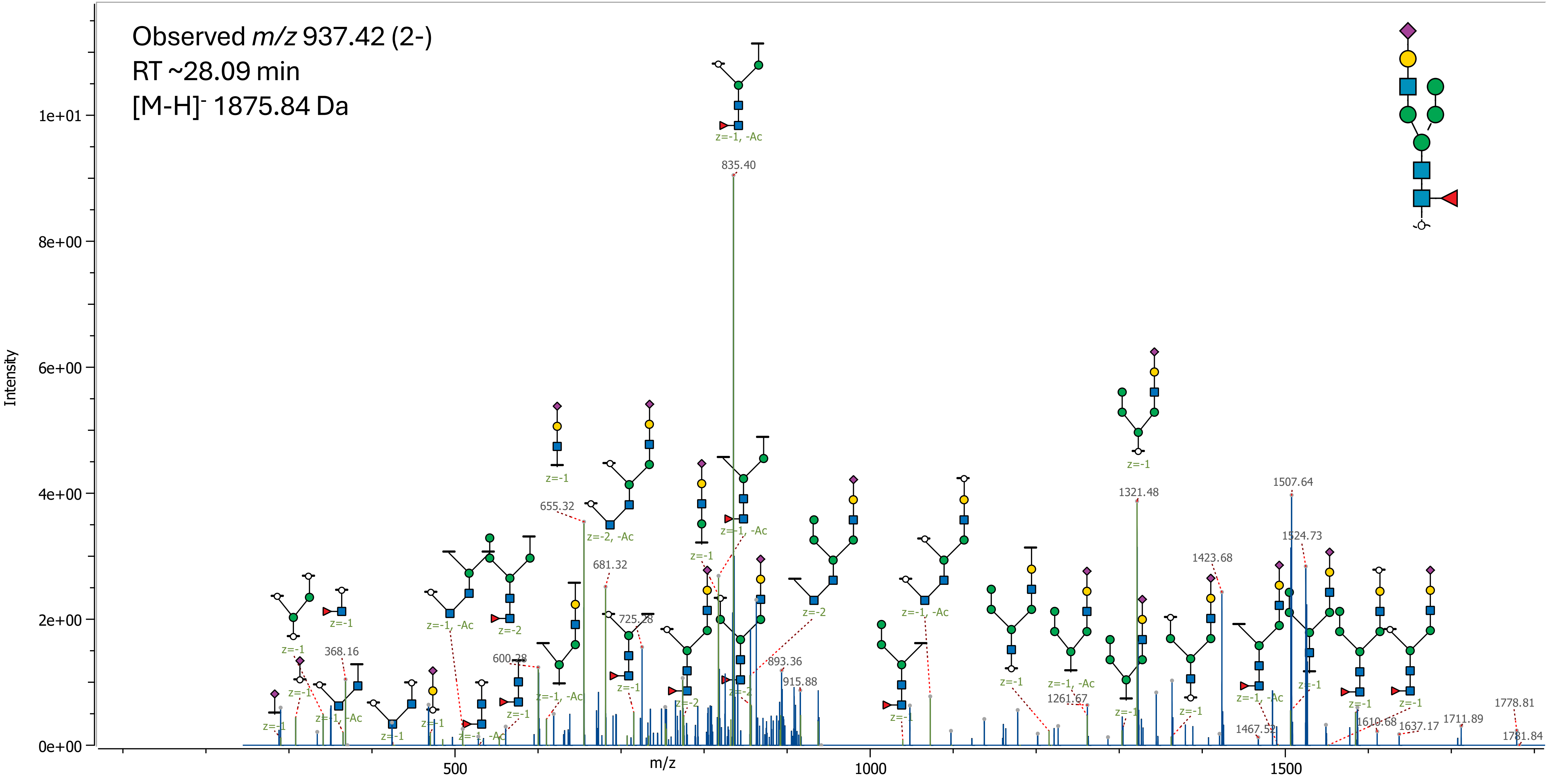

### 45 (Hex)3 (HexNAc)1 (NeuAc)1 + (Man)3(GlcNAc)2

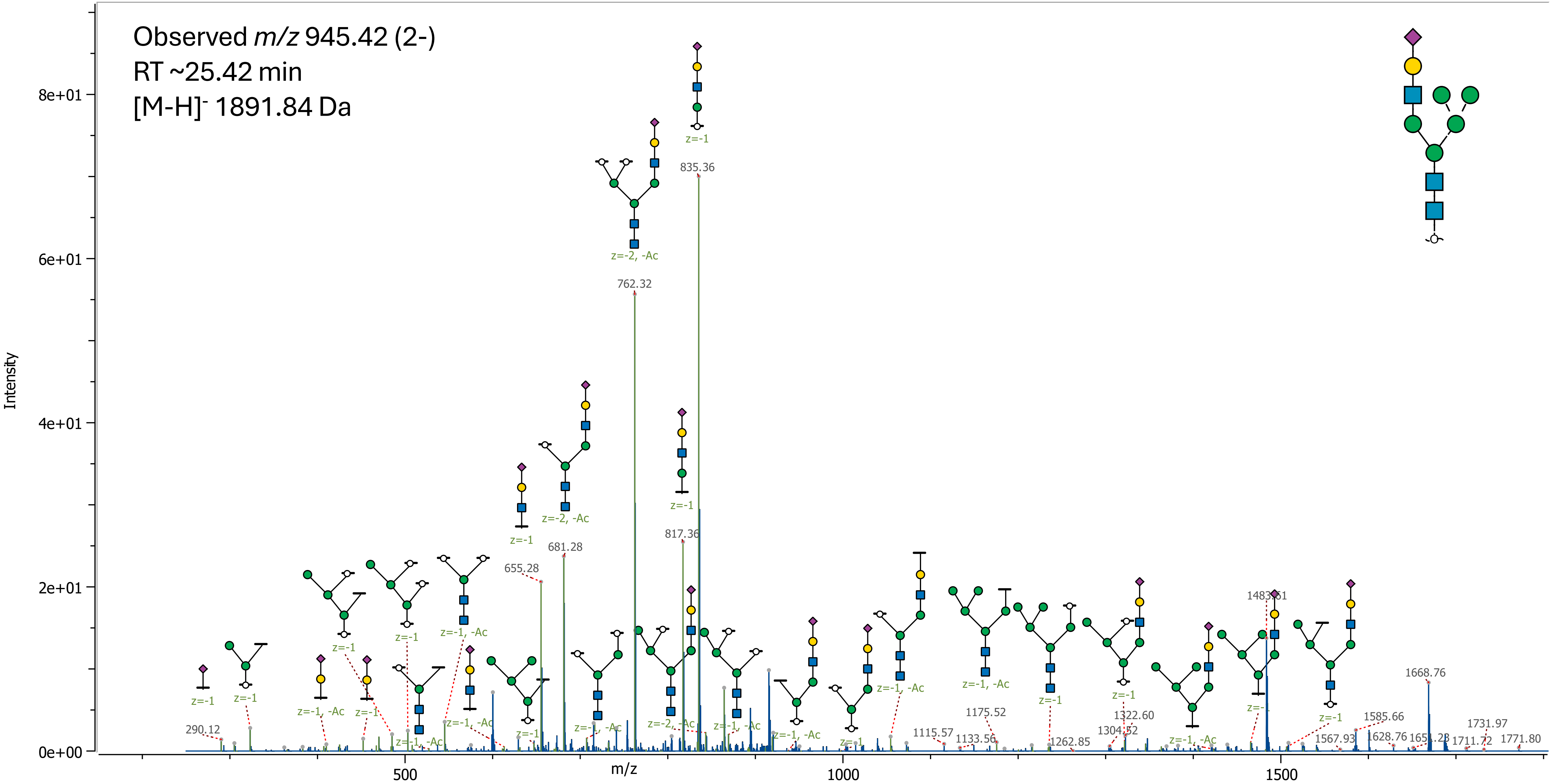

### 46 (Hex)1 (HexNAc)2 (Deoxyhexose)1 (NeuAc)1 + (Man)3(GlcNAc)2

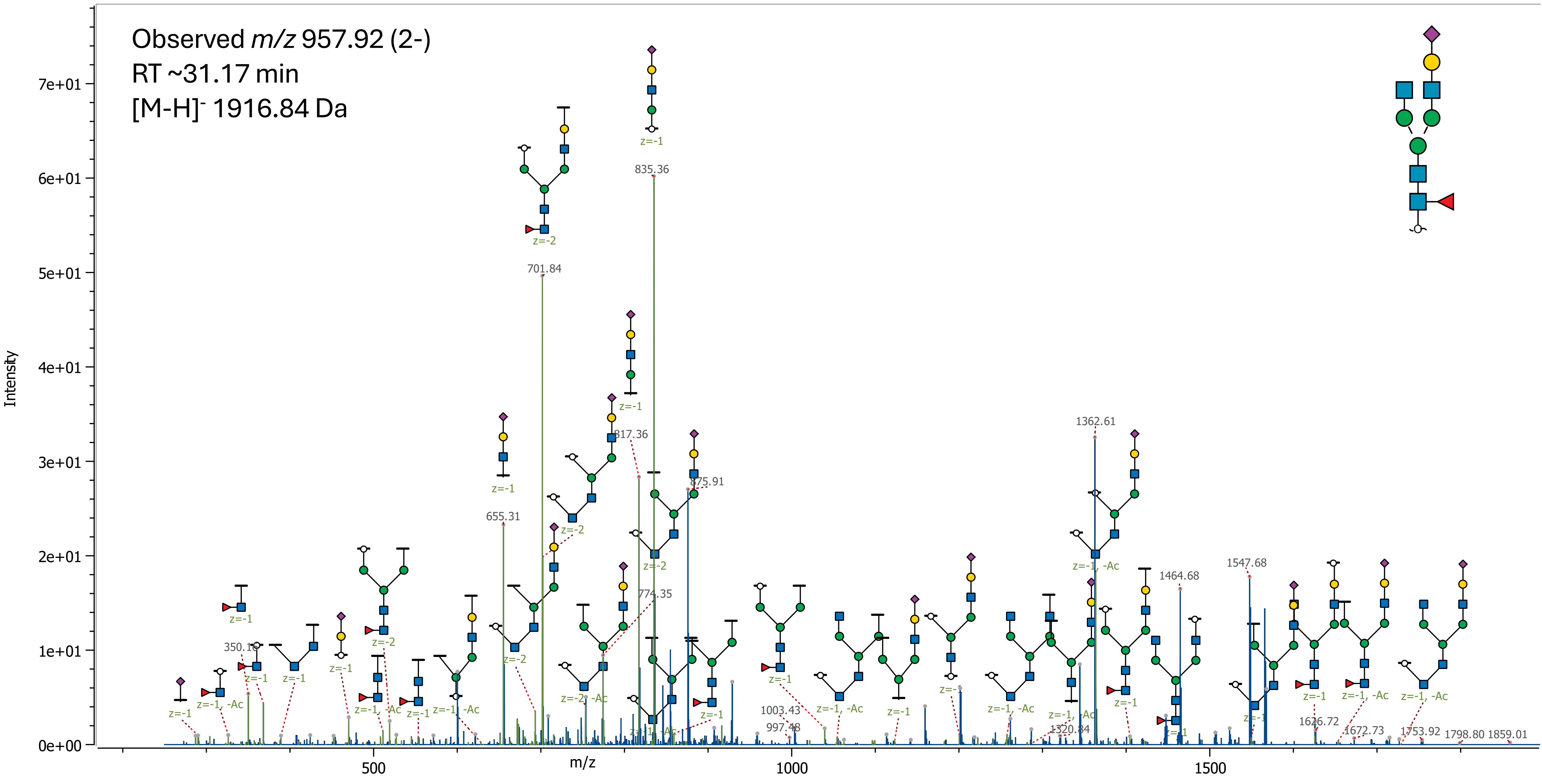

47 (Hex)1 (HexNAc)2 (Deoxyhexose)1 (NeuAc)1 + (Man)3(GlcNAc)2

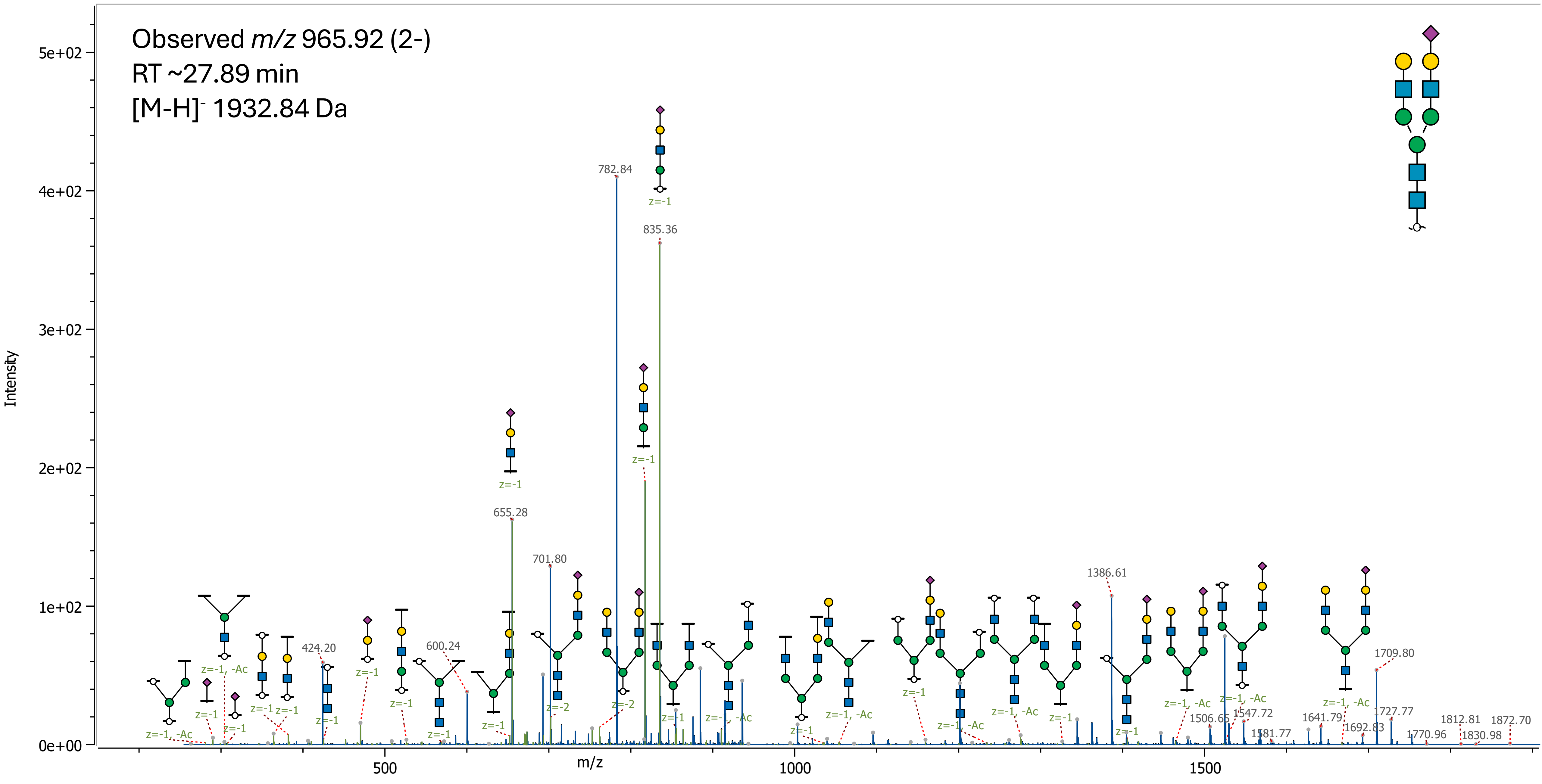

48 (Hex)3 (HexNAc)2 (Deoxyhexose)1 + (Man)3(GlcNAc)2

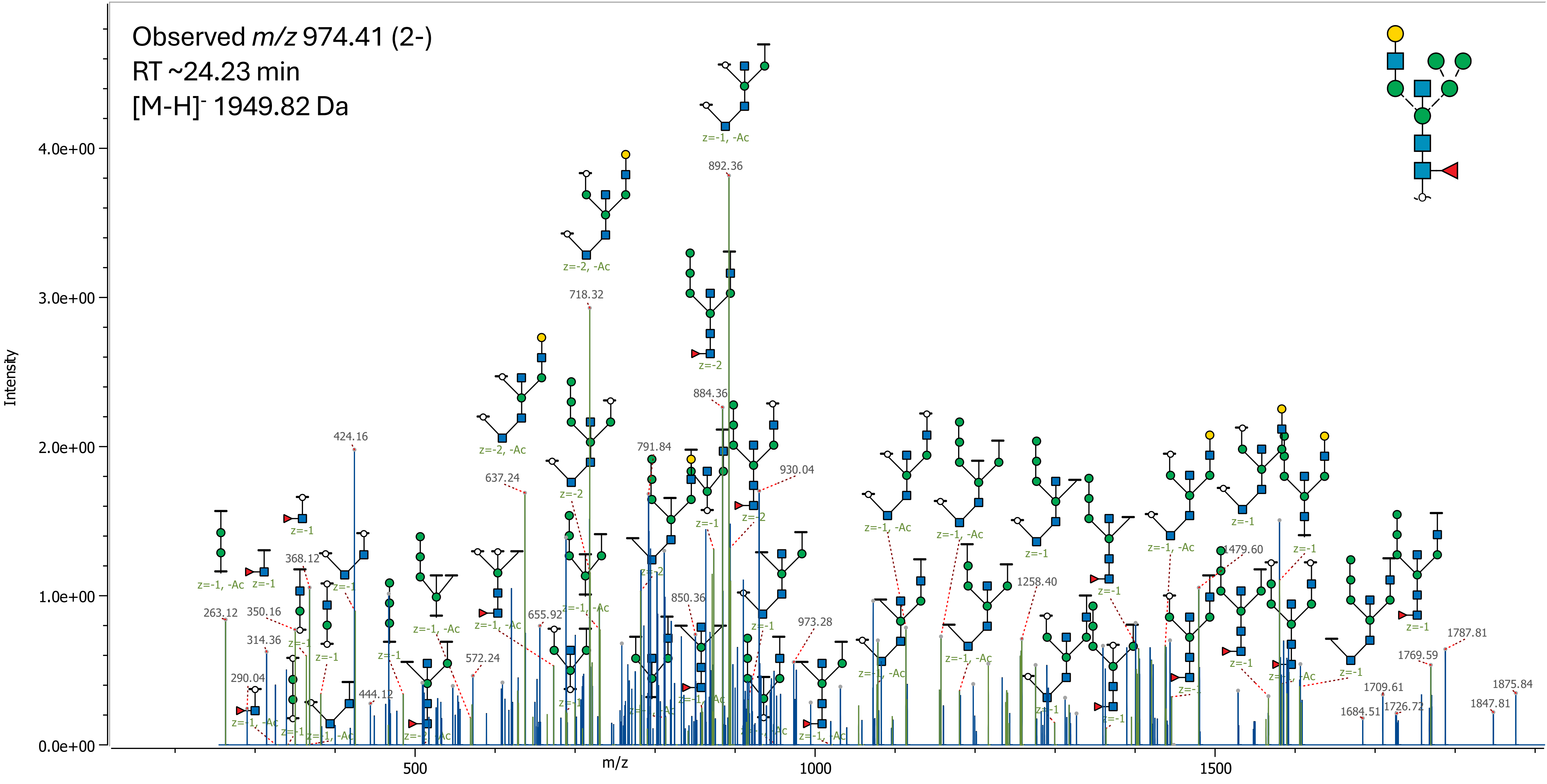

49 (Hex)1 (HexNAc)3 (NeuAc)1 + (Man)3(GlcNAc)2

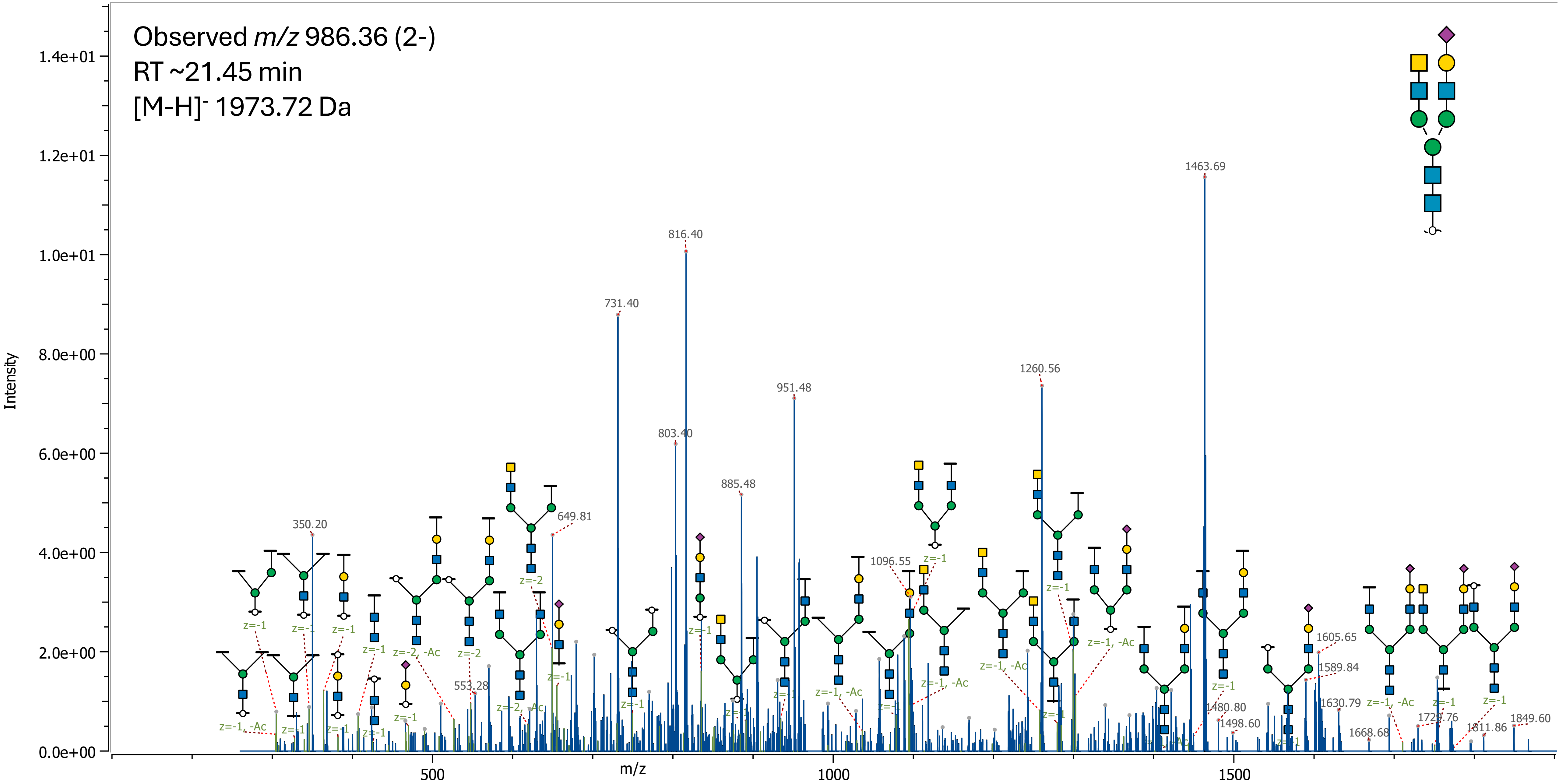

50 (Hex)2 (HexNAc)3 (Deoxyhexose)1 + (Man)3(GlcNAc)2

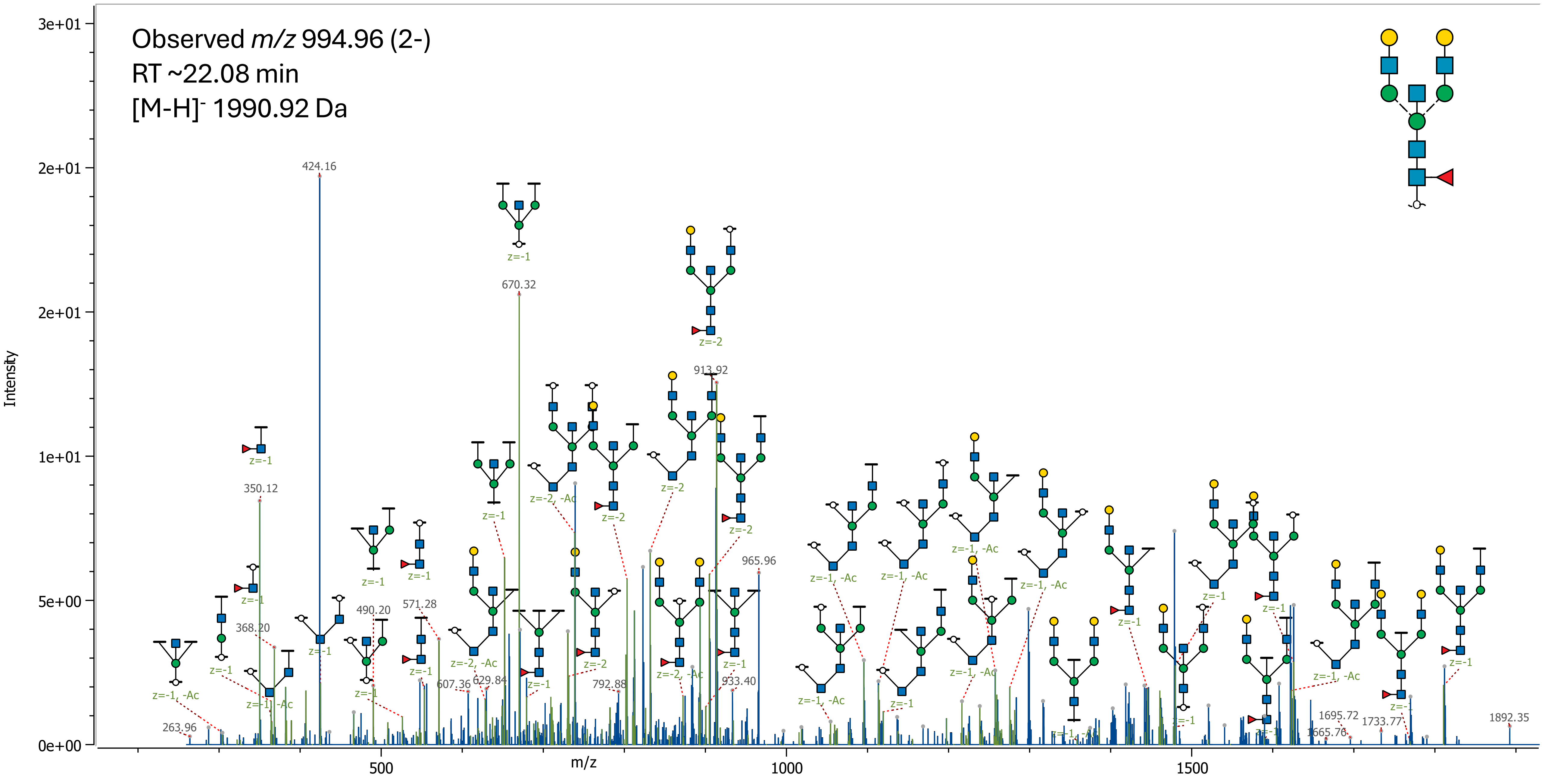

51 (Hex)3 (HexNAc)3 (Deoxyhexose)1 (NeuAc)3 + (Man)3(GlcNAc)2

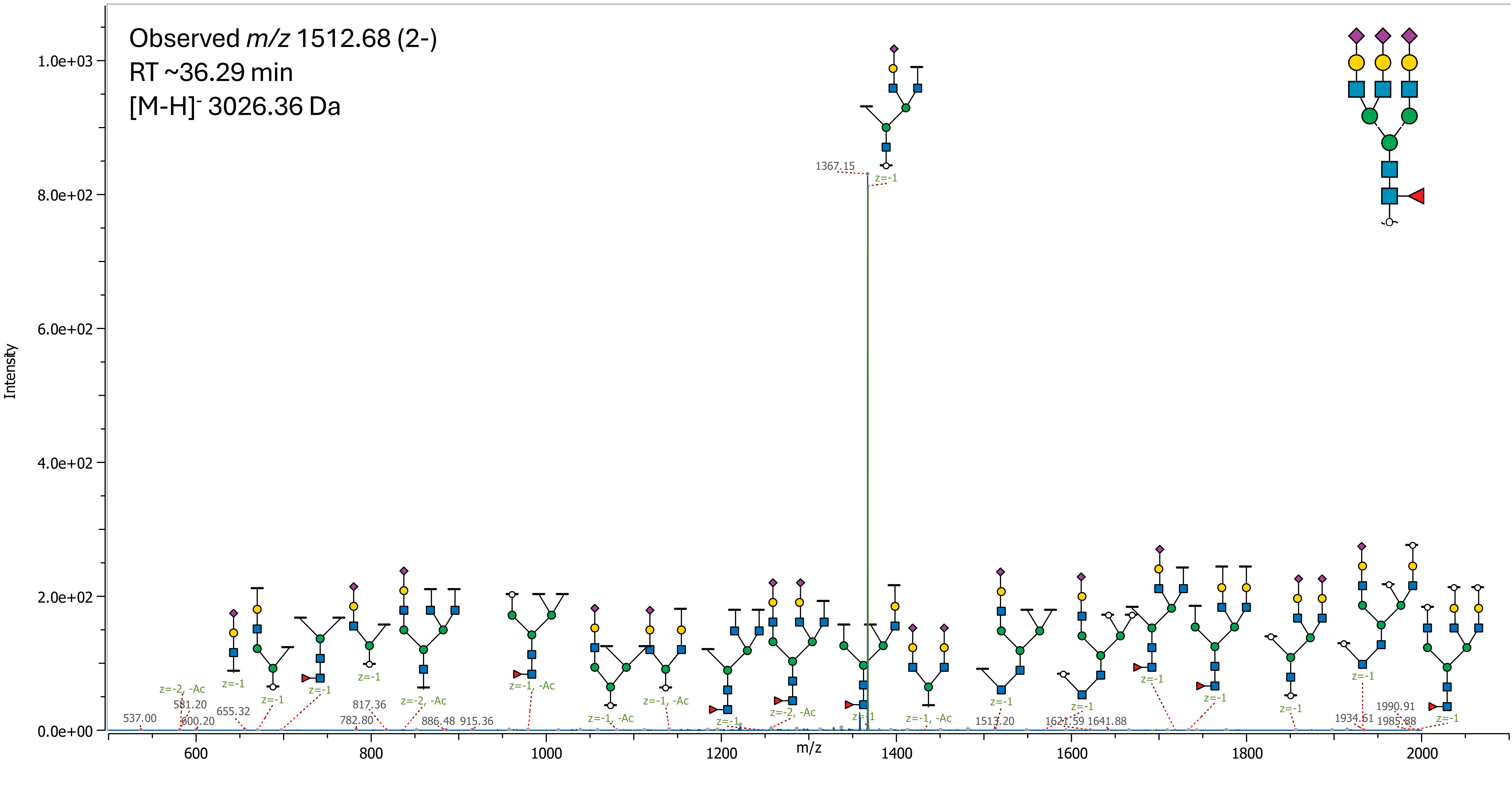

52 (Hex)2 (HexNAc)2 (Deoxyhexose)1 (NeuAc)1 + (Man)3(GlcNAc)2

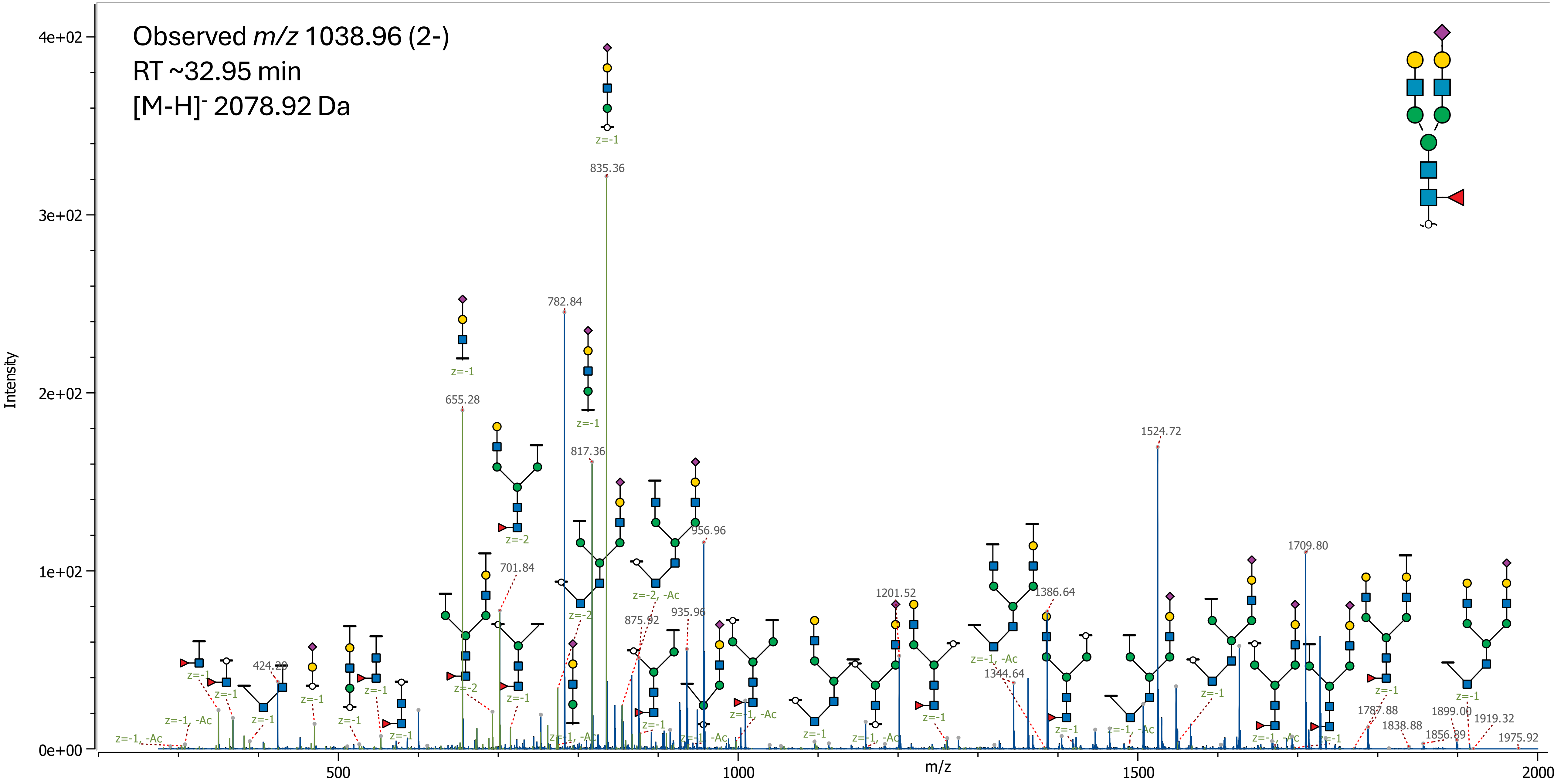

53 (Hex)1 (HexNAc)3 (Deoxyhexose)1 (NeuAc)1 + (Man)3(GlcNAc)2

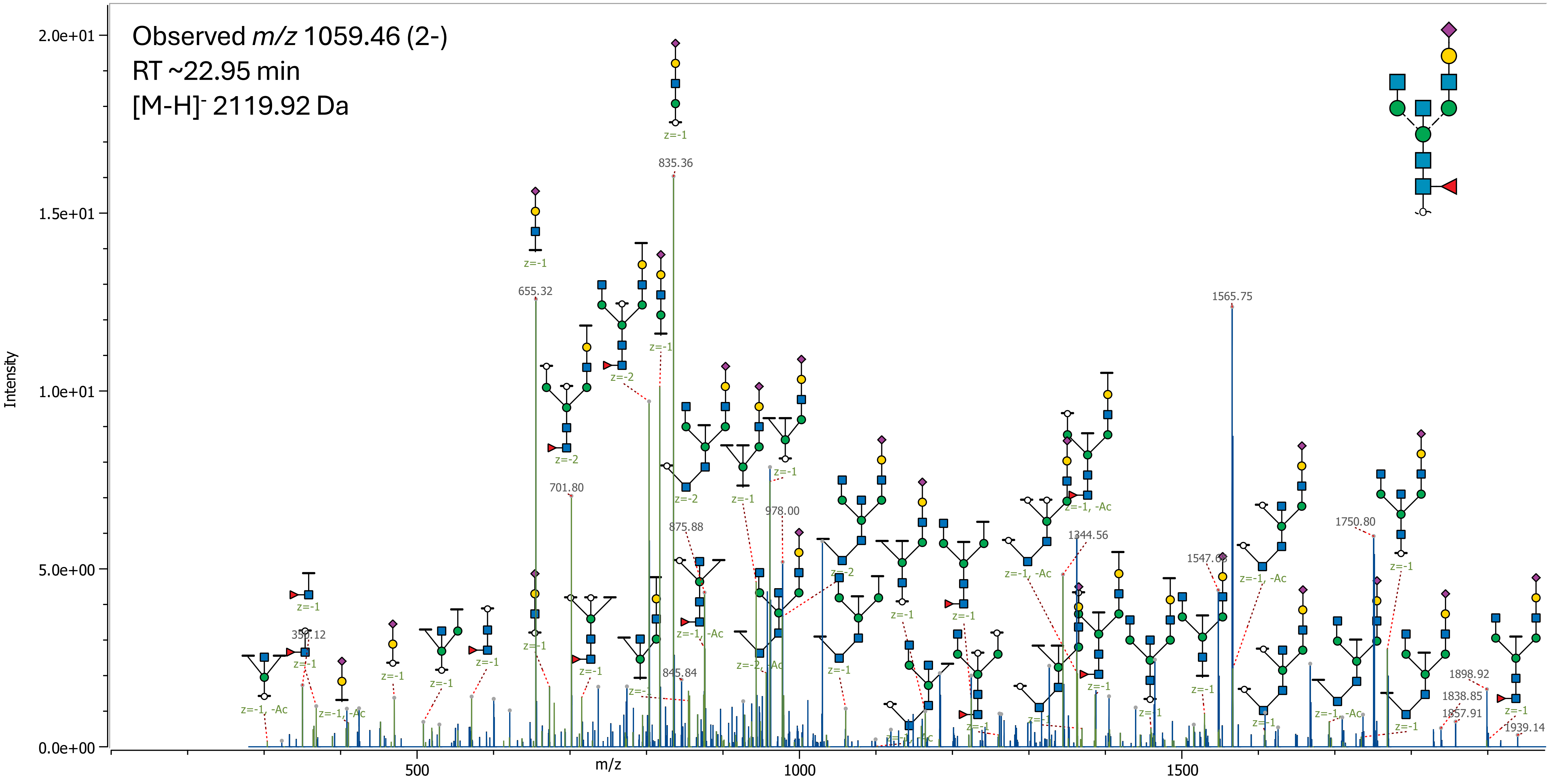

### 54 (Hex)2 (HexNAc)3 (NeuAc)1 + (Man)3(GlcNAc)2

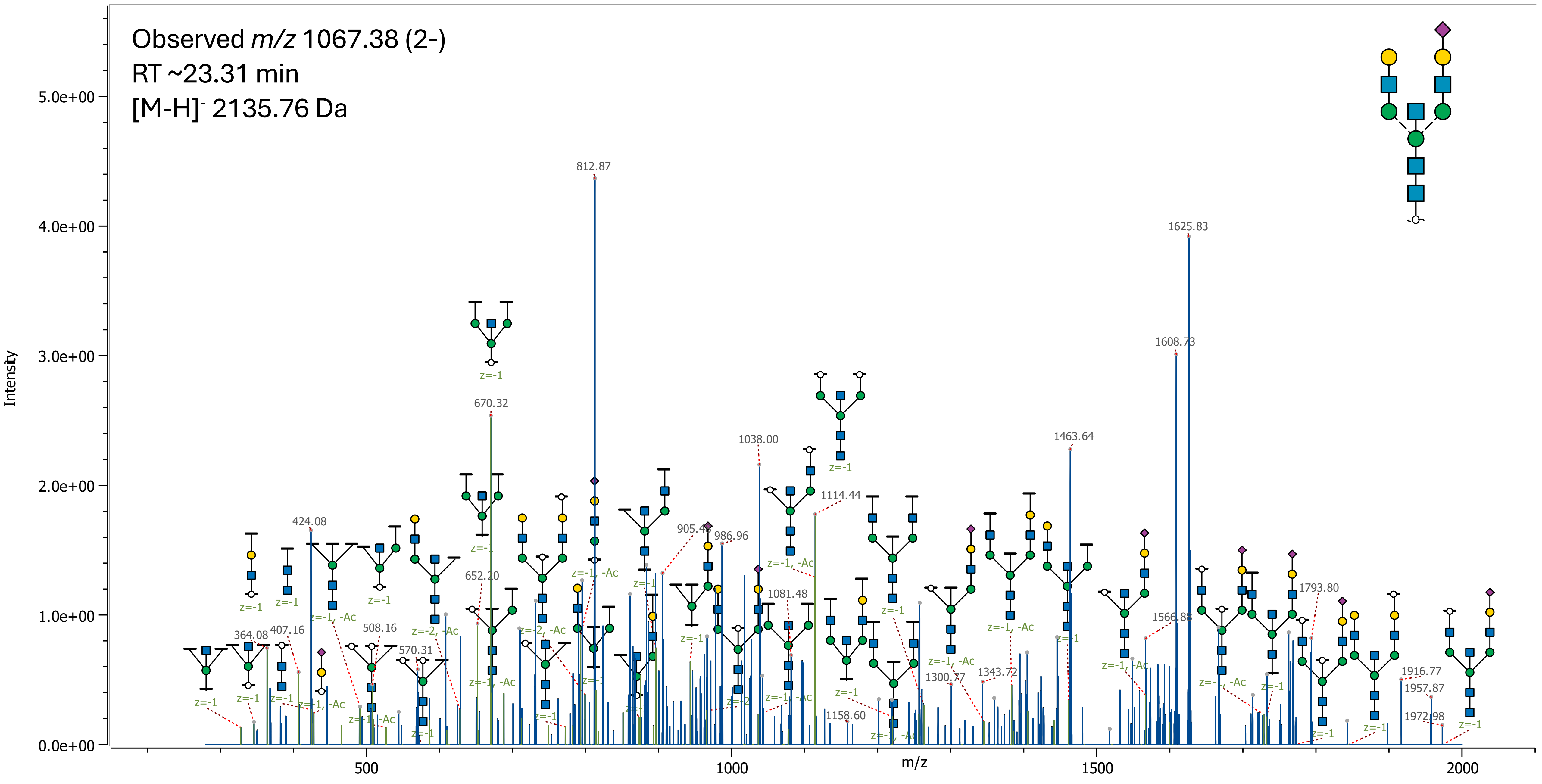

55 (Hex)3 (HexNAc)3 (Deoxyhexose)1 + (Man)3(GlcNAc)2

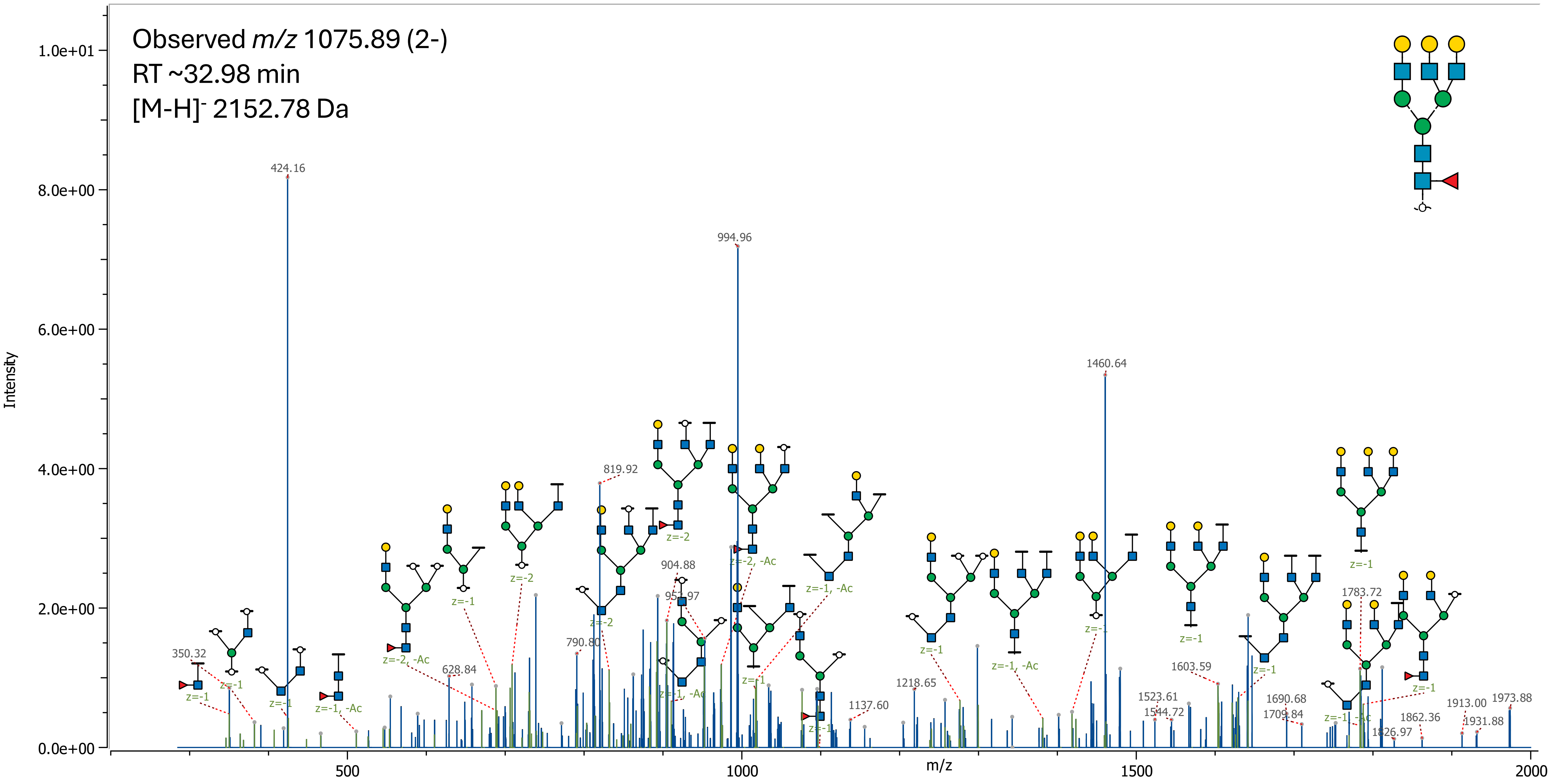

### 56 (Hex)2 (HexNAc)2 (NeuAc)2 + (Man)3(GlcNAc)2

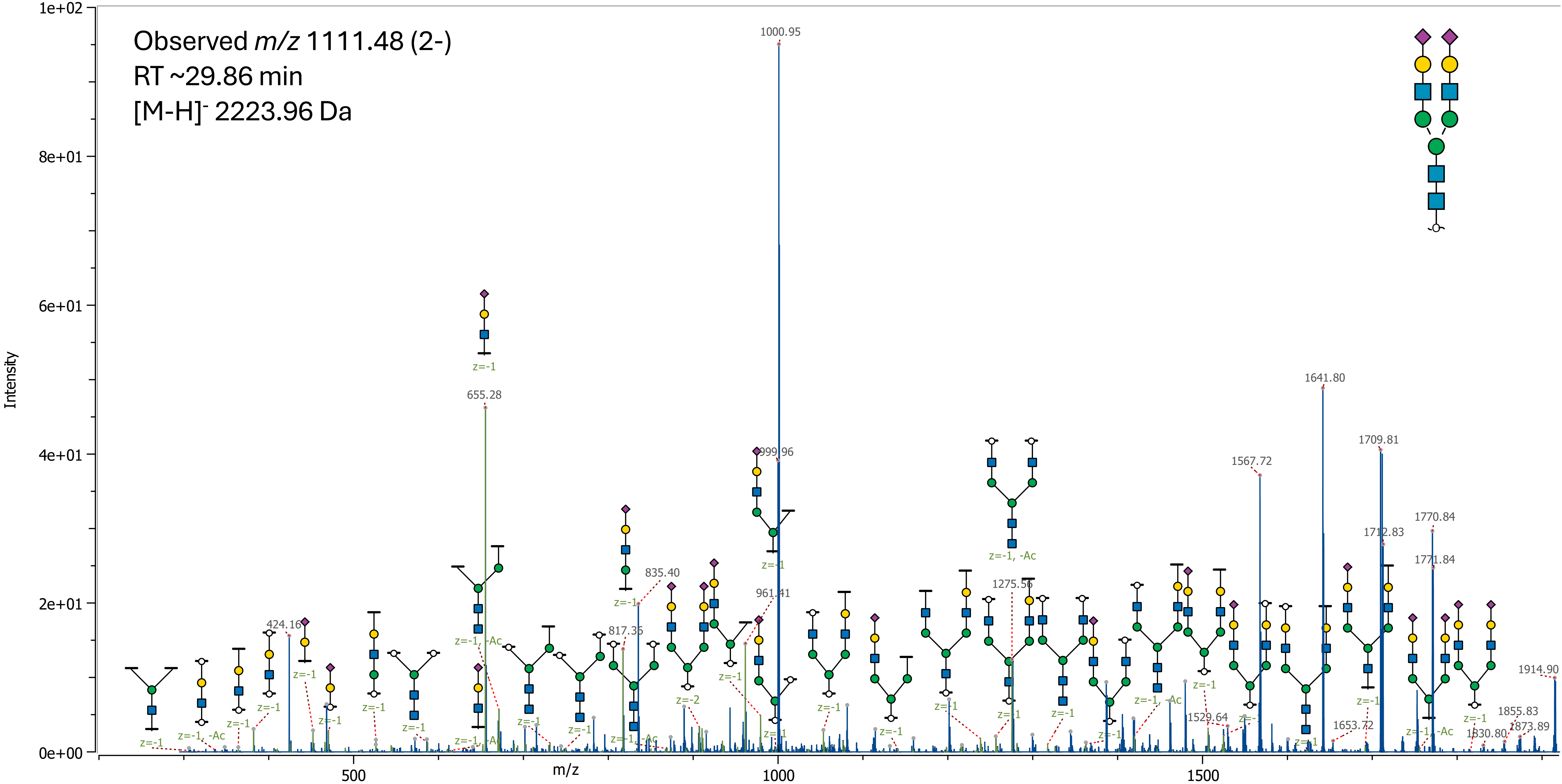

57 (Hex)2 (HexNAc)2 (Deoxyhexose)2 (NeuAc)1 + (Man)3(GlcNAc)2

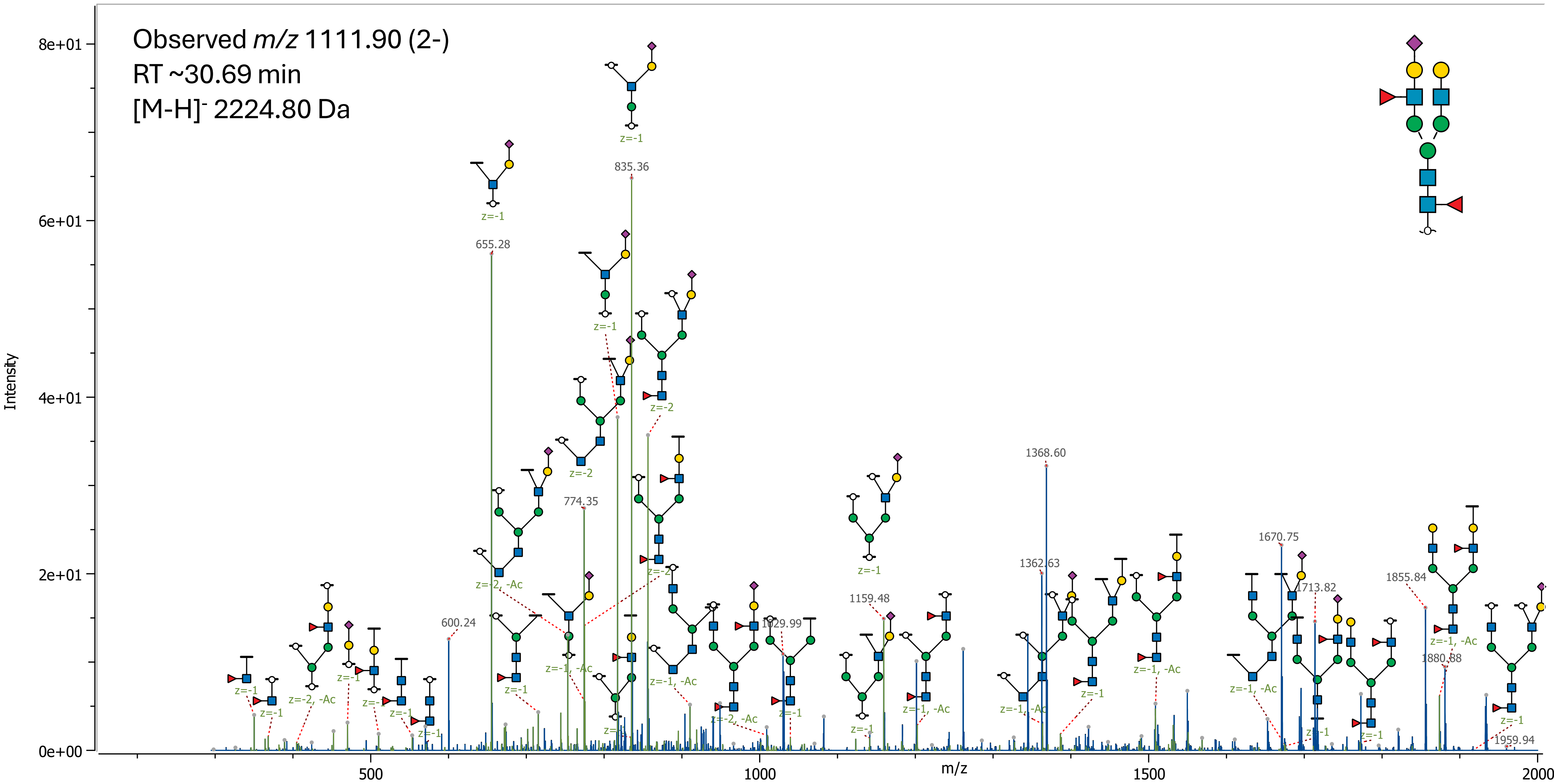

58 (Hex)1 (HexNAc)3 (Deoxyhexose)2 (NeuAc)1 + (Man)3(GlcNAc)2

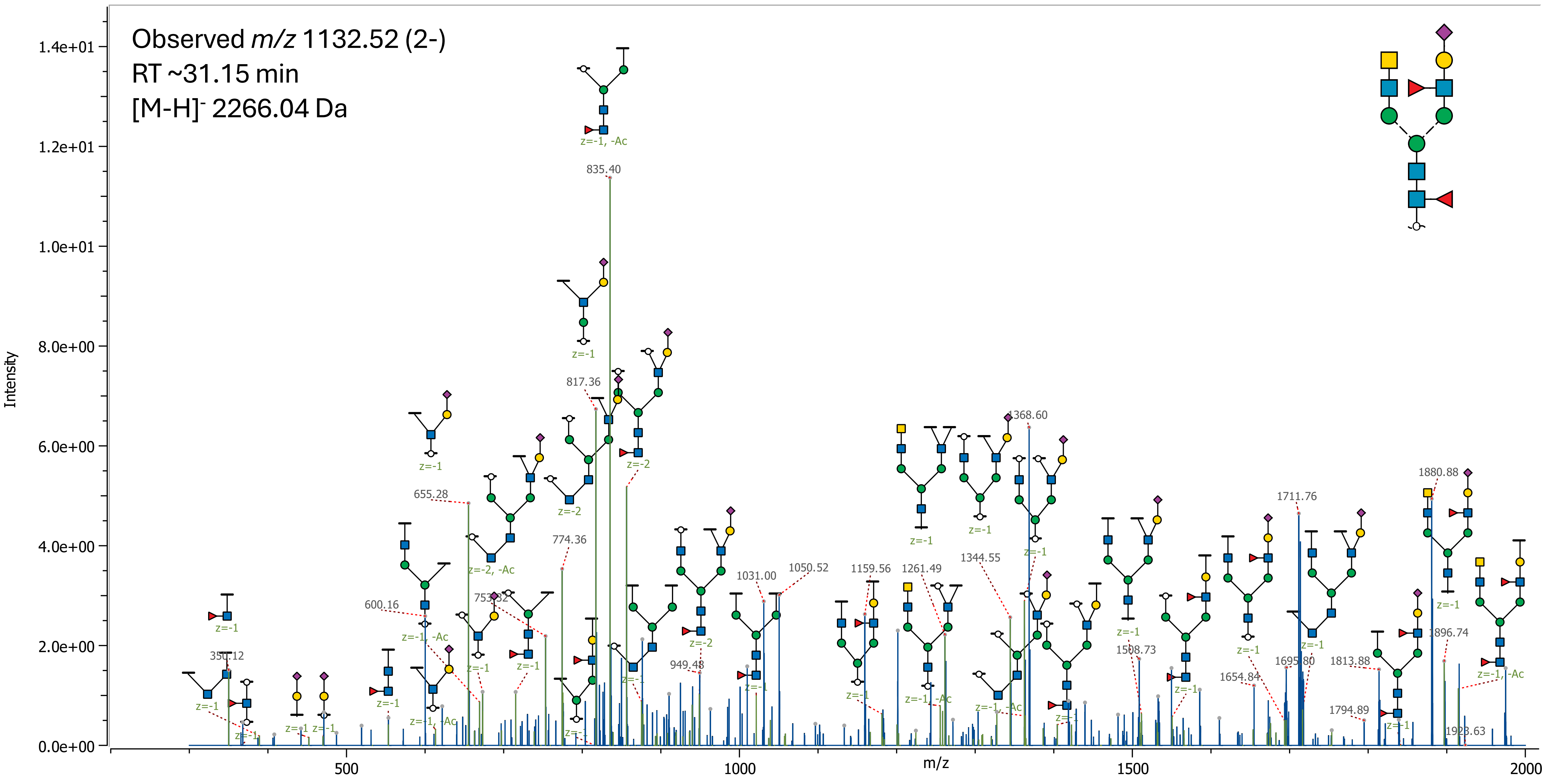

59 (Hex)2 (HexNAc)3 (Deoxyhexose)1 (NeuAc)1 + (Man)3(GlcNAc)2

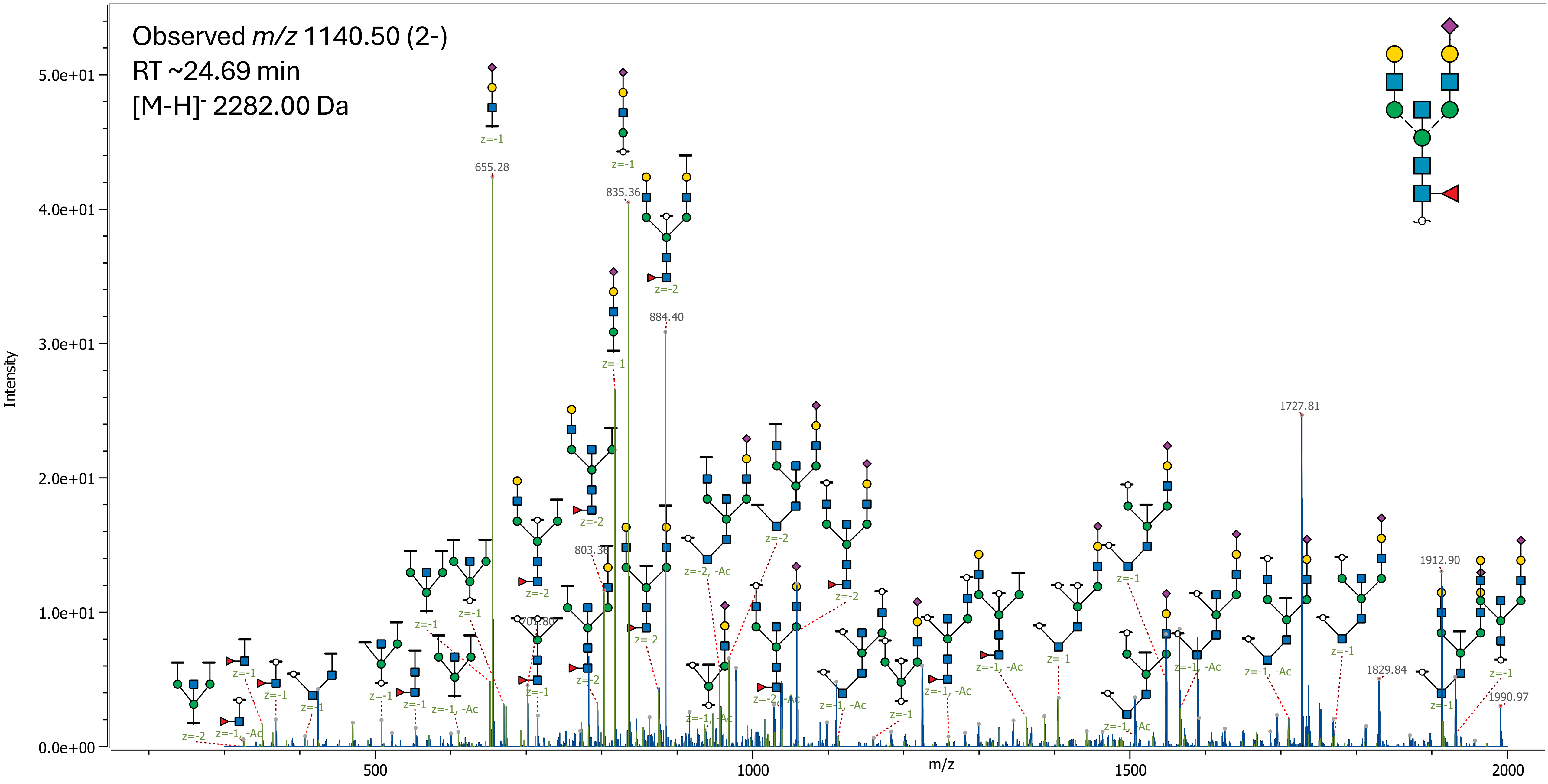

60 (Hex)3 (HexNAc)3 (NeuAc)1 + (Man)3(GlcNAc)2

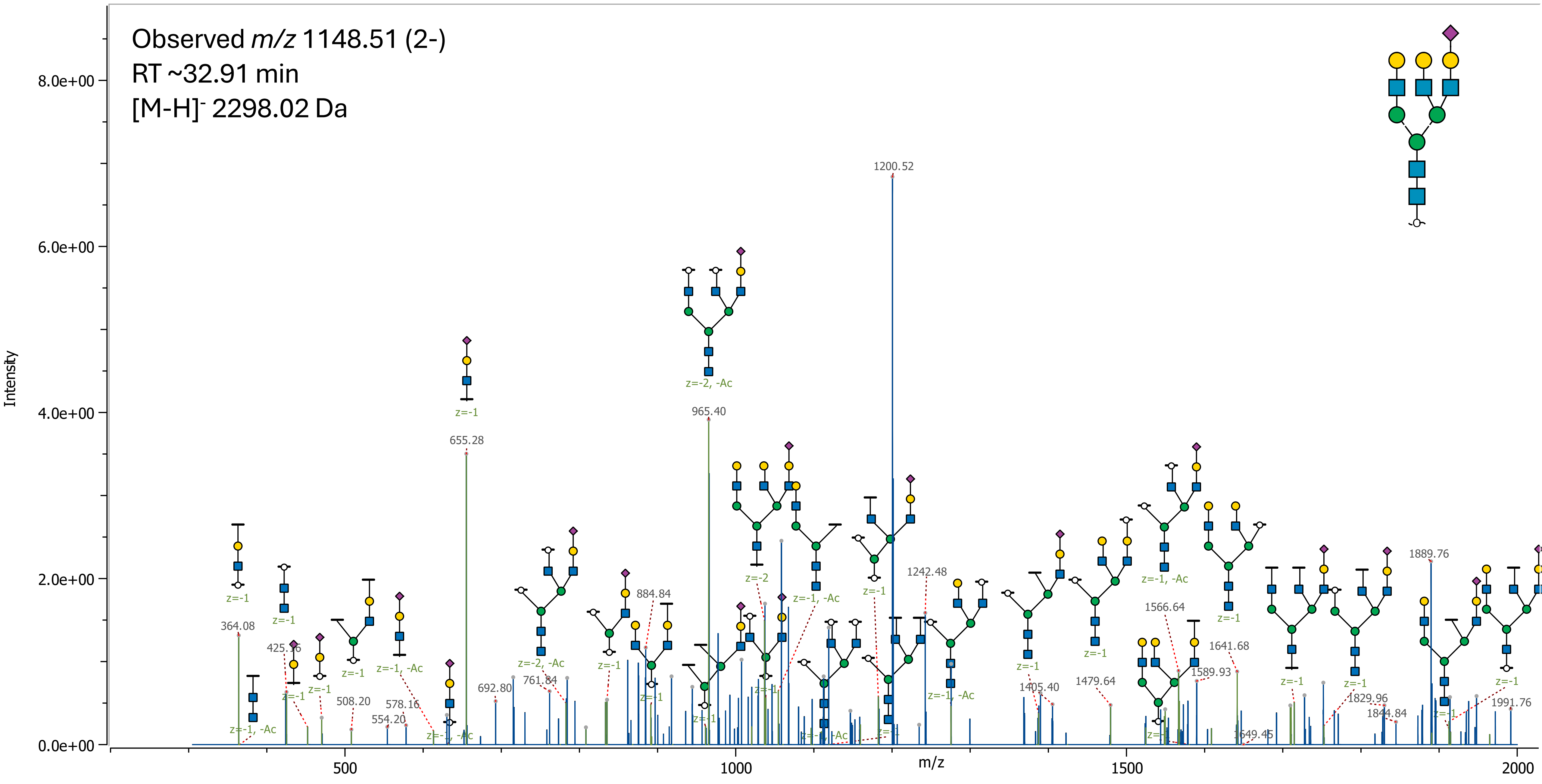

61 (Hex)1 (HexNAc)4 (Deoxyhexose)1 (NeuAc)1 + (Man)3(GlcNAc)2

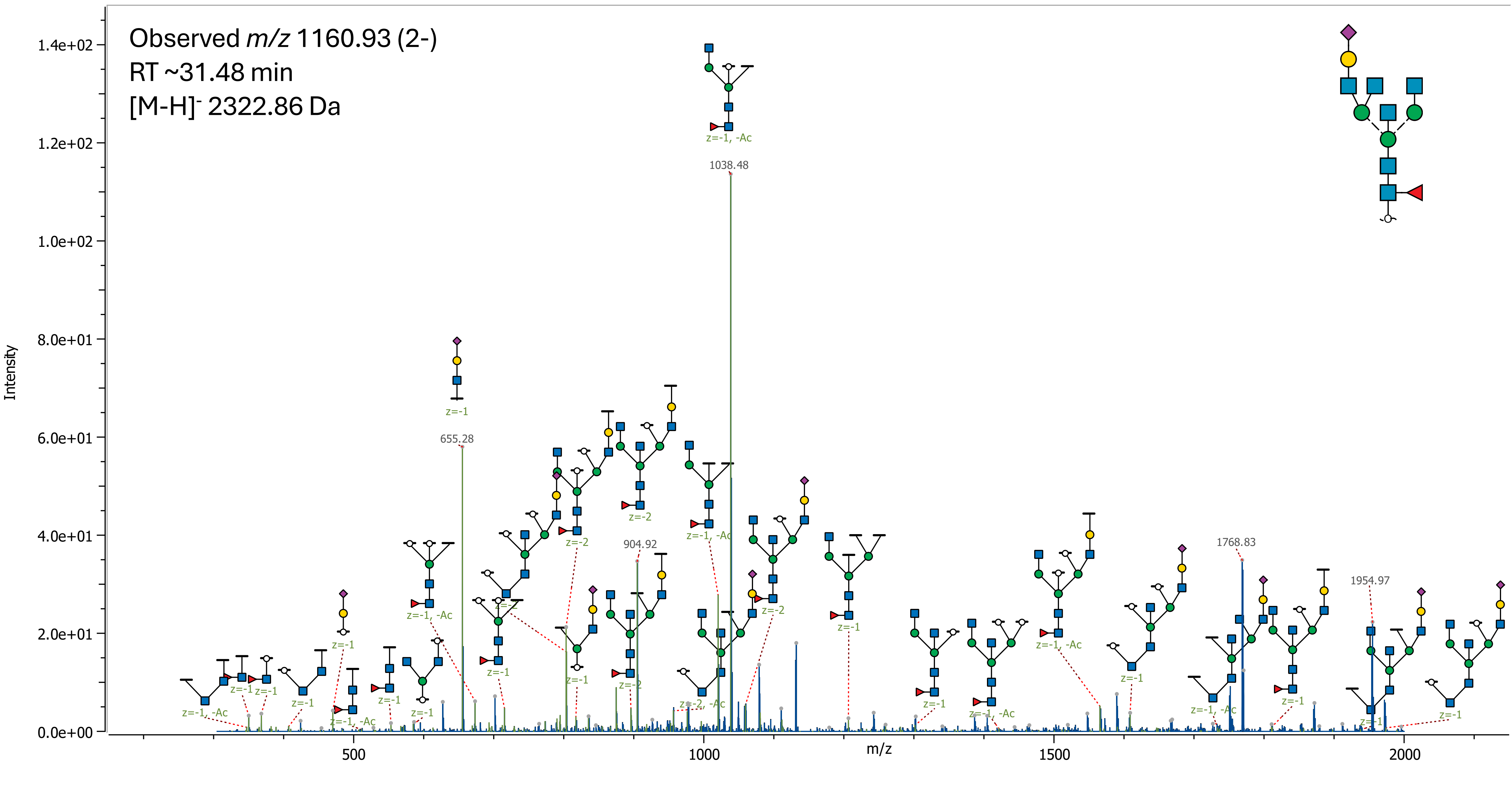

62 (Hex)2 (HexNAc)2 (Deoxyhexose)1 (NeuAc)2 + (Man)3(GlcNAc)2

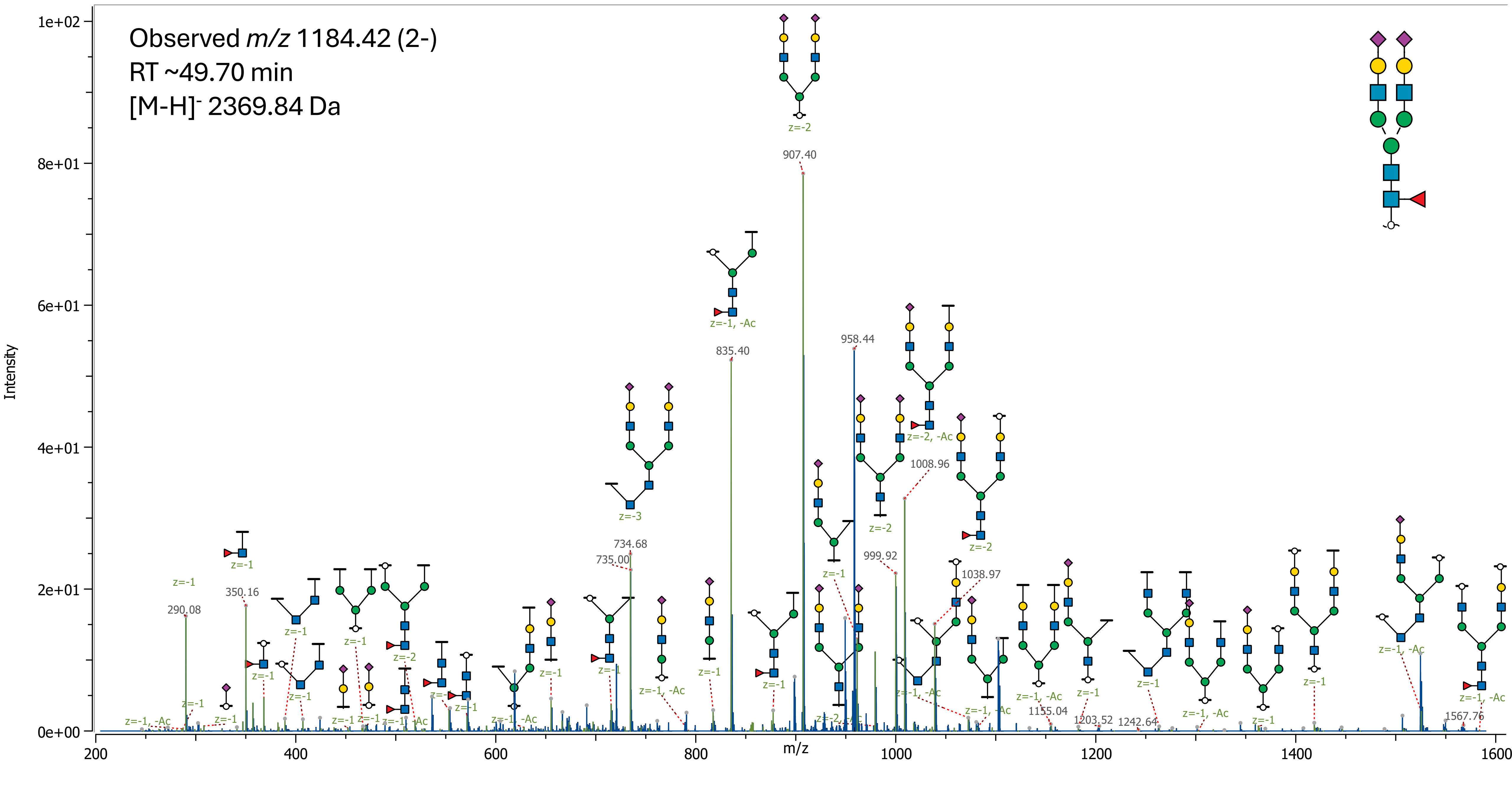

63 (Hex)1 (HexNAc)3 (Deoxyhexose)1 (NeuAc)2 + (Man)3(GlcNAc)2

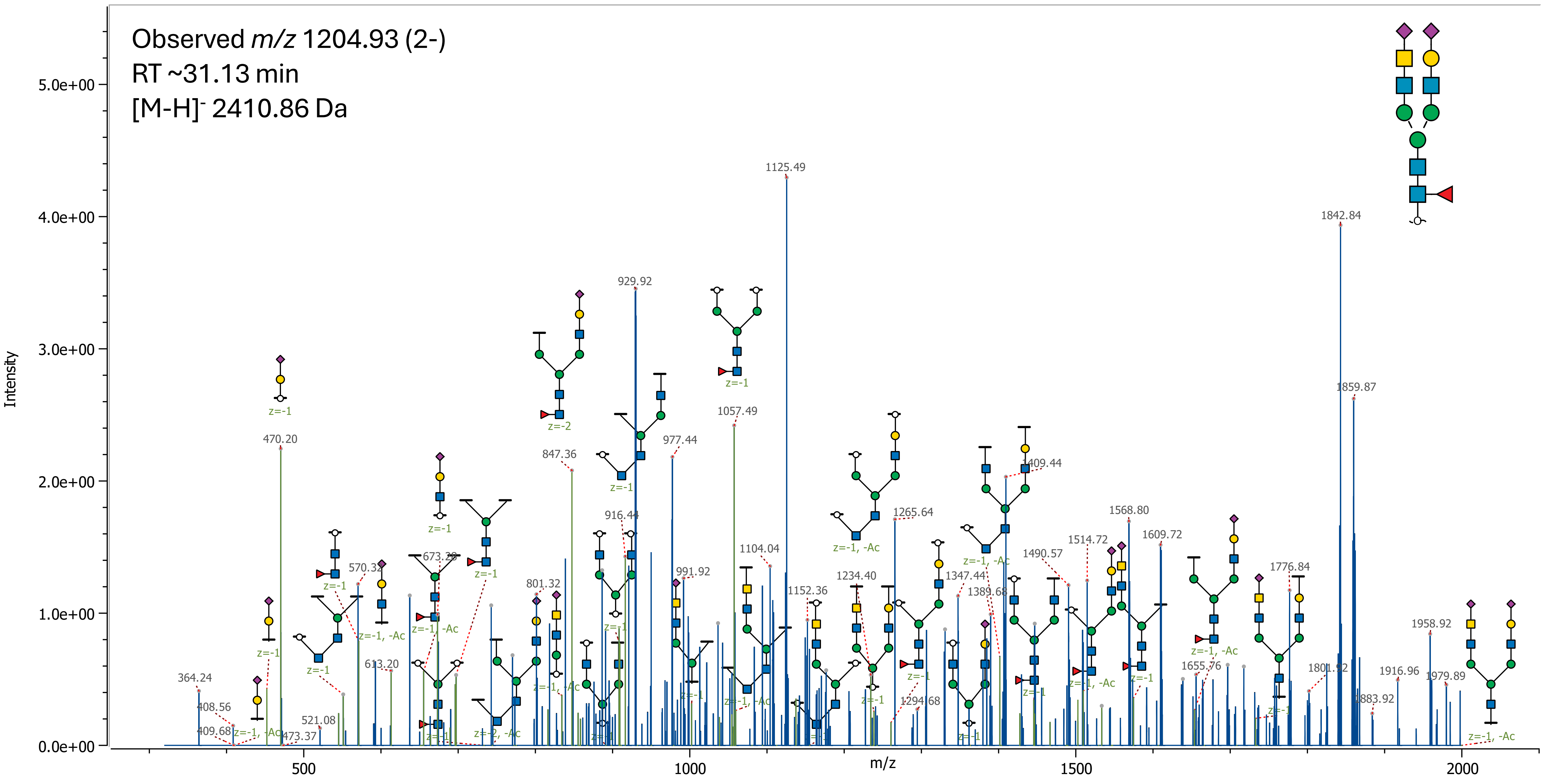

### 64 (Hex)2 (HexNAc)3 (NeuAc)2 + (Man)3(GlcNAc)2

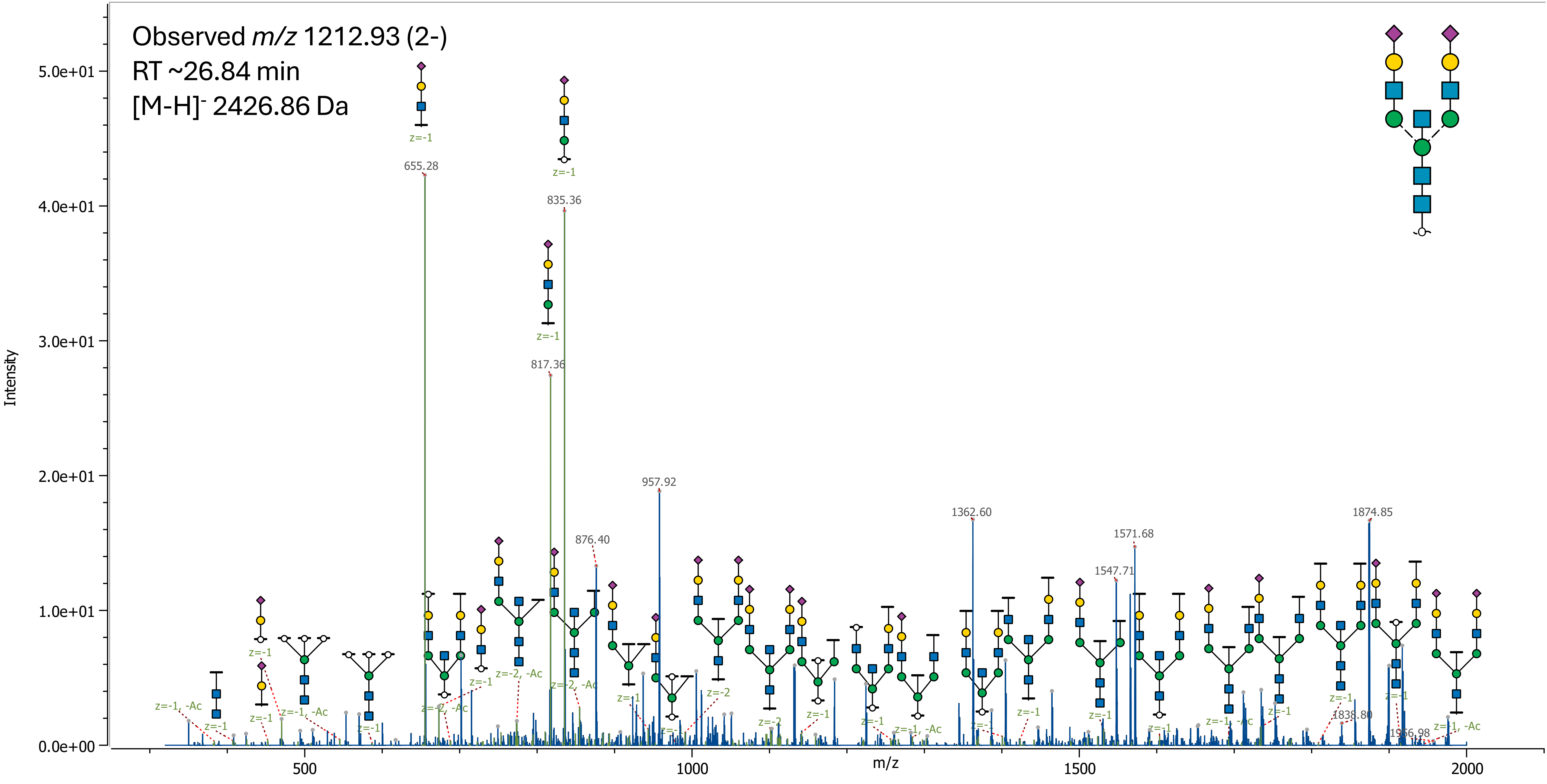

65 (Hex)2 (HexNAc)3 (Deoxyhexose)2 (NeuAc)1 + (Man)3(GlcNAc)2

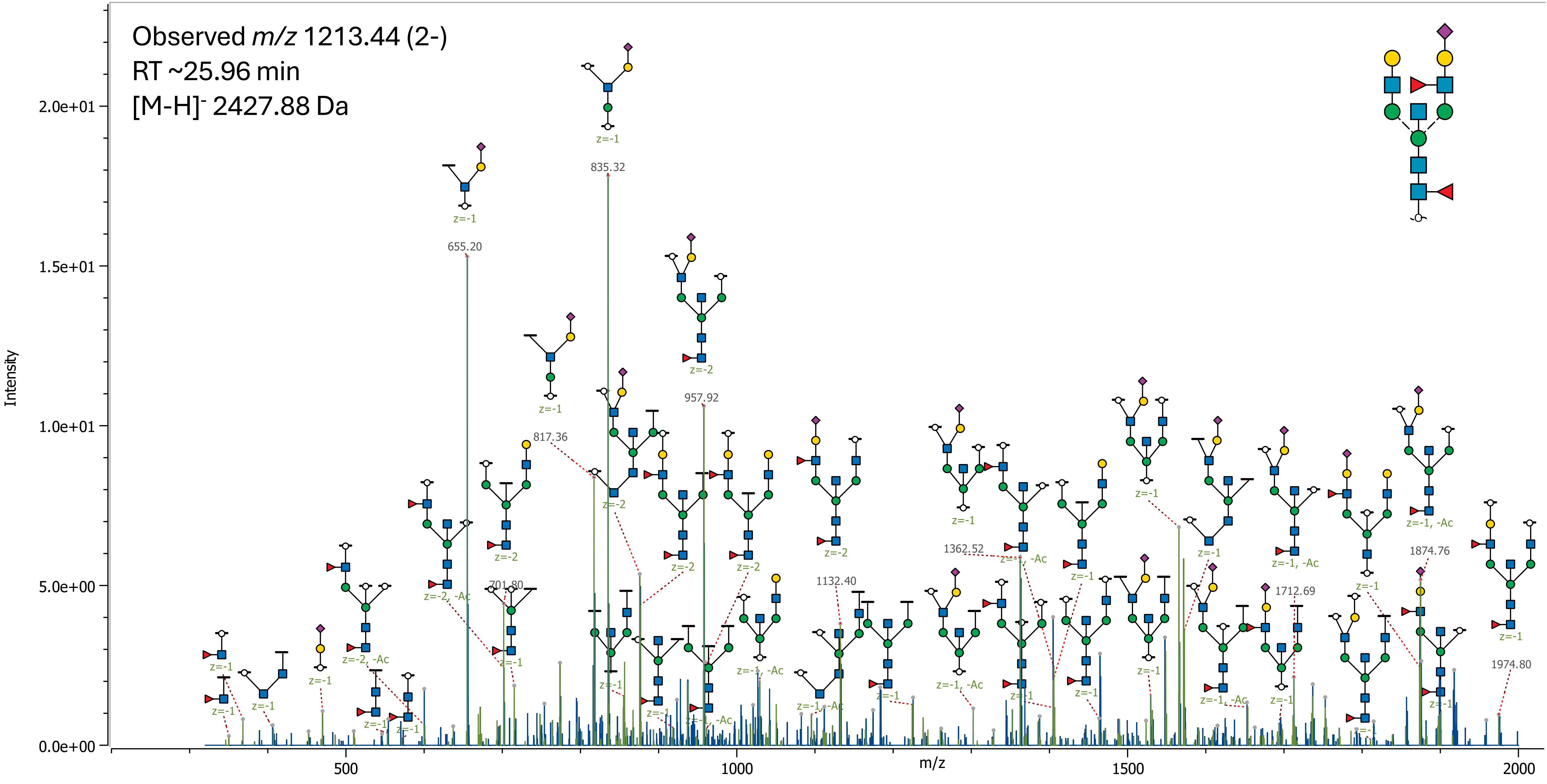

### 66 (Hex)3 (HexNAc)3 (Deoxyhexose)1 (NeuAc)1 + (Man)3(GlcNAc)2

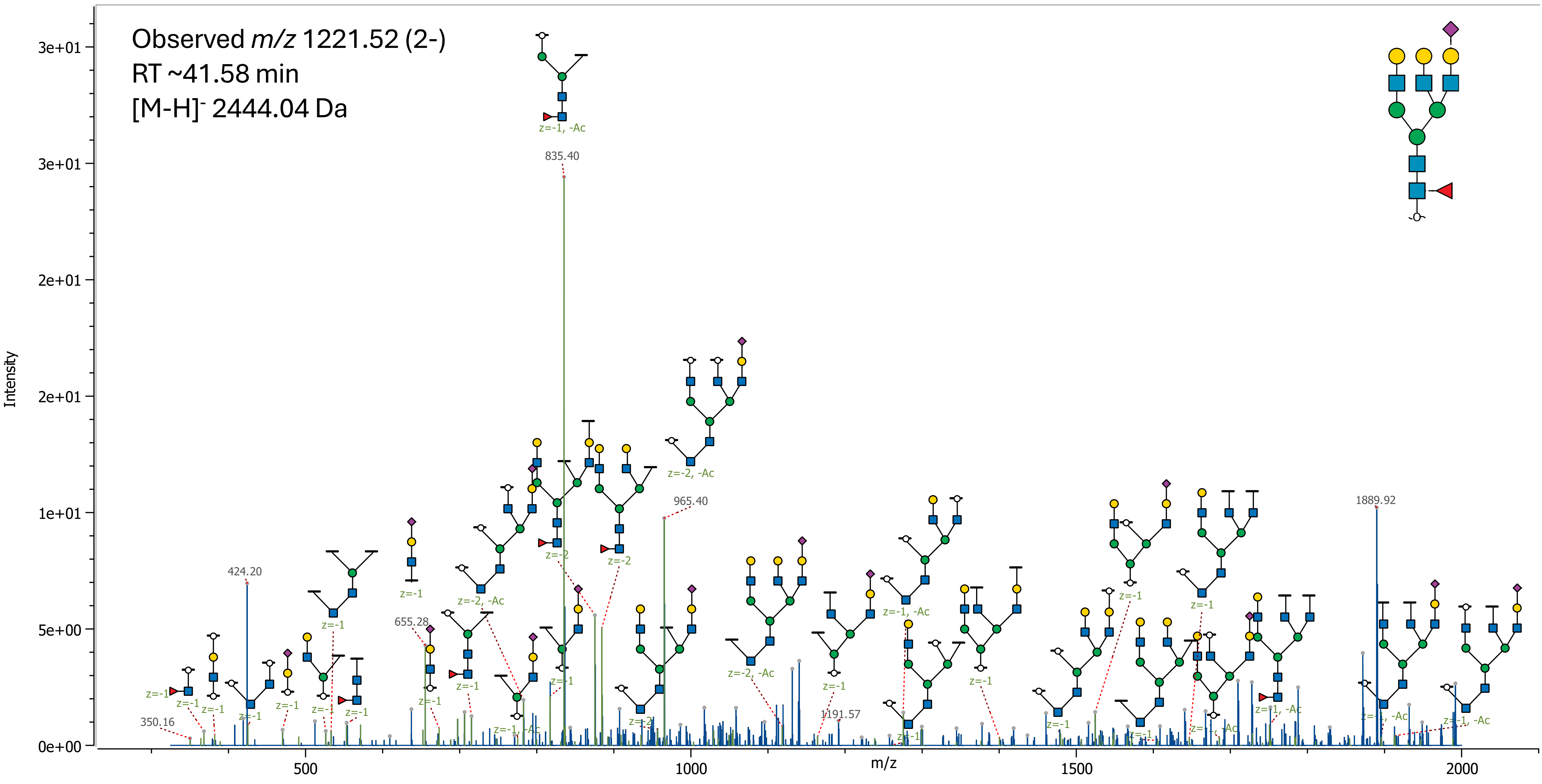

67 (Hex)2 (HexNAc)6 + (Man)3(GlcNAc)2

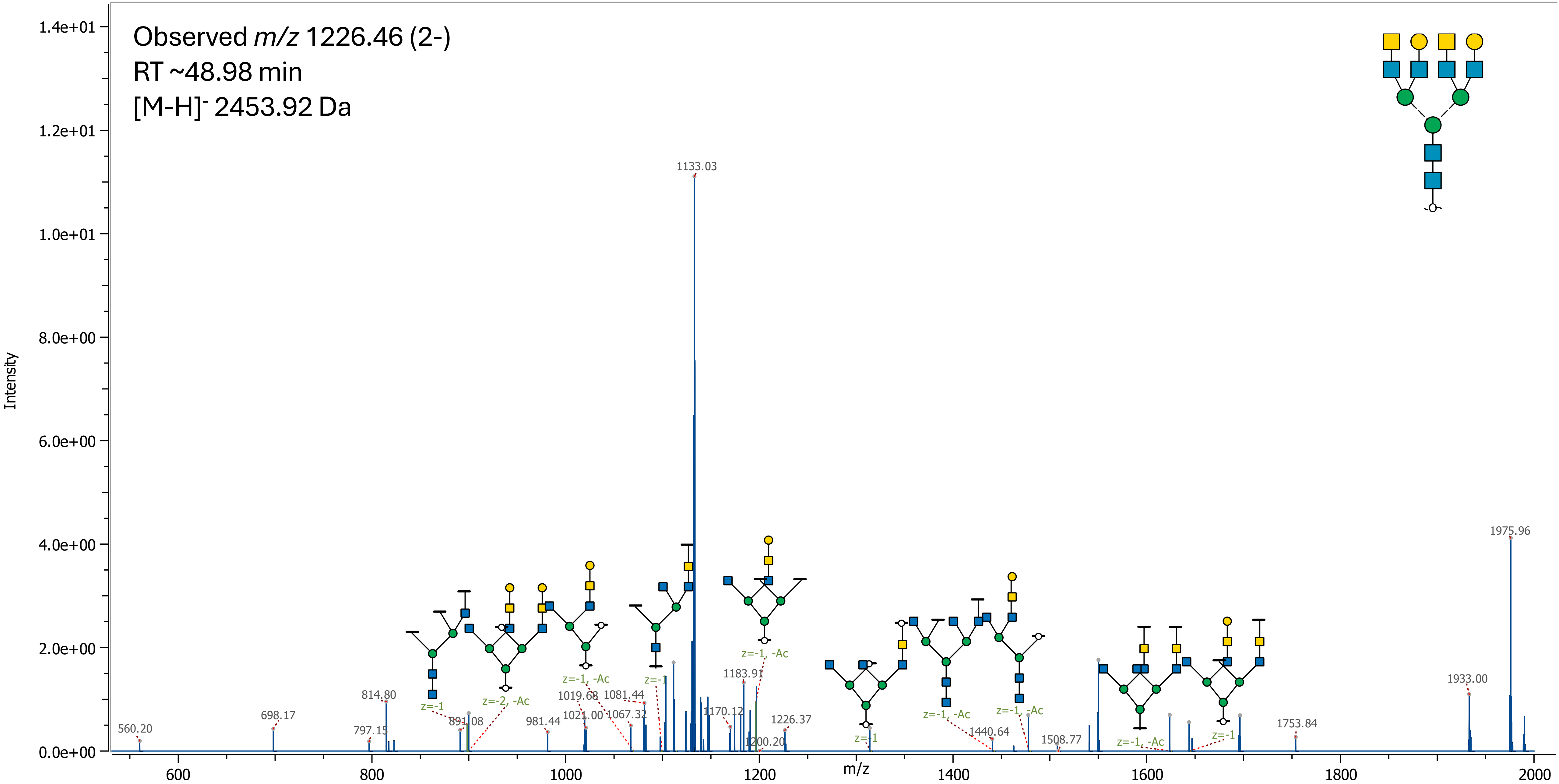

68 (Hex)2 (HexNAc)3 (Deoxyhexose)1 (NeuAc)2 + (Man)3(GlcNAc)2

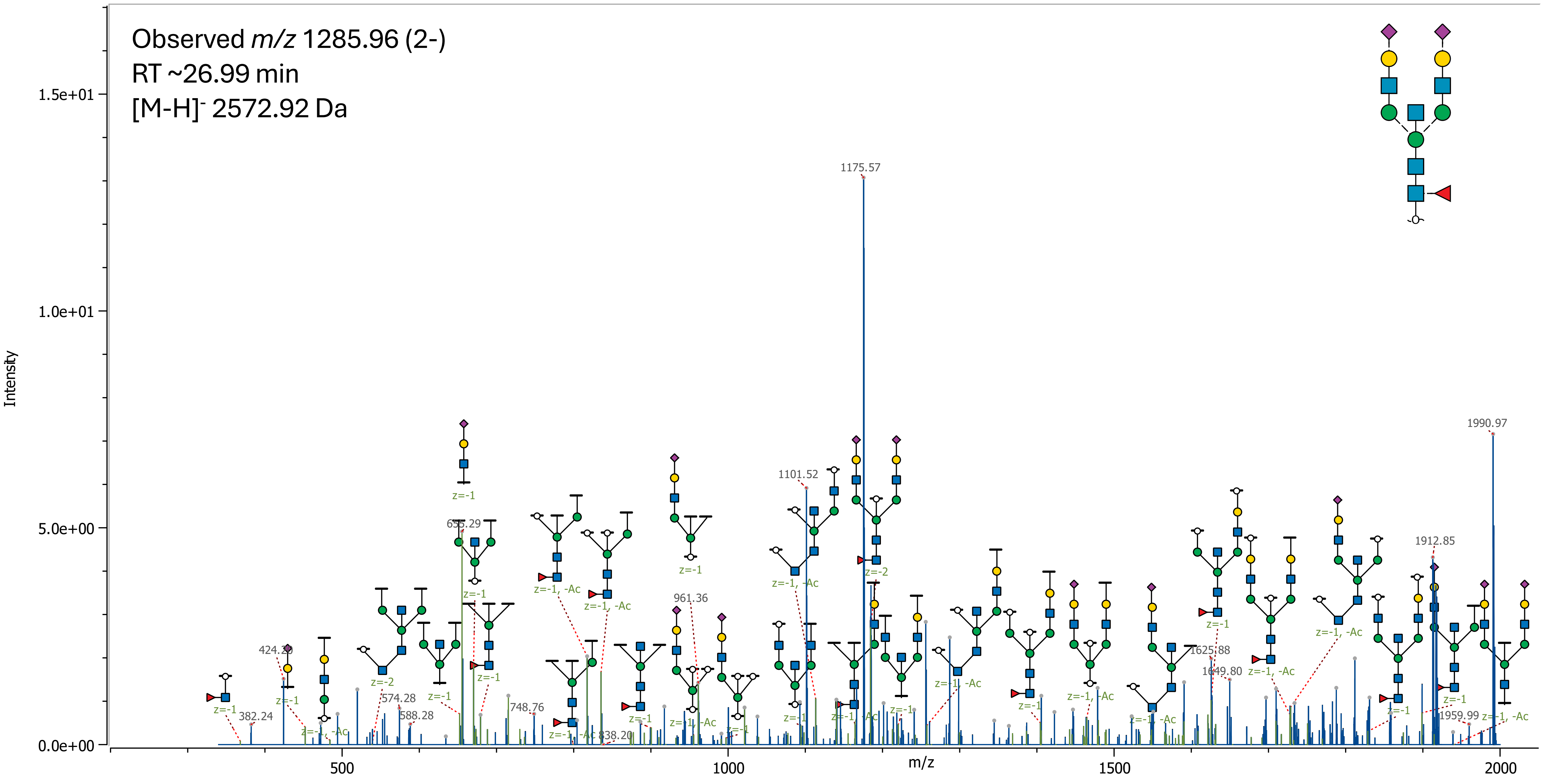

69 (Hex)3 (HexNAc)3 (NeuAc)2 + (Man)3(GlcNAc)2

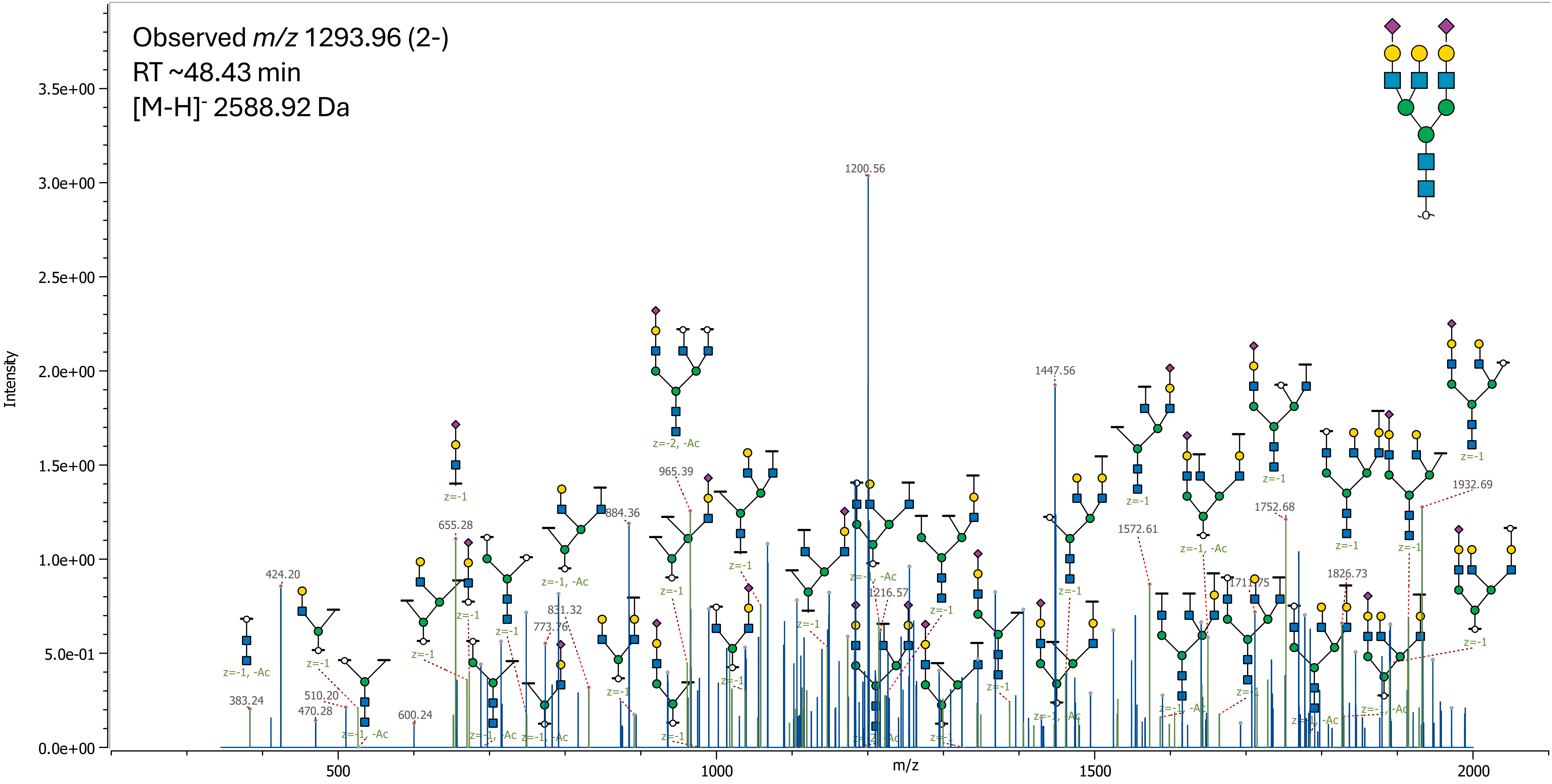

### 70 (Hex)3 (HexNAc)3 (Deoxyhexose)1 (NeuAc)2 + (Man)3(GlcNAc)2

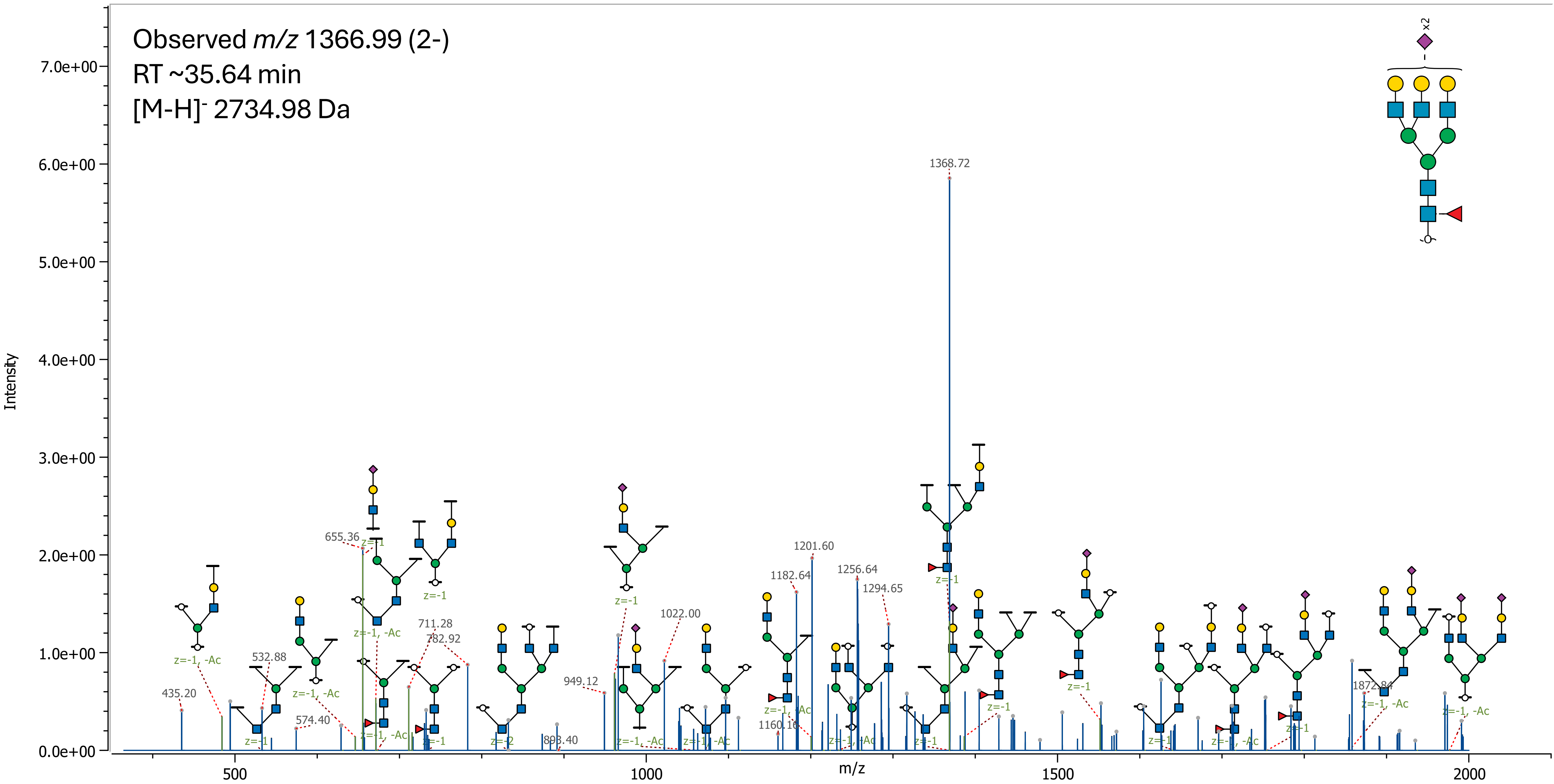

71 (Hex)3 (HexNAc)3 (NeuAc)3 + (Man)3(GlcNAc)2

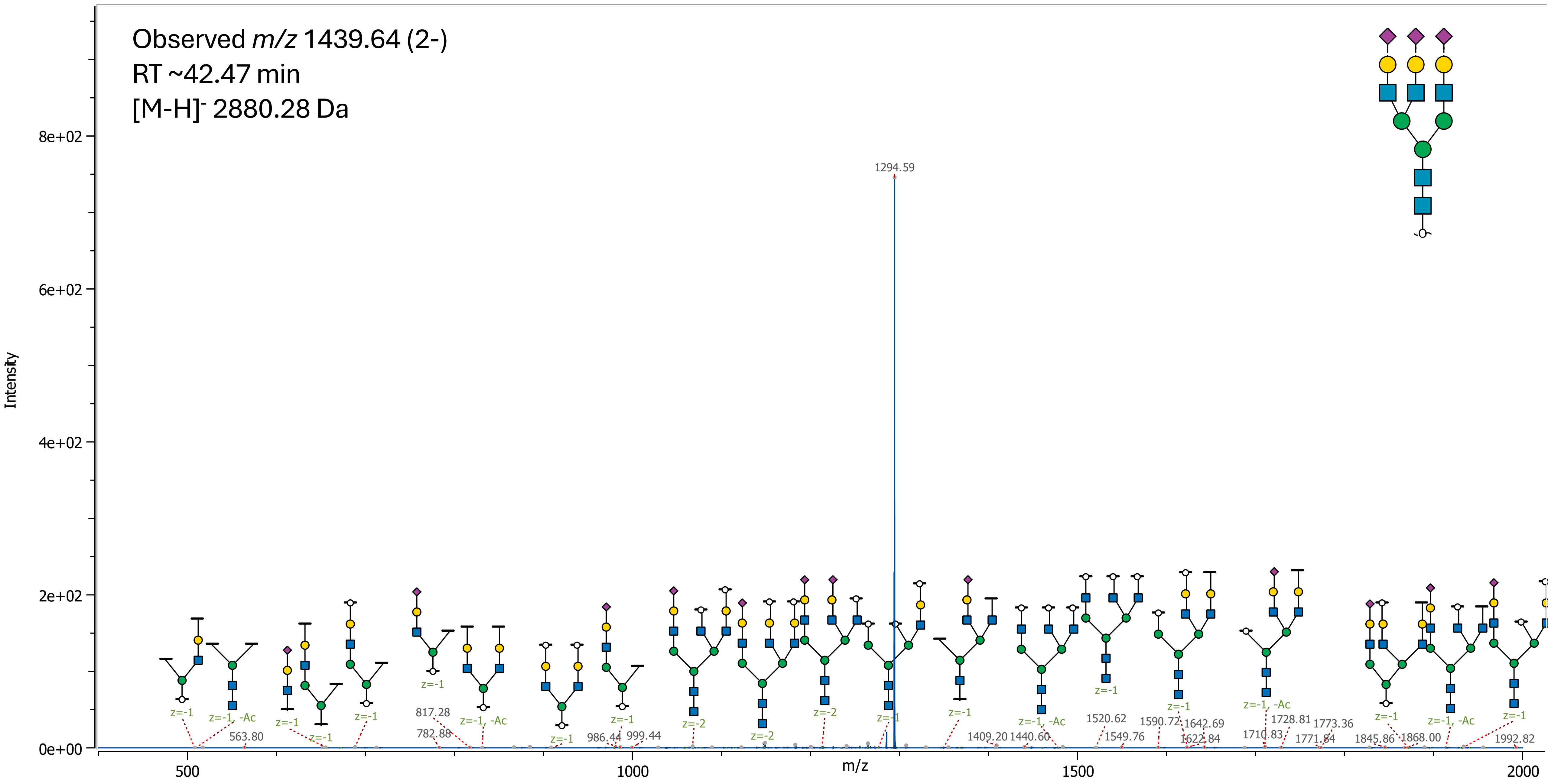

72 (Hex)3 (HexNAc)3 (NeuAc)4 + (Man)3(GlcNAc)2

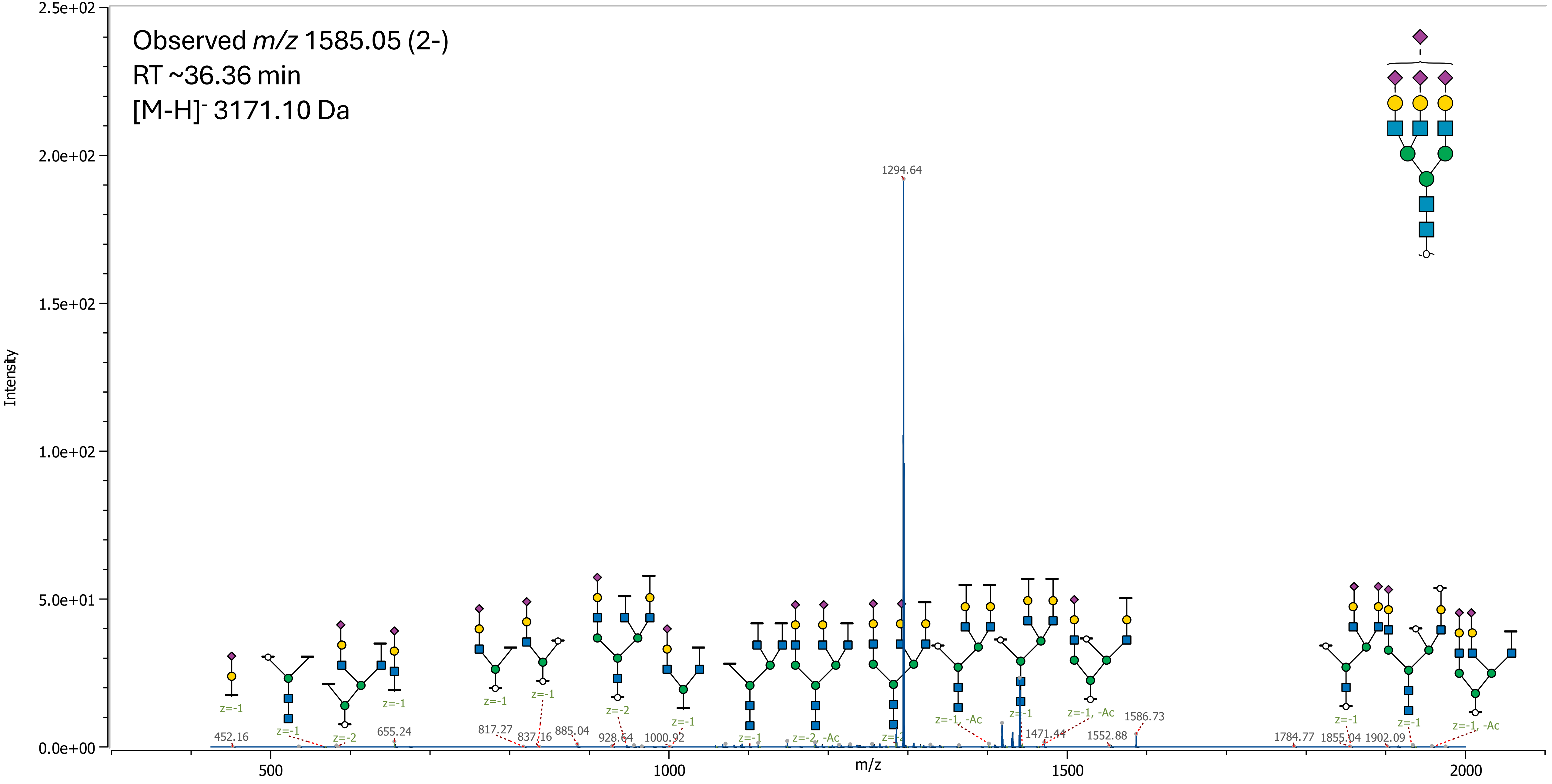

73 (Hex)4 (HexNAc)4 (Deoxyhexose)1 (NeuAc)4 + (Man)3(GlcNAc)2

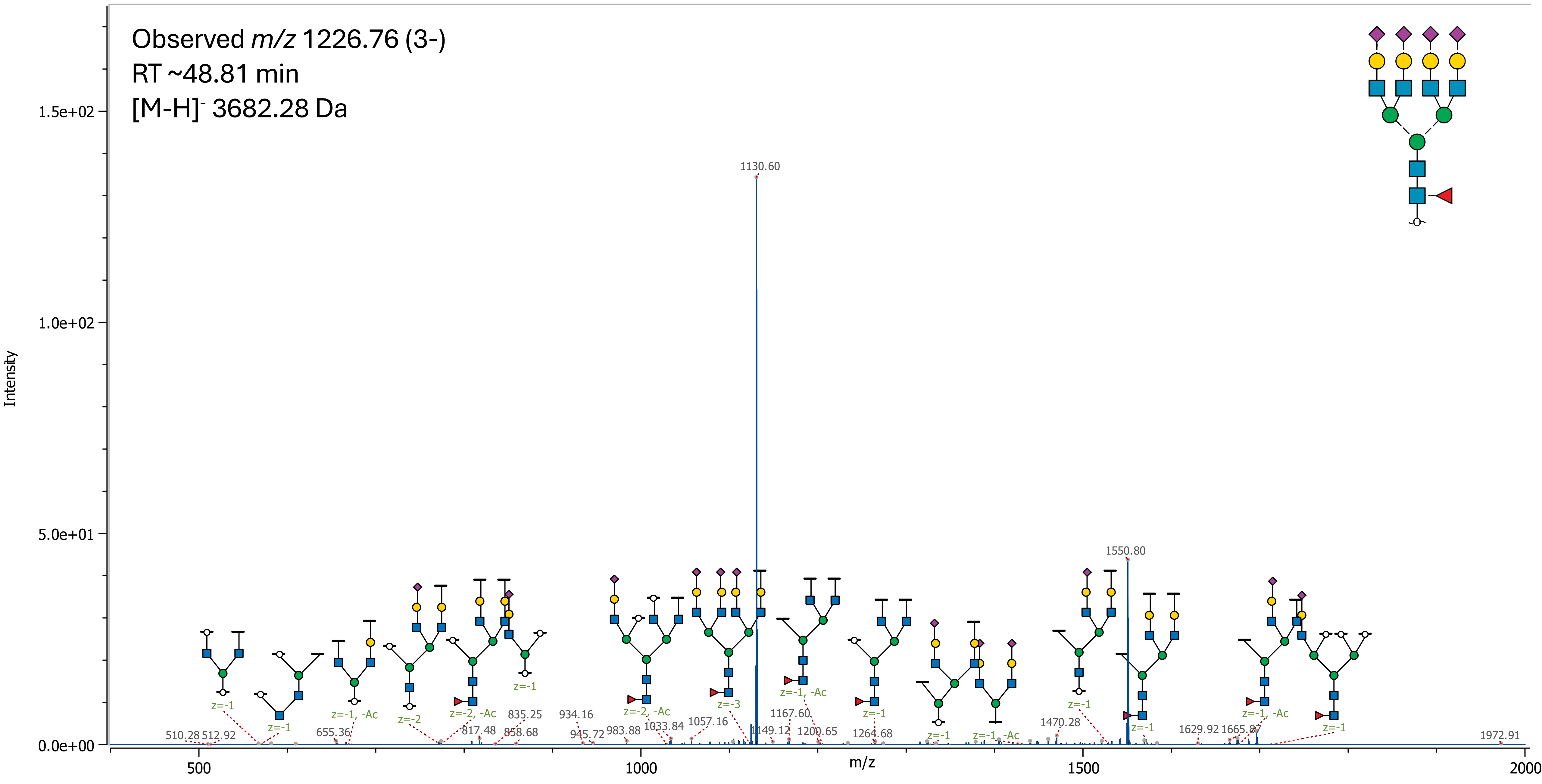

74 (Hex)4 (HexNAc)2 (Deoxyhexose)1 + (Man)3(GlcNAc)2

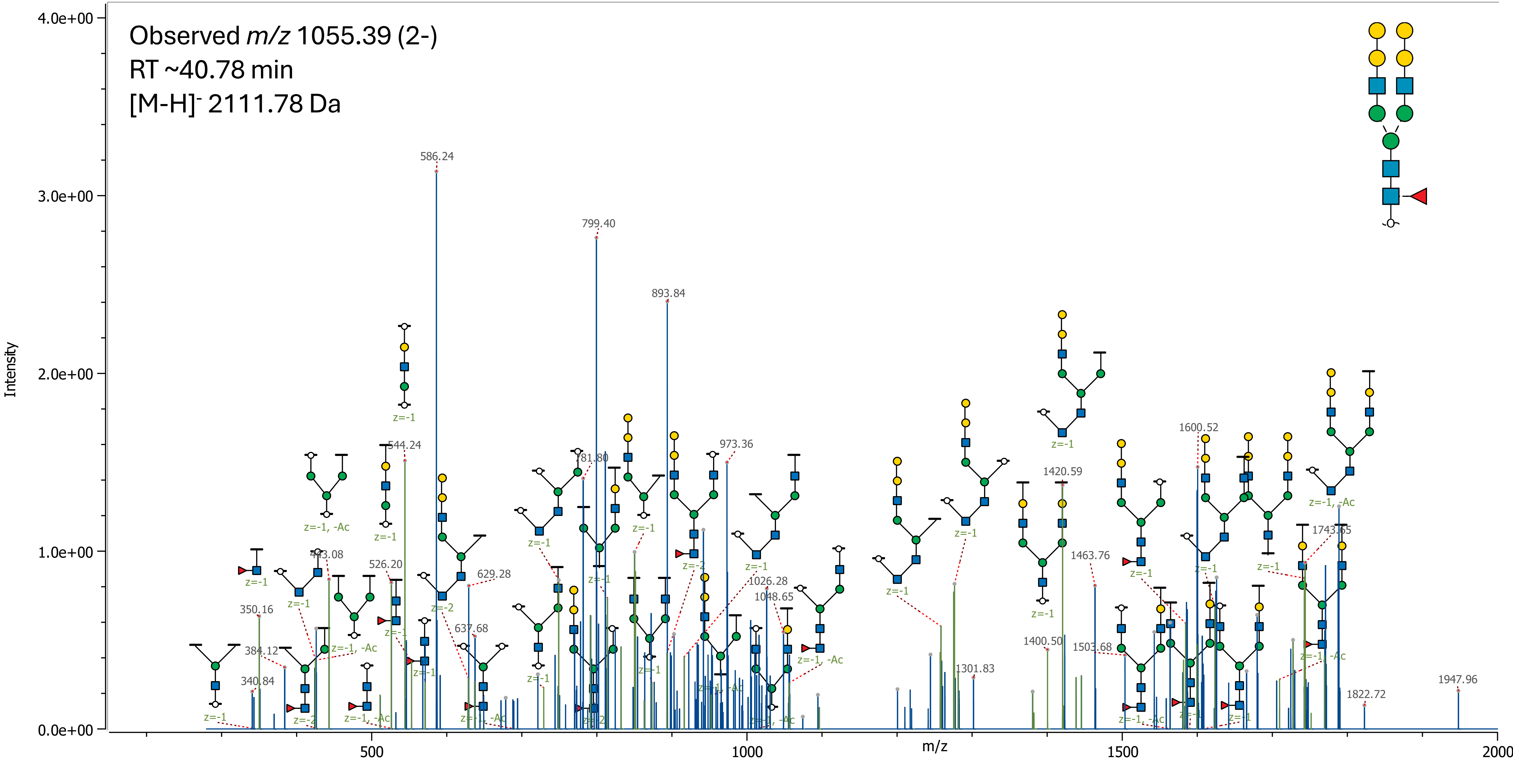

75 (Hex)2 (HexNAc)3 (Deoxyhexose)2 + (Man)3(GlcNAc)2

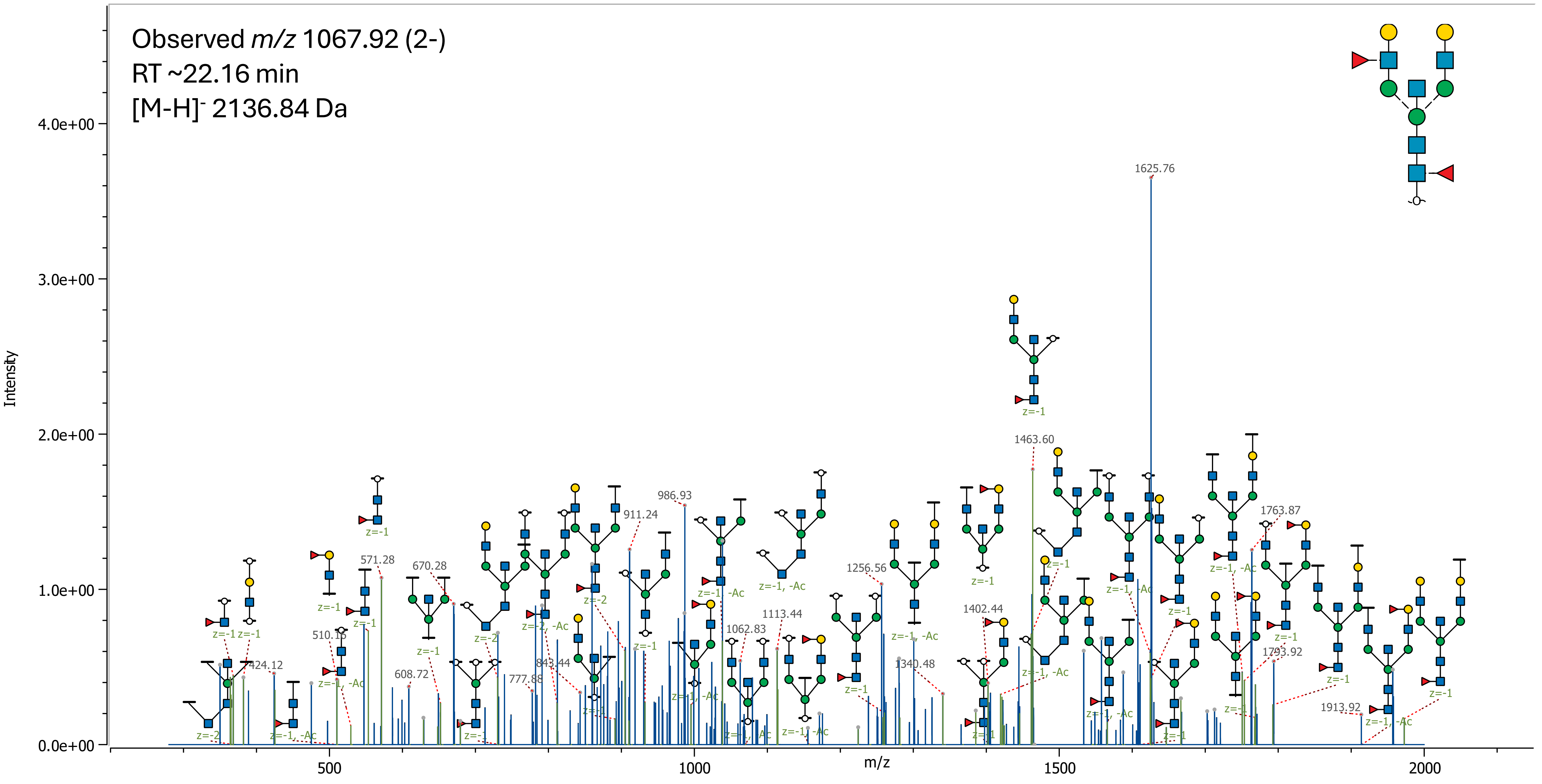

### 76 (Hex)2 (HexNAc)2 (NeuAc)3 + (Man)3(GlcNAc)2

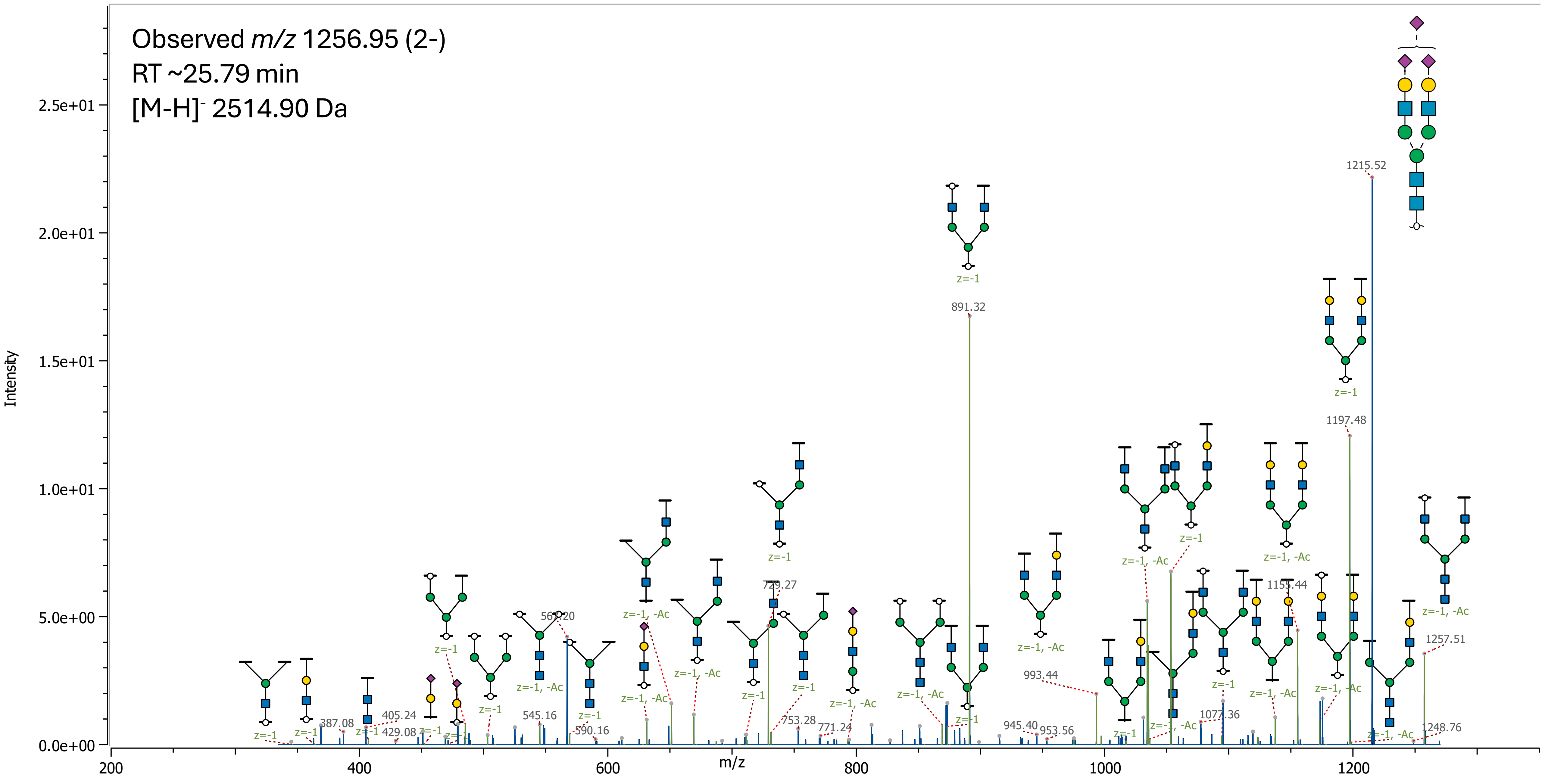
